# Phloem evolved gradually and asynchronously to xylem in early vascular plants

**DOI:** 10.64898/2026.03.23.713640

**Authors:** Laura M. Cooper, Alexander J. Hetherington

## Abstract

The evolution of the water-conducting xylem and sugar-conducting phloem tissues were key innovations in land plant evolution, enabling the origin of long-distance transport networks^1^. In extant vascular plants, phloem and xylem are linked functionally and always occur together^2^, though their evolutionary origin is unclear. This uncertainty is owed to the greater fossilisation potential of lignified xylem tracheids compared to thin-walled phloem cells^3, 4, 5^. Therefore, the fossil record of xylem is far more extensive than that of phloem, with the first definitive record of xylem being around 40 million years earlier^6^ than phloem^7^. This bias in the fossil record obscures characterisation of the origins of plant vasculature. In this study, this limitation is overcome by re-describing the “phloem-like” tissues of exceptionally preserved plants from the 407-million-year-old Rhynie chert^8-15^. We report that this tissue differs markedly from the phloem of extant plants, and propose its identification as a tissue of food-conducting cells (FCCs). Major histological differences were observed in the fossil plants, including no evidence for a pericycle, which in extant species delimits vascular from ground tissues, and the FCCs of the Rhynie chert plants were significantly larger in diameter than phloem cells. These differences suggest that early vascular plants lacked true phloem. However, putative sieve pores in the FCCs of *Asteroxylon mackiei* were identified. This represents to our knowledge the earliest record of sieve pores in the fossil record. Our results suggest an evolutionary scenario in which phloem features assembled gradually within FCCs, asynchronous to the evolution of xylem.

## Results

The Rhynie chert is a 407-million-year-old fossil site^8^ that preserves a diversity of plant species spanning the origin of the vascular plants^11, 12, 16, 17^, the separation of the lycophytes from the euphyllophytes and diversification within the lycophytes^10, 13, 17, 18, 19^. These species therefore provide an opportunity to investigate the origin and evolution of xylem and phloem at the base of the vascular plant clade. Previous work on the water-conducting tissues of the Rhynie chert plants has documented a clear trajectory from polysporangiophytes with either no water-conducting cells^20^ or water-conducting cells distinct from xylem^16^, to plants with xylem^11, 12, 13, 14, 17, 21.^ In contrast, the occurrence of phloem in the Rhynie chert land plants is highly debated. Phloem is reported in early descriptions of the Rhynie chert land plants, owing to the position of this tissue, its cellular organisation^11, 12, 13, 14^, and putative subcellular anatomy^22^ (critiqued in^23^). It has been subsequently debated if this tissue should be considered true phloem^9, 10, 16, 23, 24, 25^. Resolving the identity of the “phloem-like” tissue of the Rhynie chert plants is therefore imperative for understanding the origins and evolution of vascular tissues.

To determine if phloem was present in the Rhynie chert plants, the sporophytes of four species were investigated: the non-vascular polysporangiophyte *Aglaophyton majus* (with a central strand of water-conducting cells (WCCs)) ^12, 16^, and three species of vascular plants with true xylem *Rhynia gwynne-vaughanii*^11, 12^, and two lycophyte species, the zosterophyll *Trichopherophyton teuchansii*^18^ and the lycopsid *Asteroxylon mackiei*^13^. Histological and cellular comparisons were then made between these fossils and four species of extant lycophytes. This investigation will refer to the tissue of the Rhynie chert plants as food-conducting cells (FCCs), following recent literature^25^, and the term “phloem” will only apply to tissue exhibiting the features of phloem as it is defined in extant vascular plants. In extant vascular plants phloem is histologically distinguished from surrounding tissues by a pericycle, at the cellular level by elongated cells with very low aspect ratios, and at the subcellular level by sieve pores in the walls of sieve elements^2^. Comparison of FCCs of the Rhynie chert plants and the phloem of extant lycophytes were made on three major levels of organisation that define primary phloem in vascular plants: histological, cellular and subcellular.

### The Rhynie chert plants lack histological distinction between vascular and ground tissues

In extant vascular plants, phloem is identified histologically by its position surrounding or peripheral to xylem, and these tissues together comprise the vascular strand^2^. The vascular strand is delimited from the ground tissue by a bounding layer, the pericycle^2, 26^, and in roots also an endodermis^27^ or endodermoid^28^. The pericycle is classically defined as an irregular layer of one or more cells thick between the vasculature and the cortex in stems, and between the vasculature and endodermis in roots^29, 30, 31^. The pericycle is a widespread feature across vascular plants, though it is absent in the stem of some derived groups^2, 31^. In the anatomical literature on extant lycophytes, the pericycle is often distinguished in figures (e.g.,^2, 26, 27^), though often lacks a precise definition (e.g.,^26, 27, 32^). In this study, the pericycle was identified in the extant lycophytes as a layer of one to three cells thick forming a distinct boundary between the phloem and cortex. The cells of the pericycle layer are larger compared to the adjacent phloem, but lack airspaces seen in the cortex (**Fig. 1J**). Given the widespread occurrence of an anatomically distinct pericycle in extant vascular plant species^2, 26. 27, 31.32^, evidence for a comparable layer was examined in the Rhynie chert plants.

**Figure 1.**
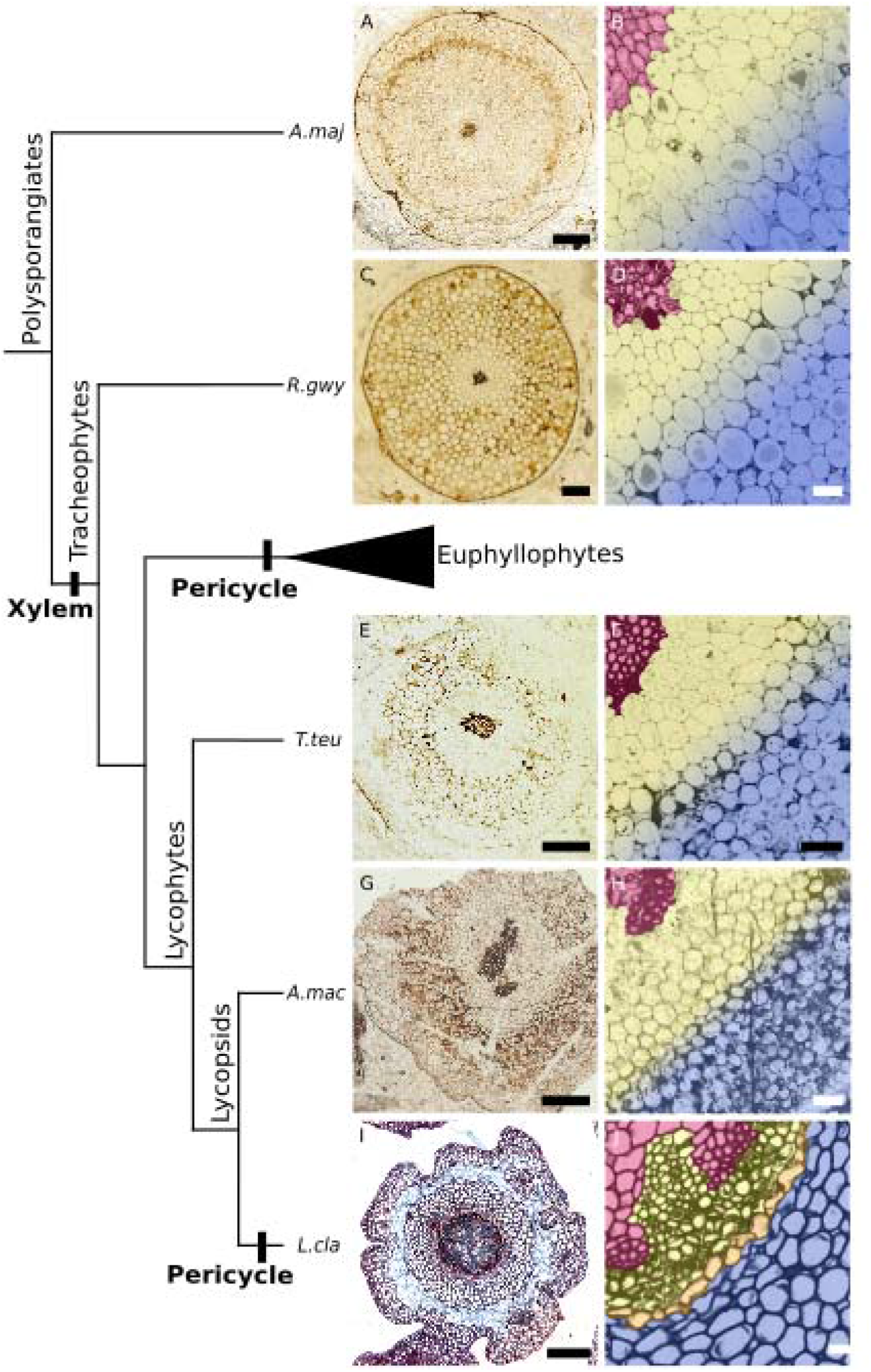
The food conducting cells (FCCs) of the Rhynie chert plants grade into the cortex tissue, and there is no pericycle between the FCCs and the cortex. (A-H) The aerial axes of the Rhynie chert plants show a central strand of water-conducting tissue (water conducting cells in *Aglaophyton majus* and xylem in the rest, shown in magenta false colouring) surrounded by an elongated, close-packing tissue (yellow), referred to as the food conducting cells (FCCs), followed by cortex (blue). (I-J) In extant lycophytes and euphyllophytes, the phloem tissue (yellow) is separated from the cortex (blue) by a pericycle (orange). A clear pericycle is lacking in Rhynie species (A-H) and instead the FCCs grade into the cortex (show by a fuzzy boundary between FCCs and cortex on B, D, F and H). *Species abbreviations: A. maj = Aglaophyton majus; R. gwy = Rhynia gwynne-vaughanii; T. teu = Trichopherophyton teuchansii; A. mac = Asteroxylon mackiei; L. cla = Lycopodium clavatum*. Phylogeny based on^17^, positions of *Aglaophyton majus* and *Rhynia gwynne-vaughanii* from^68^. Specimen codes: A-B = MPEG0015, C = SCOTT EUCM 1361, D = MAN EMu 547126, E-F = LYON 93.11, G-H = GLAHM Kid 2472 Scale bars: 500 µm in (A, E, G), 200µm in (C, I), 100µm in (F, H); 50µm in (B, D); 20µm in (J).

The FCCs and cortex cells were readily distinguished in all four of the Rhynie chert species examined. The FCCs comprise a tissue of close-packing, elongated cells which surround the central core of xylem or WCCs^9, 10, 11, 12, 13^. By contrast the cortex is characterised by shorter, rounder cells with many intracellular spaces (**Fig. S1 A-C**). These tissues have similar characteristics in the extant lycophytes. However, whereas there is clear demarcation between the tissues by the pericycle in the extant species (**Fig. 1 I, J**), a comparable layer is absent in the Rhynie chert plants. Instead, the FCCs grade out into the cortex, and this is especially pronounced in *A. majus* and *R. gwynne-vaughanii*. For these two species, the grading of the FCCs into the cortex can be seen in transverse section. Immediately adjacent to the xylem/WCC strand there is a tissue of close-packing cells, and when moving peripherally, the intracellular spaces that characterise the cortex appear initially small and infrequent, increasing in frequency and size towards the axis periphery, with no sharp transition between these tissues (**Fig. 1 A-D**). In longitudinal section, the tissue adjacent to the xylem/ WCC strand is of close-packing and elongated cells, and when moving peripherally the cells appear increasingly shorter, and with more intracellular spaces, until a tissue of the typical rounded cortex cells with many large intracellular spaces is reached (**Fig. S1**). This pattern is demonstrated quantitatively in the *R. gwynne-vaughanii* axis shown in (**Fig. S2**), where the cells immediately adjacent to the xylem strand (the FCCs) are elongated, narrow and close-packing, with an average aspect ratio (AR) of 0.20 (range 0.31-0.10). Peripheral to this is a region of looser-packing but elongated cells, referred to here as transitional tissue (at least in part corresponding to the inner cortex in e.g., ^9^). These transitional cells have an average AR of 0.30 (range 0.58-0.14), indicating that they are also elongated, though on average shorter and wider than the FCCs (**Supplementary Table 1**). Peripheral to the transitional tissue is a classical cortical tissue of shorter, more rounded cells with many intercellular spaces, and an average AR of 0.62 (range 1.22-0.20) (**Fig. S3**). This shows that there is not a sharp transition of cell morphology between the cortex and FCCs, but rather a grading of tissue types, as the identification of the transitional tissue demonstrates.

Investigation of the lycophytes *T. teuchansii* and *A. mackiei* indicate that these species have a sharper transition from FCCs to cortex tissue compared to *A. majus* and *R. gwynne-vaughanii*, which can be seen in both transverse section (**Fig. 1E-H**) and longitudinal section **(Fig. S1D-G)**. Despite this, there is still an absence of a boundary layer in these extinct lycophytes like the pericycle of extant lycophytes. The absence of an endodermal layer in *A. mackiei* rooting axes has previously been proposed^33^, but the observation in this study of the absence of a pericycle in all *A. mackiei* axes is novel.

In conclusion, our histological investigation into the Rhynie chert plants finds a distinct FCC tissue between the xylem/WCC and cortex tissues. However, unlike in extant species where the vasculature and cortex are separated by a clear pericycle layer, the Rhynie chert plants lack a bounding layer, and exhibit a grading of tissue characteristics between the cortex and FCCs. This suggests that FCCs are histologically distinct from phloem.

### Cellular characteristics of the FCCs of the Rhynie chert plants are distinct from those of the phloem of extant lycophytes

At the cellular level, the cells of the phloem, especially the specialised sugar-conducting sieve elements, are narrow, close-packing and elongated^34^. Overall, excluding some outlying angiosperm groups, extant vascular plant sieve elements tend to be around 10-20 μm in diameter, and over 500 μm in length^35, 36^, producing an aspect ratio (AR) of 0.02 or less. In light of this, the cellular characteristics of the FCCs of the Rhynie chert plants were examined and compared to those of the phloem of extant lycophytes.

The phloem cells of the extant lycophytes (sieve elements and phloem parenchyma) measure between 41.4 μm and 2.2 μm in diameter, with an average diameter of 8.9 μm across species and tissue types (n = 9947, 25 axes). In contrast, the FCCs of the Rhynie chert plants measure between 231.3 μm and 9.2 μm in diameter, with an average diameter of 56.2 μm across species and tissue types (n = 2907, 21 axes). This indicates there is a significant difference between the FCC and phloem cell diameters (p ≤ 0.001), with the FCCs of the Rhynie chert plants being on average around six times larger in diameter than the phloem cells of extant lycophytes (**Figure 2**). This difference in cell size between the Rhynie chert plants and the extant lycophytes remains significant (p ≤ 0.001) when the larger axis diameter of the Rhynie chert plants is taken into account (**Fig. S4**), as well as the overall larger cell size of the Rhynie chert plants (**Fig. S5**, p ≤ 0.001) (see **Supplementary Data Table and Supplementary Data Images**). This investigation of cellular structure of the Rhynie chert plant FCCs suggests that these cells are markedly different in diameter from the phloem cells in extant lycophytes, and that this is independent of other morphological and anatomical size differences.

**Figure 2.**
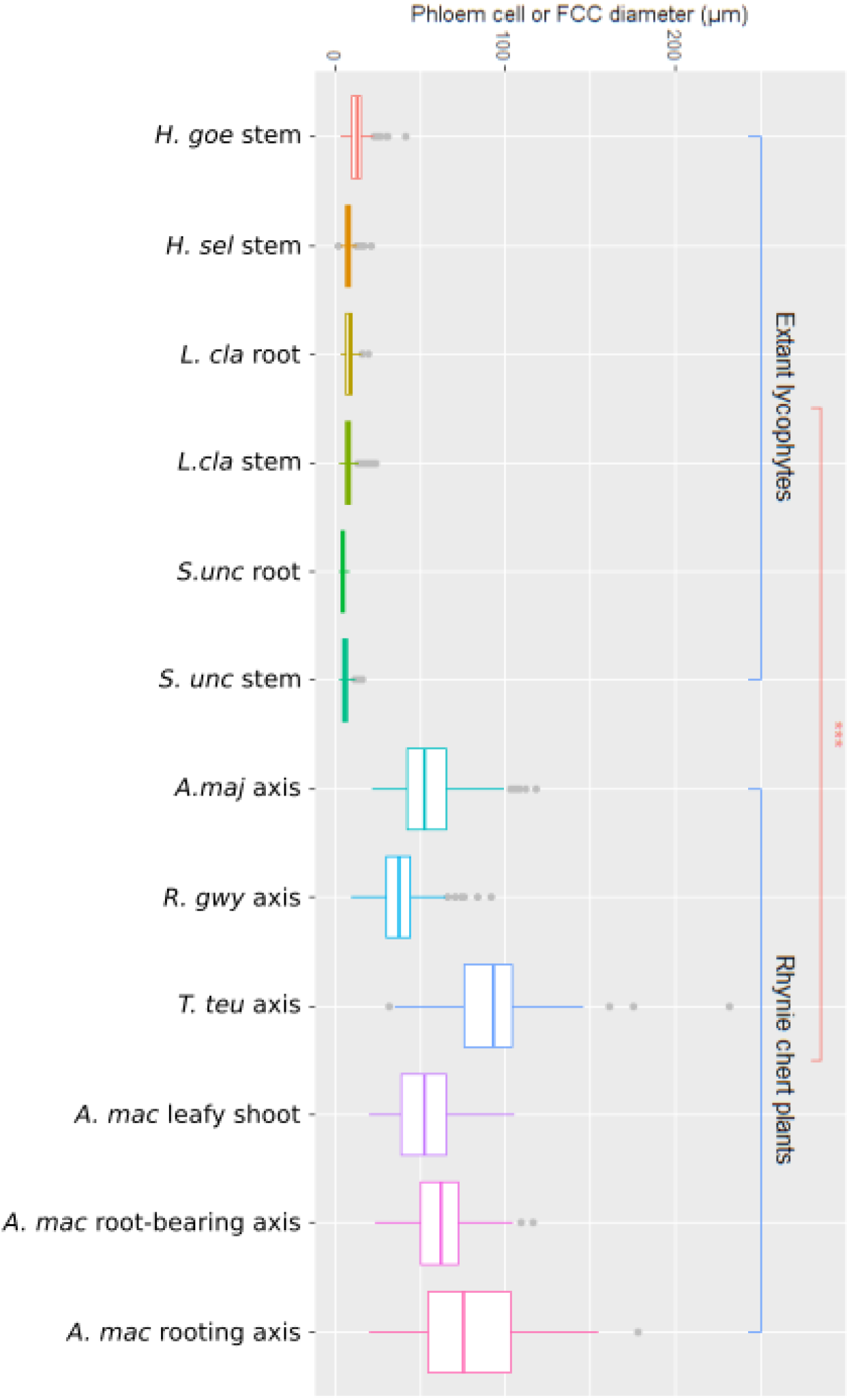
The FCCs of the Rhynie chert plants are significantly (p≤0.001) larger in diameter than the phloem cells of extant lycophytes. Measurements of the diameter of phloem or food conducting cells in extant lycophytes compared to extinct species from the Rhynie chert. Four extant lycophyte species (*Huperzia goebellii, Huperzia selago, Lycopodium clavatum, Selaginella uncinata*) were investigated covering both stem and root tissues. Four Rhynie chert species covering axis tissues (*Aglaophyton majus, Rhynia gwynne-vaughanii, Trichopherophyton teuchansii*) and leafy stem, root bearing axis, and rooting axis (*Asteroxylon mackiei*). The FCCs of the Rhynie chert species are all significantly (p≤0.001) larger than the phloem cells of the extant lycophytes, as demonstrated by a t-test.

### Discovery of putative sieve pores in A. *mackiei*

The final way that the phloem of extant vascular plants is defined is at the subcellular scale. The walls of phloem sieve elements are perforated by sieve pores, which form from the enlargement of plasmodesmatal pits, and enable rapid long-distance symplastic transport through the phloem^37^. Sieve pores are incredibly rare in the fossil record, but have been preserved in fossils with exceptional preservation, demonstrating that they can be fossilised^38 – 52^. The presence of sieve pores is often taken as the defining feature of phloem in the fossil record (e.g.,^7^), owing to their distinctness compared to the cell walls of other parenchymatous tissues, with only plasmodesmatal pits. Phloem is often described from derived extinct lineages in the absence of sieve pores (e.g., ^53^), and there are tentative descriptions of phloem from older deposits and earlier-diverging groups, but which lack the preservation quality to preserve sieve pores (e.g.,^54^). Owing to the importance of these structures, the occurrence of sieve pores in the FCCs of the Rhynie chert plants was investigated. Firstly, to obtain comparative images of extant lycophyte sieve pores and plasmodesmatal pits, the extant lycophytes *Lycopodium clavatum* and *Huperzia selago* were imaged using SEM (**Fig. 3A** and **B and Fig. S6)**. Sieve pores in the lateral walls of *Lycopodium clavatum* SEs were found to be 0.15 µm in diameter on average (**Fig. 3B**), in the ranges previously reported (0.2 to 0.86 µm^36^, ^55, 56^). Plasmodesmatal pits in the lateral walls of phloem parenchyma in *Huperzia selago* were found to be 0.07 µm (70 nm) in diameter on average (**Fig. S6**), slightly larger than reported for pits in lycophyte shoot apical meristems (26 to 41 nm^57^). This shows that the sieve pores of lycophyte sieve elements are at least twice as large in diameter as, and can be distinguished from, the plasmodesmata pits of phloem parenchyma. Using these results as a search image, *A. mackiei* FCCs were investigated for the occurrence of sieve pores.

**Figure 3.**
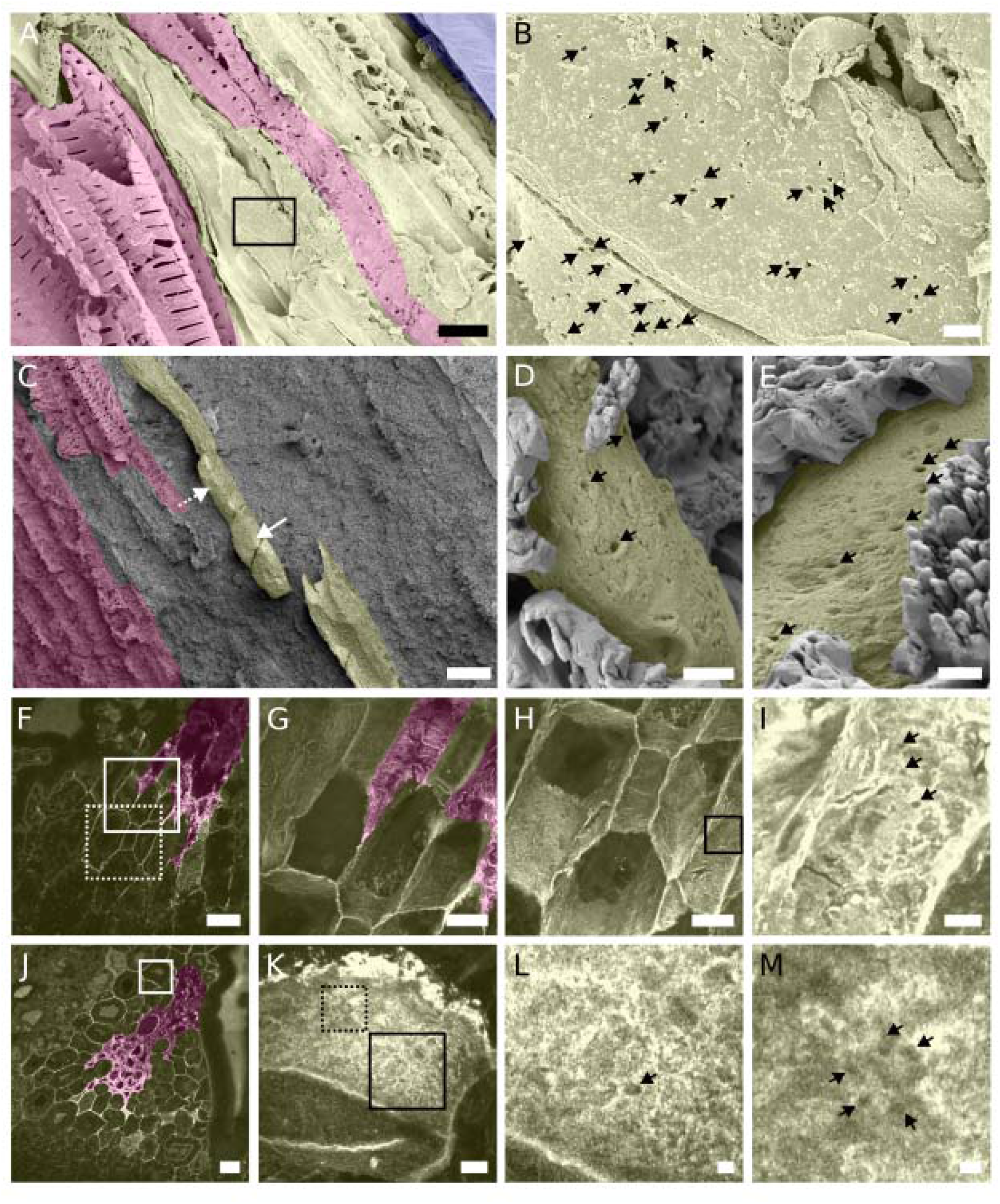
Discovery of putative sieve pores in *Asteroxylon mackiei* like the sieve pores of extant lycophytes. (A, B) SEM imaging of the lateral walls of *Lycopodium clavatum* sieve elements (yellow false colouring) shows numerous sieve pores (black arrows in B). The region within the box on A is shown in B. (C-E) SEM imaging of the root-bearing axis of *Asteroxylon mackiei* shows lycophyte-sized pores in the lateral wall of FCCs; C, HF etching exposes vascular tissue (magenta = xylem, yellow = FCCs), whilst much of the FCC indicated in C is surrounded by silica, parts of the external surfaces of the lateral cell walls are exposed (arrows); D shows the exposed cell wall in the region indicated by the dashed white arrow in C, and E shows the exposed cell wall in the region indicated by the solid white arrow in C, in both sieve pore-sized structures are indicated by black arrows. (F-M) CLSM and Airyscan CLSM imaging of rooting axes of *Asteroxylon mackiei* shows lycophyte-sized pores in the end and lateral wall of FCCs. (F-I), F shows a single plane CLSM image of an *Asteroxylon mackiei* rooting axis in oblique section; G maximum intensity projection of the region in the solid white box in F, and H shows the region in the dashed white box in f imaged using the same method; I shows a maximum intensity projection of the region in the solid black box in H, sieve pore-sized structures in the lateral cell wall are indicated by black arrows. (J-M), J shows a single plane CLSM image of an *Asteroxylon mackiei* rooting axis in cross section obtained using an x40 objective; K shows a maximum intensity projection of an Airyscan CLSM z-stack image obtained using a x100 objective of the region in the solid white box in J; l and M both show details of K showing, respectively, the region in the dashed and in the solid black box in K, sieve pore-sized structures in the end wall are indicated by black arrows. False colour overlays: magenta = xylem; yellow = phloem/ FCCs; blue = cortex. Scale bars = 50µm in (C, F, J); 20µm in (G, H); 10µm in (A); 5µm in (I, K); 1µm in (B, D, E, L, M). Specimen codes = F-I = MPEG0062, J-M = MPEG0043

*A. mackiei* alone was focused on owing to the intensive imaging required to detect sieve pores, and as this species is a lycopsid, it is the most likely to have traits in common with extant lycophytes. Samples of *A. mackiei* were subjected to Scanning Electron Microscopy (SEM) and Airyscan Confocal Laser Scanning Microscopy (CLSM) imaging to ascertain if sieve pores were found in the walls of the FCCs. Airyscan CLSM and SEM were used together as these techniques are complimentary. Whilst SEM has a resolution limit (<1 nm^58^) that allows subcellular structures to be clearly imaged, SEM preparation of Rhynie chert samples requires hydrofluoric acid etching, which could cause the collapse of exposed cells. Airyscan CLSM is a non-destructive technique used on thin sections, but does have a reduced resolving power (the resolution limit of Airyscan CLSM is 0.12 to 0.2 µm for a 100x objective lens with a 1.4 Numerical Aperture^59-61^). The FCCs of *A. mackiei* were subject to SEM imaging to identify exposed walls of FCCs otherwise supported internally with intact silica, therefore preventing cell collapse in sample preparation (see **Methods**). This revealed an FCC from a root-bearing axis partially encased in silica, but with exposed regions of intact external cell wall (**Fig. 3C**). At two exposed regions of this FCC (white arrows in **Fig. 3C**), regular circular perforations of the cell wall were identified (**Fig. 3D** and **E**, black arrows), which measure around 0.2 µm in diameter. This provided the first tentative evidence of sieve pores in *A. mackiei*. Further evidence of sieve pores was sought using Airyscan CLSM. Airyscan CLSM is capable of resolving submicron features in fossil samples^60,62^, and is readily able to image the 1-2 µm diameter xylem pits of *A. mackiei* (**Fig. S7**). Thin sections of *A. mackiei* rooting axes with well-preserved FCCs (**Fig. 3F-H** and **J-K**) were imaged using Airyscan CLSM with a x100 objective. **Fig. 3I** and **L - M**, showing regular circular perforations of lateral and end walls, respectively, which are <0.5 µm in diameter. Therefore, under both SEM (**Fig. 3C-E**) and Airyscan CLSM (**Fig. 3F-I** and **J-M**), the FCCs of *A. mackiei* were found to have perforations in the lateral and end walls similar in diameter to the sieve pores of extant lycophytes and larger than those of plasmodesmata pits, and are therefore identified as sieve pores. This represents to our knowledge the earliest record of sieve pores in the fossil record.

## Discussion

### The Rhynie chert plants exhibit a food-conducting tissue distinct from the phloem of extant vascular plants

The results presented here show that the Rhynie chert plants exhibit a tissue in the anatomical position of phloem tissue and with some adaptions for long distance transport of sugars, namely elongated, close-packing cells and, in the case of *A. mackiei*, sieve pores. However, the FCCs of the Rhynie chert plants lack other features characteristic of phloem, namely there is no pericycle separating FCCs from the cortex, and the FCCs grade into the cortex. The FCCs are also significantly larger in diameter than phloem cells of extant lycophytes, and are shorter in length than the typical SEs of lycophytes, ferns and gymnosperms. This suggests that the FCCs of the Rhynie chert plants are histologically and cellularly distinct from the phloem of the extant vascular plants, though they do share subcellular features, the sieve pores, implicated in adaptation to a sugar conducting function.

### The FCCs of the Rhynie chert plants reveal the asynchronous and independent evolutionary histories of phloem and xylem

The identification of a tissue bearing some, but not all, of the key characteristics of the phloem in the Rhynie chert plants examined here suggests a scenario for the evolution of phloem tissue in which FCCs in polysporangiophytes are homologous, phloem is therefore a derived form of FCC and evolved gradually, and this occurred asynchronously relative to the origin and evolution of xylem (**Fig. 4**). In this scenario, plasmodesmata were present in the ancestor of all embryophytes^63^. Following divergence of the polysporangiophytes, food-conducting tissue evolved in the sporophyte generation, before water-conducting tissues. Evidence supporting this comes from the early-diverging polysporangiophytes the eophytes^25^ and *Horneophyton lignieri* ^20^: in both taxa there is a central strand of elongated cells described as FCCs or transfer cells, respectively, and the FCC of eophytes exhibit pits. These pits from the images in Edwards *et al* (2022) are estimated here as being∼0.22 µm in diameter (**see Supplementary Note**). This is larger than lycophyte plasmodesmata pits but in the range of the sieve pores of extant lycophytes and *A. mackiei* as reported here. From this, we conclude that elongated FCCs with sieve pores were present early in the polysporangiophytes, before the origin of water-conducting tissues. The presence in the non-vascular *A. majus*, and the tracheophytes *R. gwynne-vaughanii, T. teuchansii* and *A. mackiei* of a food-conducting tissue with some, but not all, of the key features of extant tracheophyte phloem suggests that the assembly of all phloem characteristics occurred after the origin of xylem. The FCCs of *T. teuchansii* and *A. mackiei* also imply that key features of extant phloem tissue were acquired from FCCs independently in the euphyllophytes and the extant lycopsids, through the narrowing of sieve elements and acquisition of the pericycle. FCCs and sieve pores were also acquired independently and convergently in multiple lineages of bryophyte^64-67^ (also see **Supplementary Note**), in the gametophyte generation. This evolutionary scenario suggests that sieve pores are not a sole defining feature of the phloem, but rather one of a suite of adaptations to sugar conduction. Furthermore, sieve pores are likely one of the earlier occurring adaptations to sugar conduction, owing to their appearance in an early-diverging polysporangiophyte lineage, and convergently in multiple bryophyte lineages. Enlarging plasmodesmata to form sieve pores also only requires co-opting a developmental programme already involved in plasmodesmata pit formation^37^ and therefore is potentially more readily originated than more developmentally complex traits. Following initial specialisation in sugar conduction through elongated cells with sieve pores like those of *A. mackiei*, later in tracheophyte evolution, and independently in the euphyllophyte and the extant lycophyte lineages, features arose associated with optimisation of sugar conduction, such as narrower and longer cells.

**Figure 4.**
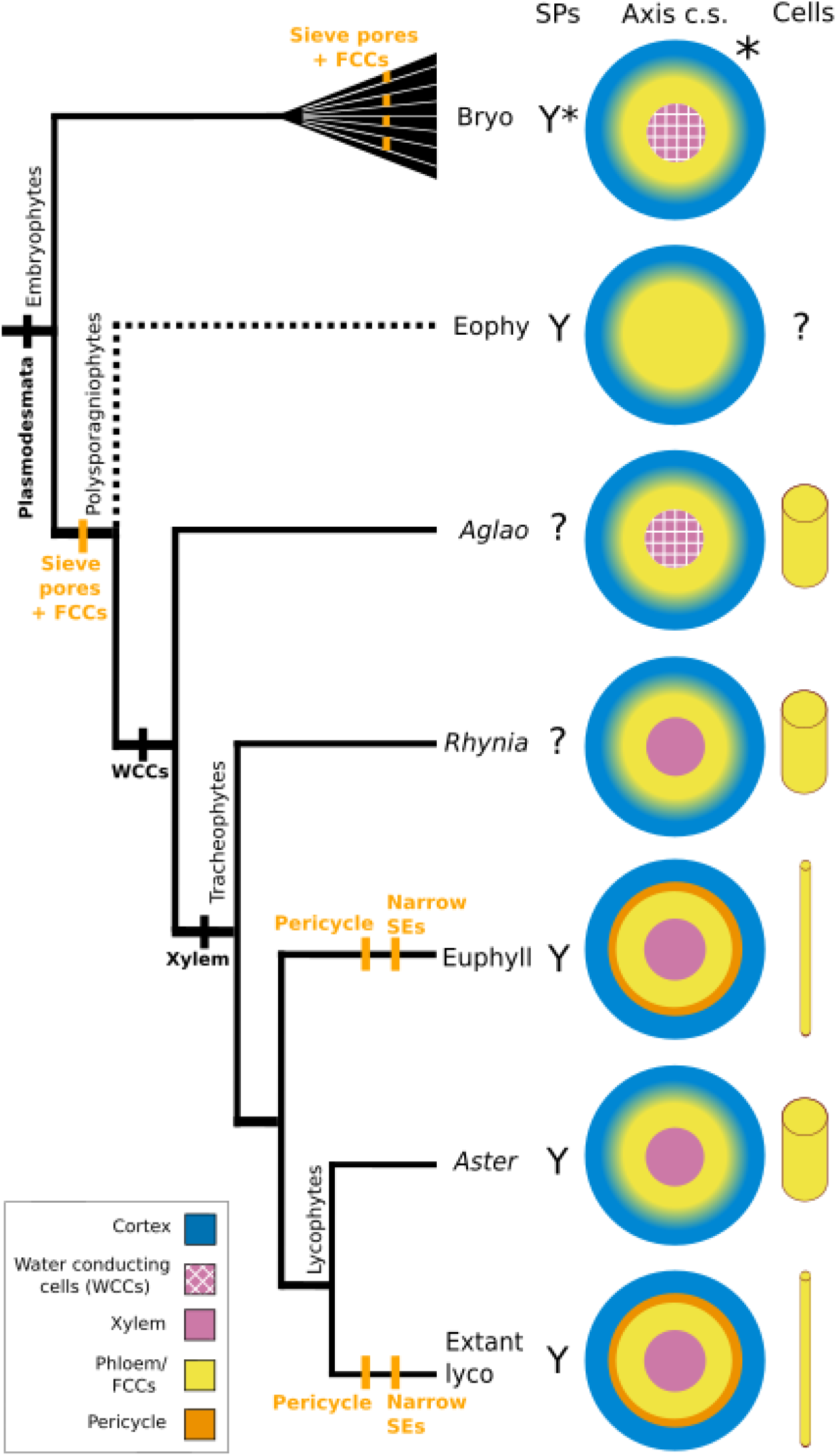
The characteristics of FCCs evolved gradually across polysporangiophytes, out of step with the evolution of xylem. Plasmodesmata were present in the ancestor of all embryophytes. The presence of FCCs and sieve pores in the eophytes that we propose implies that sieve pores originated in the common ancestor of eophytes + the rest of the polysporangiophytes, before the evolution of water conducting tissues. The occurrence of water-conducting cells (WCCs) in *Aglaophyton majus* implies that this trait was present in the common ancestor of *Aglaophyton majus* + tracheophytes. Xylem arose from WCCs in the common ancestor of extant tracheophytes. The presence in *Asteroxylon mackiei* of FCCs with sieve pores but without the narrow and very long cells of phloem and without a pericycle between the FCCs and the cortex (as indicated by the blurred cortex/FCC boundary in the axis schematics) implies that key features of extant phloem tissues were acquired from FCCs following the origin of xylem and independently in the euphyllophytes and the extant lycopsids. FCCs with sieve pores were also acquired convergently and independently in multiple lineages of bryophytes, in the gametophyte generation. This suggests that the sugar and water conducting tissues of tracheophytes followed distinct evolutionary trajectories. SPs = sieve pores, Y = sieve pores known; Axis c.s. = cross sectional schematic of axis tissue; Cells = schematic of cell dimensions; * = presence of trait in only some of the clade; dashed line on cladogram = uncertainty regarding phylogenetic placement. Taxon abbreviations = Bryo = Bryophytes; *Eoph* = *Eophytes; Aglao* = *Aglaophyton majus*; *Rhynia* = *Rhynia gwynne-vaughanii*; Euphyll = Euphyllophytes; *Aster* = *Asteroxylon mackiei;* Extant lyco = Extant lycophytes Colour key: blue = cortex, dashed magenta = WCCs, solid magenta = xylem, yellow = phloem or FCCs, orange = pericycle. Phylogeny based on ^17^, positions of *Aglaophyton majus* and *Rhynia gwynne-vaughanii* from^68^, position of eophytes from^25^.

The scenario outlined above is underpinned by interpretation of sieve pores as not the sole defining feature of phloem tissue, owing to the occurrence of similar structures in sugar-conducting tissues outside of the vascular plants, but rather favouring a more holistic definition of the tissue, incorporating histological, cellular and subcellular aspects. However, even if sieve pores are taken as the sole defining feature of phloem, our interpretation here that eophytes exhibited sieve pores also implies that phloem evolution was asynchronous to that of the xylem, as these plants lacked water-conducting tissues. Here we do not take this interpretation, owing to the histological and cellular differences between the FCCs of the Rhynie chert plants and the phloem of extant lycophytes suggesting that these were substantially distinct tissues. But regardless of how FCCs and phloem are distinguished precisely, this further demonstrates that the assembly of phloem and xylem traits were independent and asynchronous across the polysporangiophyte phylogeny.

In conclusion, the identification of FCCs in the Rhynie chert plants with only some of the features of the phloem of extant vascular plants demonstrates clearly that, whilst xylem and phloem are always found together in extant vascular plants, they did not originate simultaneously. Instead, also drawing on findings in other early polysporangiophytes^20, 25^, we propose a scenario in which the features of phloem evolved gradually from ancestral FCCs and asynchronously to that of xylem, with some features like sieve pores arising before the origin of xylem, and other associated features, such as the pericycle and narrow and very long sieve elements, arising later and independently in the euphyllophytes and the lycophytes.

*Species and organ abbreviations*: H. goe *stem =* Huperzia goebelli *stem;* H. sel *stem =* Huperzia sel *stem;* L. cla *root =* Lycopodium clavatum *root;* L. cla *stem =* Lycopodium clavatum *stem;* S. unc *root =* Selaginella uncinata *root;* S. unc *stem =* Selaginella uncinata *stem;* A. maj *axis =* Aglaophyton majus *axis;* R. gwy *axis =* Rhynia gwynne-vaughanii; T. teu *axis =* Trichopherophyton teuchansii *axis;* A. mac *leafy shoot =* Asteroxylon mackiei *leafy shoot;* A. mac *root-bearing axis =* Asteroxylon mackiei *root-bearing axis;* A. mac *rooting axis =* Asteroxylon mackiei *rooting axis*.

## Supporting information

Supplementary data images

## Lead contact

Further information and requests for resources should be directed to, and will be fulfilled by, the Lead Contact, Alexander J. Hetherington.

## Acknowledgments

We would like to thank I. Febbrari, Thin Sections and Lapidary Facility Manager, University of Edinburgh for the thin section preparation of MPEG slides. We would like to thank Y. Candela for assisting us in accessing the fossils of the National Museums Scotland collection, H. Kerp for access to the collections at the University of Münster, L. Loughtman for access to material in the Herbarium at the University of Manchester, and N. Clark for access to the Kidston collection at the Hunterian Museum at the University of Glasgow. We thank L. Pichevin for assistance with HF etching, and F. Laidlaw and A. Schofield for assistance with SEM imaging. We thank D. Kelly and T. McHugh at the Light Microscopy Core of the Wellcome Discovery Research Platform for Hidden Cell Biology, University of Edinburgh for access to the confocal imaging facilities. This work was supported by funding for the Wellcome Discovery Research Platform for Hidden Cell Biology (226791) and we gratefully acknowledge support from the Light Microscopy Core.

## Author Contributions

A.J.H. conceptualized the study. L.M.C. collected and analysed data. A.J.H. and L.M.C. wrote the paper.

## Materials availability

Thin sections and mounted peels examined in this study are housed in the following collections, collections abbreviations = Pb: Forschungsstelle für Palϊllobotanik, Institut für Geologie und Palϊllontologie, University of Münster, Germany. GLAHM Kid: Kidston Collection, The Hunterian, University of Glasgow, UK. BHUTTA: Akhlaq Ahmed Bhutta peel collection, University of Cardiff, UK. NHMUK: Natural History Museum, London, UK. MAN: Manchester Museum, University of Manchester, UK. OXF University Herbaria: Fielding-Druce Herbarium, University of Oxford, UK. LYON: Lyon Collection, University of Cardiff, UK. MPEG = Molecular Palaeobotany and Evolution Group, University of Edinburgh, UK. SCOTT = Andrew Scott Collection, University of Edinburgh, UK. STA = St Andrews Collection, University of St Andrews, UK. NMS = National Museums Scotland, Edinburgh, UK.

Accession numbers for all figured specimens: MPEG0015 (Fig. 1A-B), SCOTT EUCM 1361 (Fig. 1C), MAN EMu 547126 (Fig. 1D), LYON 93.11 (Fig. 1E-F), GLAHM Kid 2472 (Fig. 1G-H). MPEG0062 (Fig. 3F-I), MPEG0043 (Fig. 3J-M), NMS 1925.9.7 (Fig. S1A-B), NHMUK 15633 (Fig. S1C), STA355.76 (Fig. S1D), NHMUK 16433 (Fig. S1E-G), BHUTTA BL29A.185 (Fig. S2), STA 355.76 (Fig. S7A), STA 355.61 (Fig. S7B), MAN EMu 547408 (Fig. S7C), NHMUK 16433 (Fig. S7D).

All other data are available in the main text or supplementary materials.

## Data and code availability

- All data described in this paper is included within the paper and supplementary information.
- This paper does not report original code.

## Supplementary Figures and Captions

**Figure S1:**
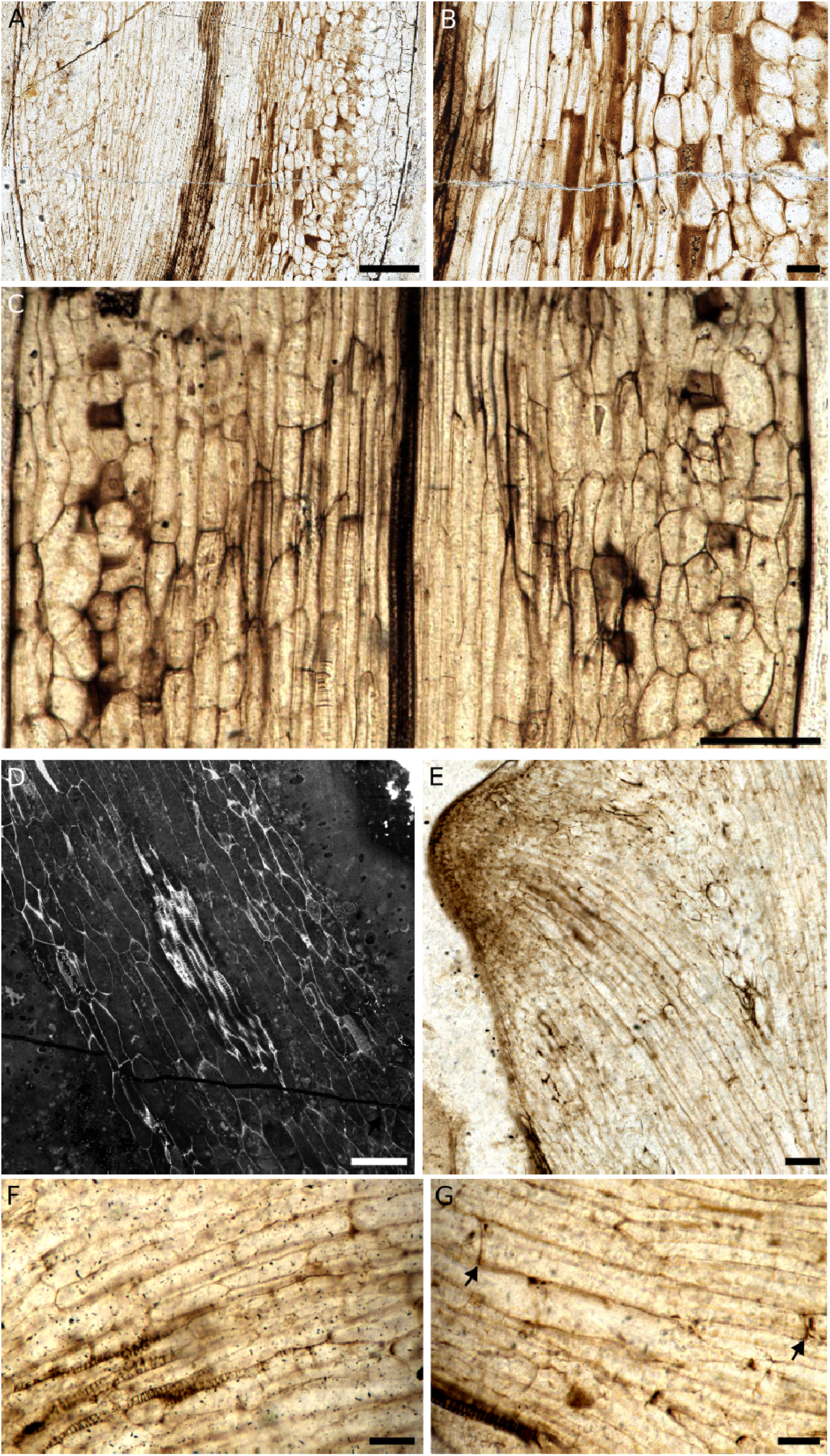
Longitudinal sections of food conducting cells (FCCs) of the Rhynie chert species shown in Figure 1. (A) Longitudinal section of Aglaophyton majus axis showing central WCCs and surrounding FCCs, transitional tissue and cortex. (B) Detail of the image in (A) showing WCCs (left), FCCs and transitional tissue (middle) and cortex (right). (C) Longitudinal section of Rhynia gwynne-vaughanii axis showing central xylem, surrounding FCCs, transitional tissue and cortical tissue. (D) CLSM of longitudinal section of Asteroxylon mackiei rooting axis showing central xylem, surrounding FCCs and cortical tissue. (E) Longitudinal section of Asteroxylon mackiei root-bearing axis with rooting axis meristem showing central FCCs and decayed cortex tissue, FCCs measuring 194-405 µm. (F) Detail of the specimen shown in (E) showing tracheids and elongated FCCs. (G) Detail of the specimen shown in (E) showing a 407 µm long FCC (arrows). Specimen codes = A-B = NMS 1925.9.7, C = NHMUK 15633, D = STA355.76, E-G = NHMUK 16433 Scale bars = 500 µm in (A), 200 µm in (C, D), 100 µm in (B, E), 50 µm in (F, G).

**Figure S2:**
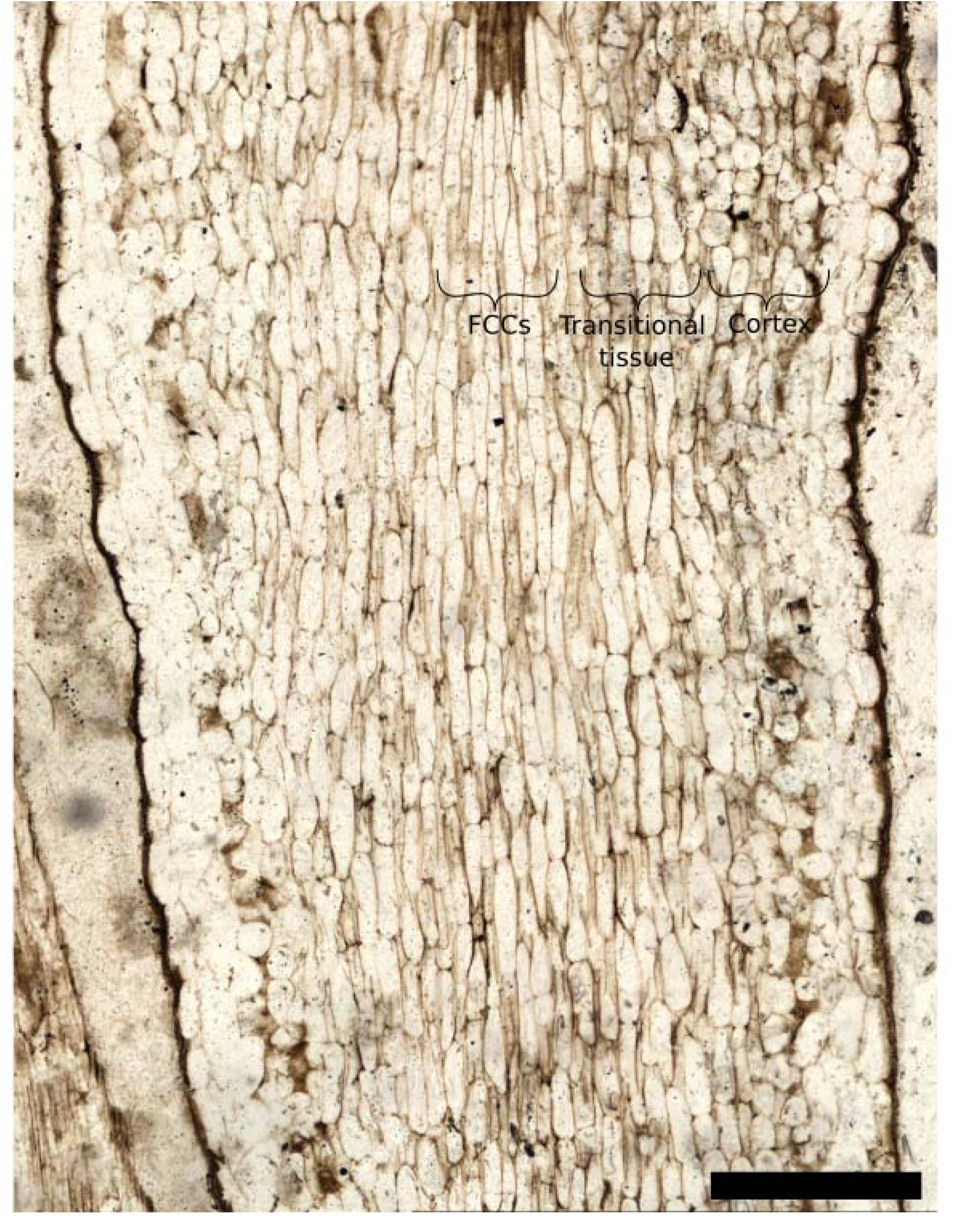
*Rhynia gwynne-vaughanii* (in cross section in Fig. 1 C-D) longitudinal section showing FCCs, transitional tissue and cortex cells. The FCCs are elongated and close packing, whilst the cells of the transitional tissue are slightly shorter and looser-packing, with air spaces between cells visible. The cortex cells are rounded with many intracellular spaces. Specimen code = BHUTTA BL29A.185. Scale bar = 500µm.

**Figure S3:**
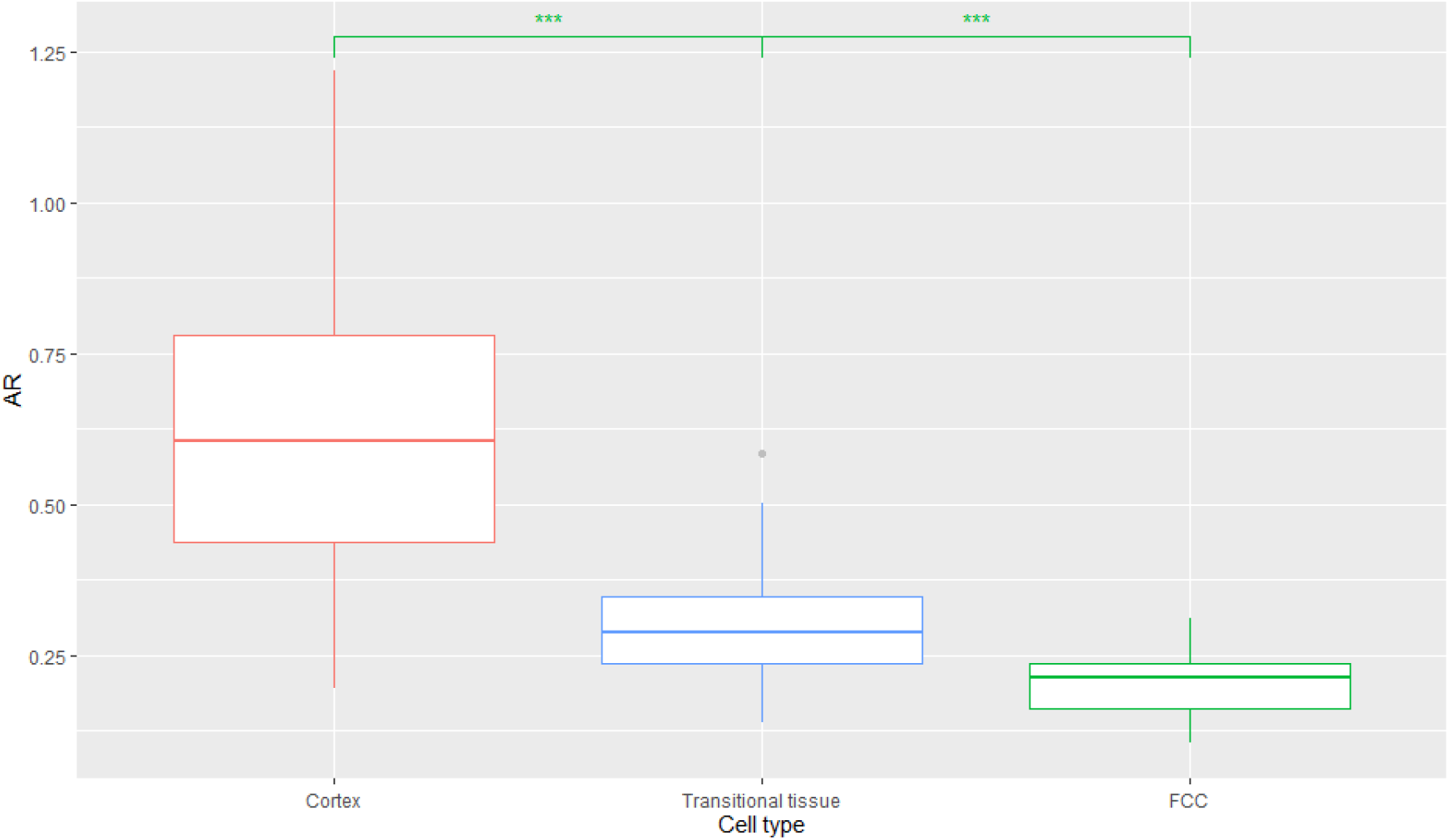
Aspect ratio (AR) of the parenchymatous cell types of the *Rhynia gwynne-vaughanii* axis in Figure S2 (c.s. of species in Figure 1 C-D). The FCCs have an average AR of 0.20 (range 0.31-0.10), indicating that these cells are narrow and elongated. Adjacent to the FCCs is a transitional tissue of elongated cells (average AR of 0.30, range 0.58-0.14) with some intercellular spaces, these cells have a significantly (p≤0.001, as demonstrated by a t-test) larger AR than the FCCs, as these cells are wider and shorter than the FCCs on average **(supplementary table 1)**. The cortex cells are more rounded than the cells of the transitional tissue and the FCCs (average AR of 0.62, range 1.22-0.20, significantly larger than the transitional tissue). Measurements from axis in Fig. S2.

**Figure S4:**
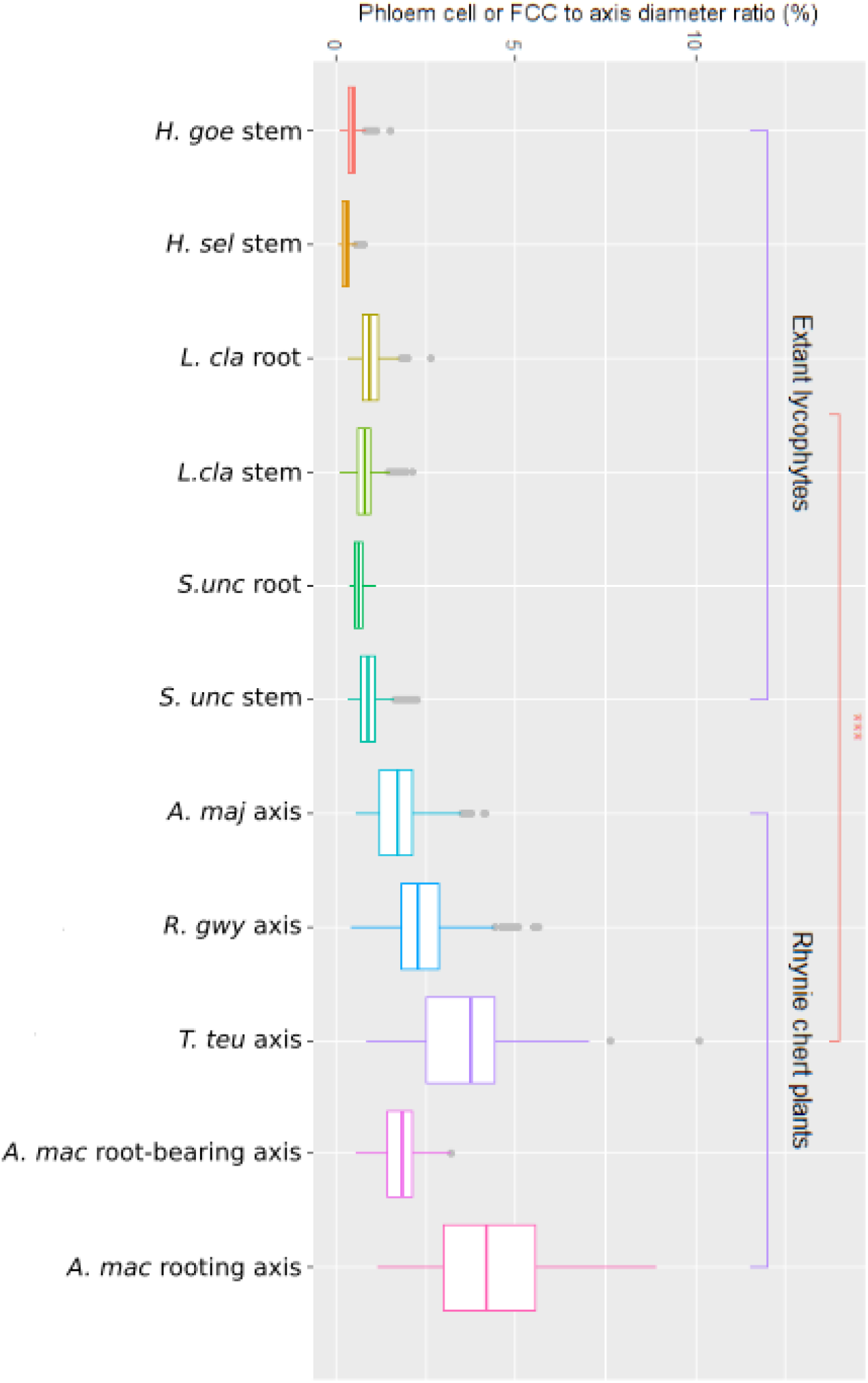
When differences in axis diameter are accounted for, the trend shown in Figure 2, that the FCCs of the Rhynie chert plants are significantly (p≤0.001) larger in diameter than the phloem cells of extant lycophytes, persists. Measurements of the diameter of phloem or food conducting cells expressed as a ratio to the axis diameter and presented as a percentage in extant lycophytes compared to extinct species from the Rhynie chert. Four extant lycophyte species (*Huperzia goebellii, Huperzia selago, Lycopodium clavatum, Selaginella uncinata*) were investigated covering both stem and root tissues. Four Rhynie chert species covering axis tissues (*Aglaophyton majus, Rhynia gwynne-vaughanii, Trichopherophyton teuchansii*), and root bearing axis and rooting axis (*Asteroxylon mackiei*). The FCCs of the Rhynie chert species are all significantly (p≤0.001) larger than the phloem cells of the extant lycophytes, as demonstrated by a t-test. *Species and organ abbreviations: H. goe stem = Huperzia goebelli stem; H. sel stem = Huperzia sel stem; L. cla root = Lycopodium clavatum root; L. cla stem = Lycopodium clavatum stem; S. unc root = Selaginella uncinata root; S. unc stem = Selaginella uncinata stem; A. maj axis = Aglaophyton majus axis; R. gwy axis = Rhynia gwynne-vaughanii; T. teu axis = Trichopherophyton teuchansii axis; A. mac root-bearing axis = Asteroxylon mackiei root-bearing axis; A. mac rooting axis = Asteroxylon mackiei rooting axis*.

**Figure S5:**
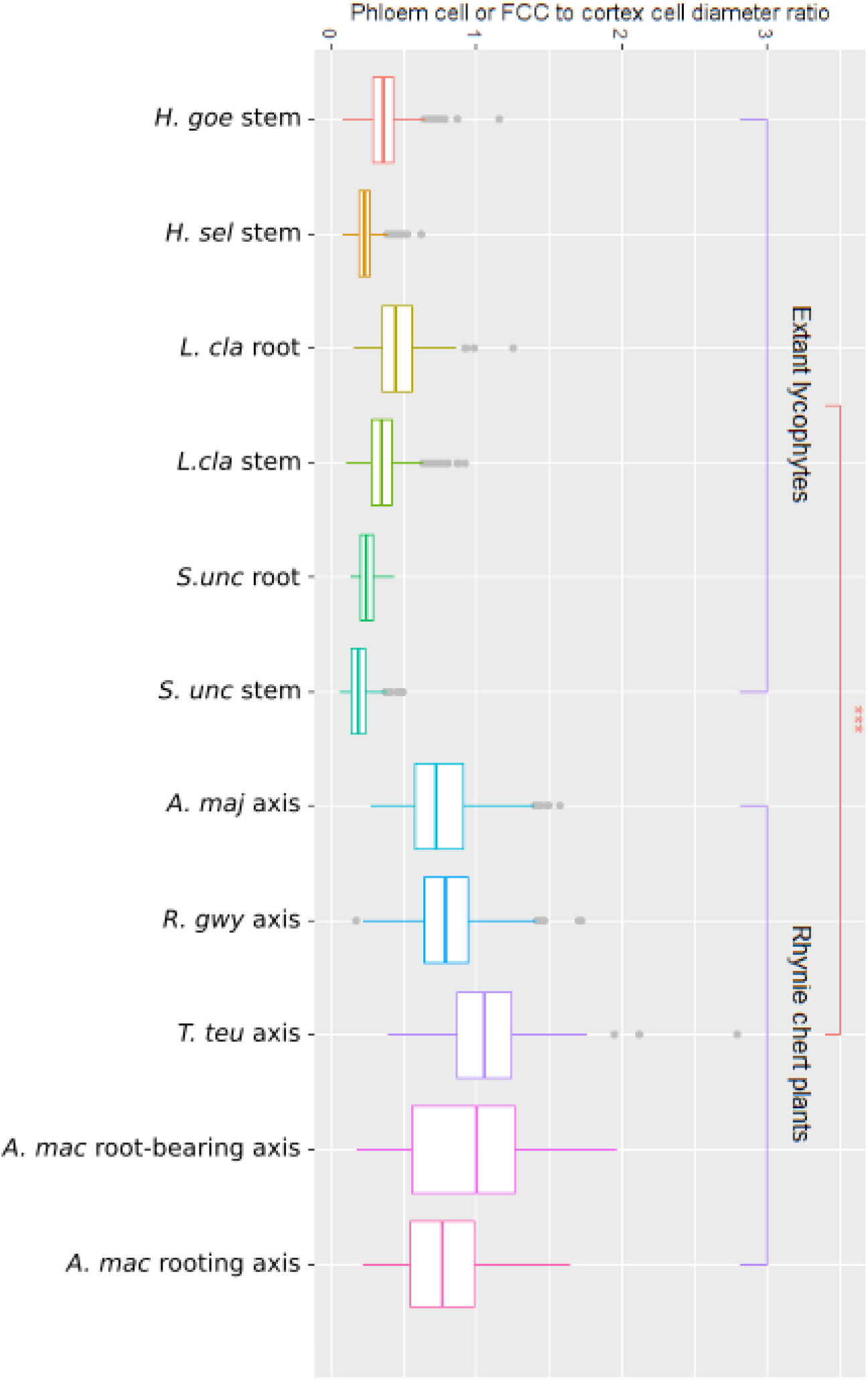
When differences in cell diameter are accounted for, the trend shown in Figure 2, that the FCCs of the Rhynie chert plants are significantly (p≤0.001) larger in diameter than the phloem cells of extant lycophytes, persists. Measurements of the diameter of phloem or food conducting cells expressed as a ratio to the average cortex cell diameter of the same axis in extant lycophytes compared to extinct species from the Rhynie chert. Four extant lycophyte species (*Huperzia goebellii, Huperzia selago, Lycopodium clavatum, Selaginella uncinata*) were investigated covering both stem and root tissues. Four Rhynie chert species covering axis tissues (*Aglaophyton majus, Rhynia gwynne-vaughanii, Trichopherophyton teuchansi*i), and root bearing axis and rooting axis (*Asteroxylon mackiei*). The FCCs of the Rhynie chert species are all significantly (p≤0.001) larger than the phloem cells of the extant lycophytes, as demonstrated by a t-test. *Species and organ abbreviations: H. goe stem = Huperzia goebelli stem; H. sel stem = Huperzia sel stem; L. cla root = Lycopodium clavatum root; L. cla stem = Lycopodium clavatum stem; S. unc root = Selaginella uncinata root; S. unc stem = Selaginella uncinata stem; A. maj axis = Aglaophyton majus axis; R. gwy axis = Rhynia gwynne-vaughanii; T. teu axis = Trichopherophyton teuchansii axis; A. mac root-bearing axis = Asteroxylon mackiei root-bearing axis; A. mac rooting axis = Asteroxylon mackiei rooting axis*.

**Figure S6:**
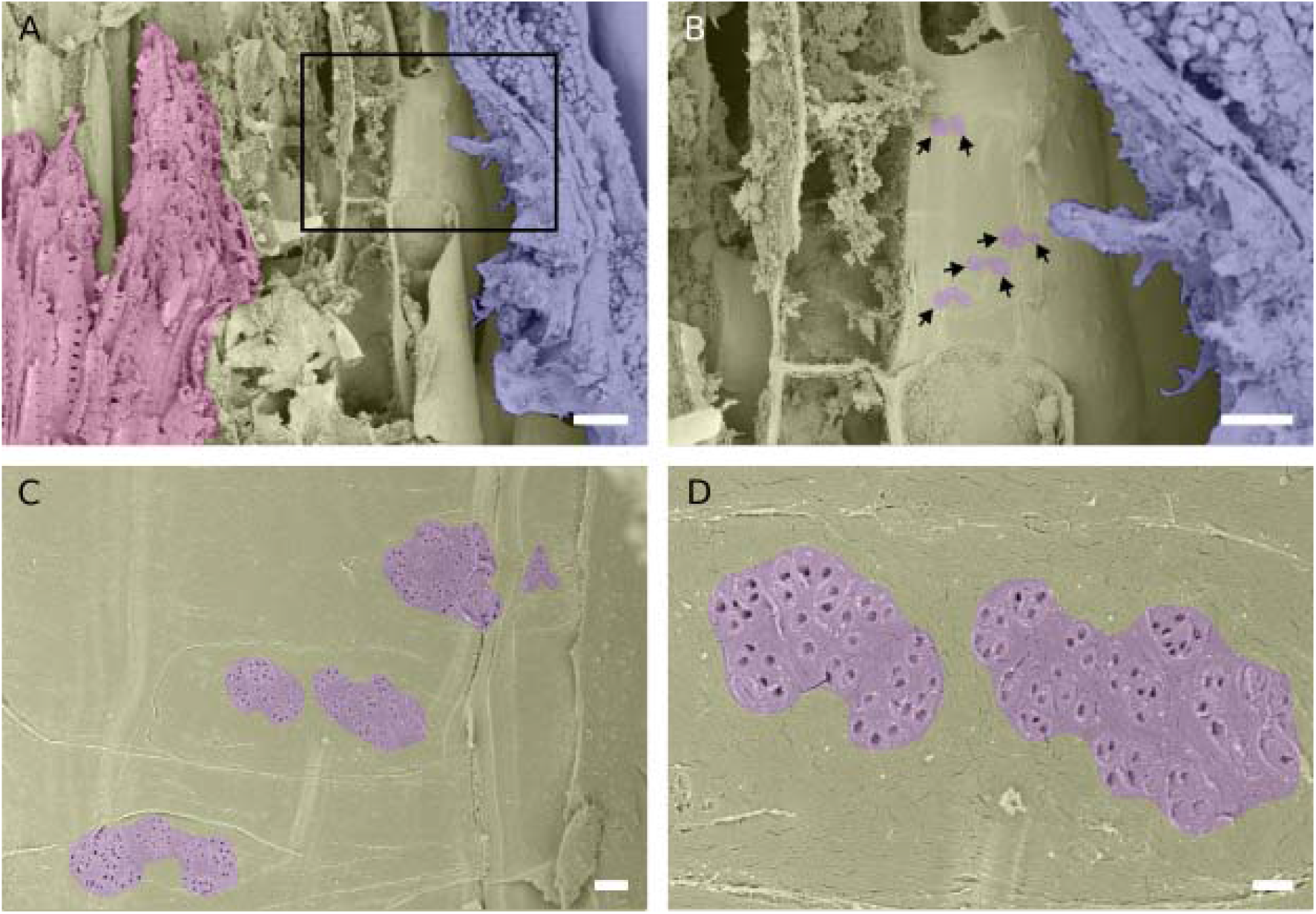
Plasmodesmata pits in the lateral walls of the phloem parenchyma of Huperzia selago are distinguishable from sieve pores (Figure 3B). A, SEM imaging of longitudinally cut *Huperzia selago* stem showing xylem (magenta false colouring), phloem tissue (yellow), and cortex (blue); B, higher magnification image of the region enclosed by the box in (A) showing phloem parenchyma lateral cell walls exhibiting multiple plasmodesmata pit areas (arrows, outlined in purple); C, higher magnification image showing the plasmodesmata pit areas in (B); D, higher magnification image of the central pit areas in (C) with individual plasmodesmata pits clearly visible. Scale bars = 20 µm in (A), 10 µm in (B), 1 µm in (C), 400nm in (D).

**Figure S7:**
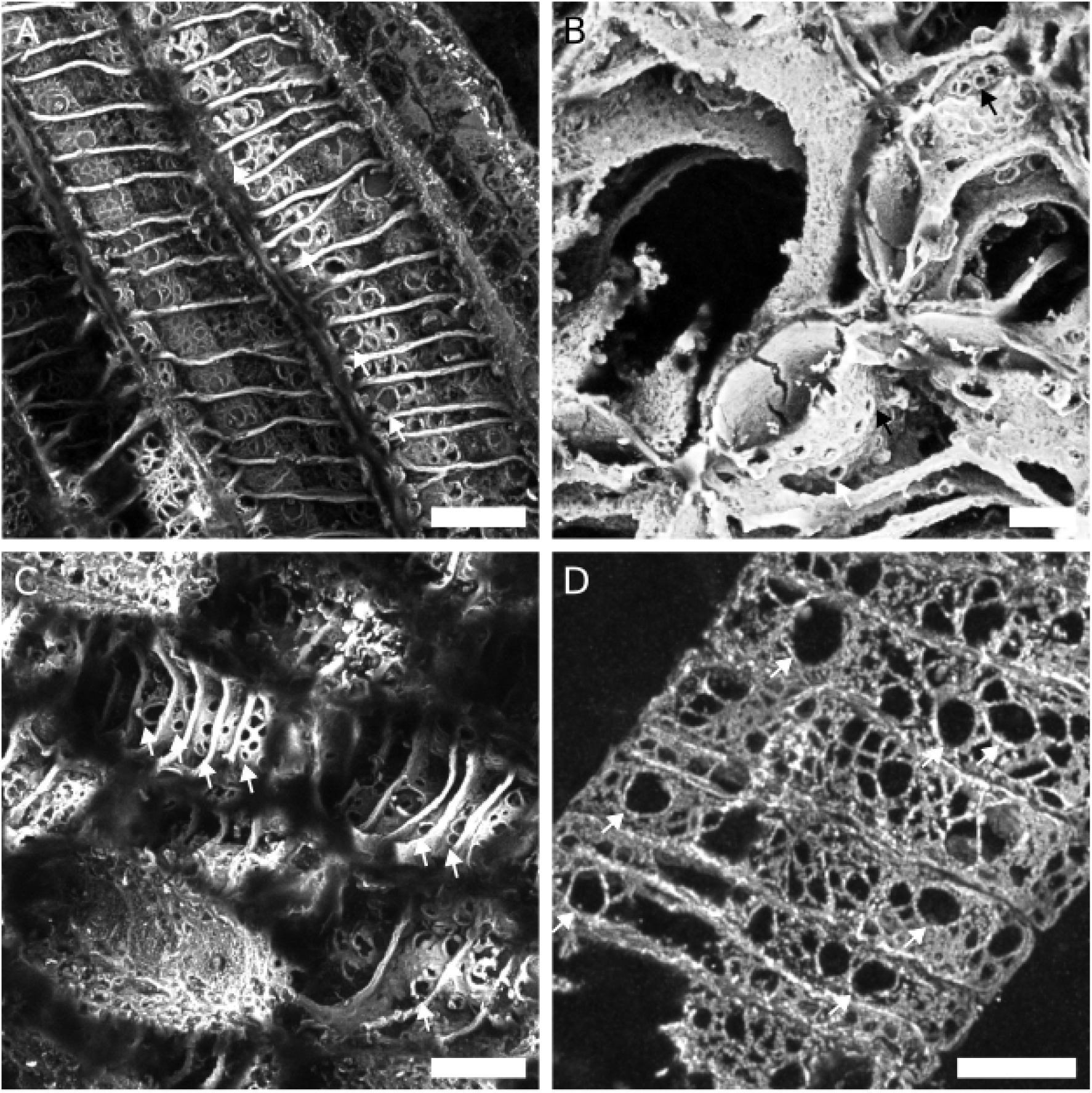
Test imaging shows that xylem pits of *Asteroxylon mackiei* are readily imaged using Airyscan CLSM, providing grounds for imaging of sieve pores in *Asteroxylon mackiei* (Fig. 3C-M). A, Xylem tracheids in longitudinal section B, Xylem tracheids in oblique-transverse section C, Xylem tracheids in oblique-longitudinal section D, Xylem tracheids in longitudinal section White arrows indicate xylem pits. Scale bars: 20 µm in (A, C), 10 µm in (B, D). Accession codes: STA 355.76 (A); STA 355.61 (B); MAN EMu 547408 (C); NHMUK 16433 (D).

**Supplementary Table 1:** Cell length, diameter and Aspect Ratio (AR) summary for *Rhynia gwynne vaughanii* parenchymatous tissue types. Measurements from the axis shown in Fig. S2, data shown in Fig. S3.

| | Average cell length ($\mu\text{m}$) | Cell length range ($\mu\text{m}$) | Average cell diameter ($\mu\text{m}$) | Cell diameter range ($\mu\text{m}$) | Average Aspect Ratio | Aspect Ratio range |
| --- | --- | --- | --- | --- | --- | --- |
| FCCs | 226.1 | 142.3-346.7 | 44.1 | 31.8-61.5 | 0.2 | 0.1-0.31 |
| Transitional Tissue | 189.6 | 93.9-282.1 | 53.7 | 3.1-72.6 | 0.3 | 0.14-0.58 |
| Cortex cells | 102.3 | 49.9-182.7 | 59.0 | 30.7-94.0 | 0.62 | 0.2-1.22 |

