## Supplementary text and methods for "Phloem evolved gradually and asynchronously to xylem in early vascular plants"

### KEY RESOURCES TABLE

#### TABLE FOR AUTHOR TO COMPLETE

***Please do not add custom subheadings.*** *If you wish to make an entry that does not fall into one of the subheadings below, please contact your handling editor or add it under the “other*” subheading*.* ***Any subheadings not relevant to your study can be skipped.*** *(****NOTE:*** *references should be in numbered style, e.g., Smith et al.^1^)*

**Key resources table**

| REAGENT or RESOURCE | SOURCE | IDENTIFIER |
| --- | --- | --- |
| Biological samples |  |  |
| *Huperzia goebellii* (Nessel) Holub (Lycopodiales)  stems | Royal Botanic Gardens Edinburgh, UK. | n/a |
| *Huperzia selago* (L.) Bernh. ex Schrank & Mart. (Lycopodiales) stems | Royal Botanic Gardens Edinburgh, UK. | n/a |
| *Lycopodium clavatum* L. (Lycopodiales)  stems and roots | Royal Botanic Gardens Edinburgh, UK. | n/a |
| *Selaginella uncinata* (Desv.) Spring (Selaginellales)  stems and roots | Royal Botanic Gardens Edinburgh, UK. | n/a |
| Software and algorithms | | |
| Fiji (Fiji Is Just ImageJ) 2.15 | Fiji | <https://github.com/fiji>; <https://imagej.net/software/fiji> |
| Other | | |
| Thin sections of *Aglaophyton (Rhynia) majus* (Kidston & Lang 1921a) D.S. Edwards 1986 | Rhynie chert, Aberdeenshire, UK. Block NSC.23, deposited in the University of Edinburgh, UK. | MPEG0015; MPEG0018 |
| Thin section of *Aglaophyton (Rhynia) majus* (Kidston & Lang 1921a) D.S. Edwards 1986 | Rhynie chert, Aberdeenshire, UK. Deposited in the University of Münster, Germany. | Pb1870 |
| Thin section of *Aglaophyton (Rhynia) majus* (Kidston & Lang 1921a) D.S. Edwards 1986 | Rhynie chert, Aberdeenshire, UK. Deposited in the National Museums Scotland, UK. | NMS 1925.9.7 |
| Thin section of *Aglaophyton (Rhynia) majus* (Kidston & Lang 1921a) D.S. Edwards 1986 | Rhynie chert, Aberdeenshire, UK. Deposited in Andrew Scott Collection, University of Edinburgh, UK. | SCOTT RC59, SCOTT RY36 |
| Thin section of *Aglaophyton (Rhynia) majus* (Kidston & Lang 1921a) D.S. Edwards 1986 | Rhynie chert, Aberdeenshire, UK. Block NSC.12, deposited in the University of Edinburgh, UK. | MPEG0007; MPEG0015 |
| Thin sections of *Rhynia gwynne-vaughani* Kidston & Lang 1917 | Rhynie chert, Aberdeenshire, UK. University of St Andrews, UK. | STA 424.23 |
| Thin sections of *Rhynia gwynne-vaughani* Kidston & Lang 1917 | Rhynie chert, Aberdeenshire, UK. University of Manchester, UK. | MAN EMu 547126 (R.1165) |
| Thin sections of *Rhynia gwynne-vaughani* Kidston & Lang 1917 | Rhynie chert, Aberdeenshire, UK. Deposited in the Bhutta Collection, University of Cardiff, UK. | BHUTTA BL29A.185 (Peel series BL29A, mounted peel 185). |
| Thin sections of *Rhynia gwynne-vaughani* Kidston & Lang 1917 | Rhynie chert, Aberdeenshire, UK. Deposited in the Andrew Scott Collection, University of Edinburgh, UK. | SCOTT EUCM1361 |
| Thin sections of *Rhynia gwynne-vaughani* Kidston & Lang 1917 | Rhynie chert, Aberdeenshire, UK. Deposited in the University of Münster, Germany. | Pb5007; Pb1588; Pb5006; Pb1809; Pb1876 |
| Mounted peels of *Trichopherophyton teuchansii* Lyon & Edwards 1991 | Rhynie chert, Aberdeenshire, UK. Deposited in the Lyon Collection, University of Cardiff, UK. | LYON 93.11 and 93.15 (block 93, mounted peels 11 and 15) |
| Thin section of *Asteroxylon mackiei* Kidston & Lang 1921b | Rhynie chert, Aberdeenshire, UK. Deposited in the Natural History Museum, London, UK. | NHMUK 16433 |
| Thin section of *Asteroxylon mackiei* Kidston & Lang 1921b | Rhynie chert, Aberdeenshire, UK. Deposited in the Hunterian Museum, University of Glasgow, UK. | GLAHM Kid 2472 |
| Thin sections of *Asteroxylon mackiei* Kidston & Lang 1921b | Rhynie chert, Aberdeenshire, UK. From the University of St Andrews, UK. | STA 355.61; STA 355.76 |
| Thin sections of *Asteroxylon mackiei* Kidston & Lang 1921b | Rhynie chert, Aberdeenshire, UK. Deposited in the University of Manchester, UK. | MAN EMu 547408 (R.1193); MAN EMu 547404 (R.1190) |
| Thin sections of *Asteroxylon mackiei* Kidston & Lang 1921b | Rhynie chert, Aberdeenshire, UK. Block NSC.10, deposited in the University of Edinburgh, UK. | MPEG0062; MPEG0043 |
| Thin sections of *Asteroxylon mackiei* Kidston & Lang 1921b | Rhynie chert, Aberdeenshire, UK. Deposited in the Natural History Museum, UK. | NHMUK 16433 |
| Thin section of *Asteroxylon mackiei* Kidston & Lang 1921b | Rhynie chert, Aberdeenshire, UK. Deposited in the University of Oxford, UK. | OXF University Herbaria 112 (423.7) |
| Block NSC10 containing *Asteroxylon mackiei* Kidston & Lang 1921b | Rhynie chert, Aberdeenshire, UK. | NSC10 |
| *Asteroxylon mackiei* Kidston & Lang 1921b axes on SEM stubs from block NSC10 | Rhynie chert, Aberdeenshire, UK. | n/a |

##### **Experimental model and subject details**

The Rhynie chert is a hot spring deposit in Aberdeenshire, UK, which dates to the Lower Devonian (late Pragian to earlier Emsian^69^ around 407 million years old^8, 70-73^). It preserves a wetland valley ecosystem through silica permineralization, and the organisms preserved here include a wide diversity of early terrestrial life^8, 74^. The Rhynie chert is especially known for the exceptional quality of its preservation^8,9,75,76^, with organisms preserving delicate tissues and features rarely found at other fossil deposits (e.g.,^77-80^). The land plants described from the Rhynie chert encompass much of the diversity of early Devonian land plants (with the exception of the bryophytes and the euphyllophytes)^10^; the species used in this study are the non-vascular polysporangiophyte *Aglaophyton (Rhynia) majus* (Kidston & Lang 1921a) D.S. Edwards 1986^12, 16^, the early vascular plant *Rhynia gwynne-vaughani* Kidston & Lang 1917^11, 12^*,* and the lycophytes *Trichopherphyton teuchansii* Lyon & Edwards 1991 (a zosterophyll^18^) and *Asteroxylon mackiei* Kidston & Lang 1921b (a lycopsid^13^). These four species span the origin of the vascular plant lineage^11, 12, 16, 17^, the divergence of the lycophytes from the euphyllophytes, and the diversification of the lycophytes^10, 13, 17, 18, 19^, and therefore the occurrence of traits in these fossil taxa can help infer the ancestral condition of these traits in the corresponding polysporangiate plant groups. This has been done previously in the context of using the traits of *A. mackiei* and other Rhynie chert plants to make inferences about the ancestral rooting system of vascular plants^33^. In order to place findings in these extinct species in their wider evolutionary context, extant lycophytes from two of the three extant order (Lycopodiales (*Lycopodium clavatum* L., *Huperzia selago* (L.) Bernh. ex Schrank & Mart., *Huperzia goebellii* (Nessel) Holub) and Selaginellales (*Selaginella uncinata* (Desv.) Spring)) were used for anatomical comparison.

##### **Method details**

Thin section preparation

All Molecular Palaeobotany and Evolution Group (MPEG) Rhynie chert thin sections in this study were prepared by Ivan Febbrari in the Thin Sections and Lapidary Facility, School of Geosciences, University of Edinburgh. The thin sections MPEG0015 and MPEG0018; MPEG0007; MPEG0062 and MPEG0043 were produced from NSC blocks (NSC.23, NSC.12, and NSC.10, respectively). These blocks were originally collected in farmland owned by the Windyfield Farm (NJ 349642 mE, 827852 mN) adjacent to the Rhynie chert site of special scientific interest (SSSI) by a local land owner before 2021. Specimens were passed to North Sea Core to help distribute the samples for academic research with the mutual agreement of NatureScot. Blocks were distributed using the accession numbers North Sea Core NSC.01-NSC.45. The thin sections were prepared with a coverslip and thicknesses of 30µm, with the exception of MPEG0062 (50µm thickness).

Histological preparation

Samples of stems and roots of *Huperzia selago* and *H. goebelli, Lycopodium clavatum* and *Selaginella uncinata* were collected from RBGE and fixed in FAA (5ml 37% formaldehyde: 25 ml 96% ethanol: 2.5ml concentrated glacial acetic acid, made up to 50ml with sterile water) immediately after harvesting, then placed under vacuum at 500mbar until samples had sunk. The samples were dehydrated with an ethanol series and stored in 70% ethanol, then embedded in wax, sectioned and stained with Astra Blue and Safranin.

Brightfield Microscopy

Thin sections and histology slides were observed and photographed with a Nikon ECLIPSE LV100D compound microscope, a Nikon SMZ18 Stereoscope and a Keyence VHX-X1.

Confocal Laser Scanning Microscopy

A Zeiss LSM 880 confocal microscope with Airyscan in the Light Microscopy Core of the Wellcome Discovery Research Platform for Hidden Cell Biology, University of Edinburgh was used to produce regular and Airyscan CLSM images. Single plane CLSM images were taken with the following settings: x40 oil immersion lens, 488nm and 561nm lasers, a long pass emission window, a 1Au pinhole and image stitching. Airyscan CLSM z-stack images were taken with the following settings: x40 oil immersion lens, 488nm and 561nm lasers, a long pass emission window and a 2.5 Au pinhole. Z-stack images are presented as maximum intensity projections produced in Fiji.

Scanning Electron Microscopy

*Asteroxylon mackiei*

*Asteroxylon mackiei* samples prepared for SEM originated from fragments of the NSC.10 block (provenance as for the NSC blocks above). Samples of *A. mackiei* were selected for SEM through fracturing of small chert samples containing *A. mackiei* from block NSC.10 with hammer and chisel, and five samples of <2cm with surface-exposed plant vasculature were selected. All samples were photographed before etching to document content of samples. HF etching was carried out in the Oceanography Clean Laboratory (School of Geosciences, University of Edinburgh) with Dr Laetitia Pichevin. Samples were placed with tweezers into Teflon containers and 0.6ml 12N HCl was first added to all samples to prevent deposition of fluoride residues. Then ~1.2ml 28N concentrated hydrofluoric acid was added to cover the samples. Samples were left for varying lengths of time (10, 15, 20 and 25 minutes) before boric acid was added to stop the reaction. The sample shown in **Figure 3 C-E** were exposed to HF for 20 minutes. The resulting mixture was then removed with a pipette from the containers and samples were washed gently with water twice to ensure safety of the sample and to remove residue. Samples were then removed with tweezers from the Teflon containers, with care taken to ensure orientation of the samples remained consistent and exposed tissue was not damaged, and dried on paper towels. Once dry, samples were adhered to SEM stubs using carbon black tabs, examined using a stereoscope to ensure that the correct orientation was maintained, then secured using carbon cement. Samples were photographed again before coating to identify the tissues exposed in the axes. Samples gold alloy sputter coated at 30mA for 3 minutes initially, and samples which exhibited charging were coated again for another minute.

Extant lycophytes

Samples of *Lycopodium clavatum* and *Huperzia selago* stems were harvested from RBGE and fixed in FAA (5ml 37% formaldehyde: 25 ml 96% ethanol: 2.5ml concentrated glacial acetic acid, made up to 50ml with sterile water) immediately after harvesting, then placed under vacuum at 500mbar until samples had sunk. Samples were then dehydrated through an ethanol series to 100% ethanol, then subject to critical point drying. Samples were then cut horizontally through the midpoint of the stem with a razor blade to expose the vascular tissue. Samples were then gold alloy sputter coated at 30mA for 3 minutes, and imaged using a JEOL JSM-6010 SEM for initial scanning of samples, then a Zeiss Crossbeam 550 FIB-SEM for taking higher resolution images (both in the School of Physics, University of Edinburgh).

**Quantification and Statistical Analysis**

Extant lycophytes

Phloem cell diameter was measured from all of the cells of the phloem tissue (therefore including both phloem parenchyma and sieve elements), to ensure a fair comparison with the Rhynie chert taxa, where different cell types within the FCCs have not been described. Only those cells which lacked evidence of deformation were measured, and the widest point of the cell was measured.

Cortex cell diameter was measured as, for a given sample, a rectangle whose short end was the same size as the diameter of the stele was drawn from the stele out to the stem or root epidermis in a region where preservation was determined to be good, and here all the cell without evidence of deformation were measured (except for *Selaginella uncinata*, where there was some deformation to all cortex cells, the least deformed cells were measured here i.e., that exhibited less warping of the cell walls). In all cases the widest point of the cells was measured. For phloem cell to cortex cell diameter ratios, average cortex cell diameter was calculated for each species and organ type.

Axis diameter was measured from at least four measurements of the axis radius, aiming to capture the largest and smallest points of the axis radius (where deformation was absent), the average radius was taken and this was doubled for the diameter. For phloem cell to axis diameter ratios, average axis diameter was calculated for each species and organ type.

Total extant lycophyte samples measured: *Huperzia goebelli* stems (five), *Huperzia selago* stems (five), *Lycopodium clavatum* stems (five) and roots (four), *Selaginella uncinata* root (one) and stem (five). Total number of extant lycophyte axes samples = 25. Total number of extant lycophyte phloem cells measured = 9947.

Rhynie samples

FCC diameter: Cells that were identified as FCCs in cross section were those that were adjacent to the WCC/ xylem strand and close-packing, and the cortex cells taken as the cells with intracellular spaces, initially smaller but increasing in size when moving peripherally. The widest point of the cell was measured, and all cells identified as FCCs were measured owing to the limited number of well-preserved axes and to capture variation in cell size within an axis.

Cortex cell diameter: For a given sample, a rectangle whose short end was the same size as the diameter of the vasculature strand was drawn out from the vasculature to the edge of the axis in a region where preservation was determined to be good, and here all the cells without evidence of deformation were measured from the widest point of the cell.

Axis diameter: Several measurements of the axis radius were taken, aiming to capture the largest and smallest points of the radius (where deformation was absent) where possible, average taken, and doubled for the diameter. For FCC to axis diameter ratios, average axis diameter was calculated for each species and organ type.

Rhynie chert plant axes measured: *Aglaophyton* *majus* (five), *Rhynia* *gwynne-vaughanii* (eight), *Trichopherophyton teuchansii* (two), *Asteroxylon mackiei* leafy shoot (one), *Asteroxylon mackiei* root-bearing axes (two), *Asteroxylon mackiei* rooting axes (three). Total number of Rhynie chert plant axes sampled = 21. Total number of Rhynie chert plant FCCs measured = 2907.

Analysis

FCC and phloem cell diameters for the Rhynie chert plants and the extant lycophytes, respectively, were plotted as box plots in R with the replicates for each species and organ type pooled, and the p value between the Rhynie chert and the extant lycophyte groups is presented in **Figure 2**. In order to test if the difference in FCC and phloem cell diameters was due to the observed larger axis diameter of the Rhynie chert plants compared to the extant lycophytes, the ratio of the FCC or phloem cell diameters to the axis diameter was calculated as a percentage for each sample, using the average of the axis diameter measurements for each sample. The replicates for each species and organ type were then pooled, plotted as box plots in R, and the p value between the Rhynie chert and the extant lycophyte groups is presented in **Figure S4**. In order to test if the difference in FCC and phloem cell diameters was due to the observed larger overall cell diameters of the Rhynie chert plants compared to the extant lycophytes, the ratio of the FCC or phloem cell diameters to the cortex diameter was calculated for each sample, using the average of the cortex cell diameter measurements for each sample. The replicates for each species and organ type were then pooled, plotted as box plots in R, and the p value between the Rhynie chert and the extant lycophyte groups is presented in **Figure S5**.

**Supplementary Note: Interpretation of enlarged plasmodesmatal pits in food conducting tissues**

Edwards *et al* 2022 report and present images of “pits” in the walls of the FCCs of the eophytes. They describe these pits as “fall[ing] within plasmodesmata dimensions” but are “too small to measure accurately from SEM images”. However, we believe that the authors of Edwards *et al* 2022 do have images of sufficient quality to measure these pits. Measurement of the pits shown in Figure 5h of Edwards *et al* 2022 by the same methodology as the measurements of sieve pores in this paper finds these pits to have an average diameter of 0.17 µm (range 0.08 to 0.40 µm). As shrinkage of 14-47% has been reported in experimental charcoalification of seed plant organs^81^, applying shrinkage correction by multiplying by 1.27 suggests these structures were originally 0.1 to 0.51 µm, with an average of 0.22 µm. This average is closer to the diameters of lycophyte sieve pores reported in this paper (average of 0.15 µm) than to the diameters of plasmodesmatal pits in phloem parenchyma (average of 0.07 µm). Therefore, these pits are, on the whole, enlarged beyond plasmodesmata dimensions, suggesting they are sieve pores owing to their occurrence in putative sugar conducting tissue.

Similarly, some bryophyte groups exhibit, in the gametophytic generation, cells specialised in sugar conduction, characterised by elongated cells, a central axis position and enlarged plasmodesmatal openings, which have been described as sieve elements (though not as phloem)^65, 66^. These structures are termed leptoids in the polytrichales mosses, where the cells most closely resemble the sieve elements of phloem in their cell architecture and cytological features^64^. Leptoids or similar but less anatomically specialised FCCs appear in the Bryiales, Polytrichales and Sphagnales moss lineages^67^, and a similar tissue is also found in the Marchantiales liverworts^82^ and *Takakia*^66^. The enlarged plasmodesmatal pores in the polytrichaceous moss leptoids have been reported as ca. 100-200nm in diameter^66^ and under 0.2 µm^64^. Additionally, food conducting cells that have also been described as sieve elements, with enlarged plasmodesmatal pits reaching the range of lycophyte sieve pores and in some cases exceeding it, are also found across a diversity of red and brown algal groups, where the tissue also arose independently^83^. In these cases, it is argued that enlarged plasmodesmata that fall within the range of the sieve pores of vascular plants should be considered as sieve pores, owing to their role in sugar conduction, but that the term “sieve pore” should not imply homology with the sieve pores of vascular plants.
