## Supplementary data images for "Phloem evolved gradually and asynchronously to xylem in early vascular plants"

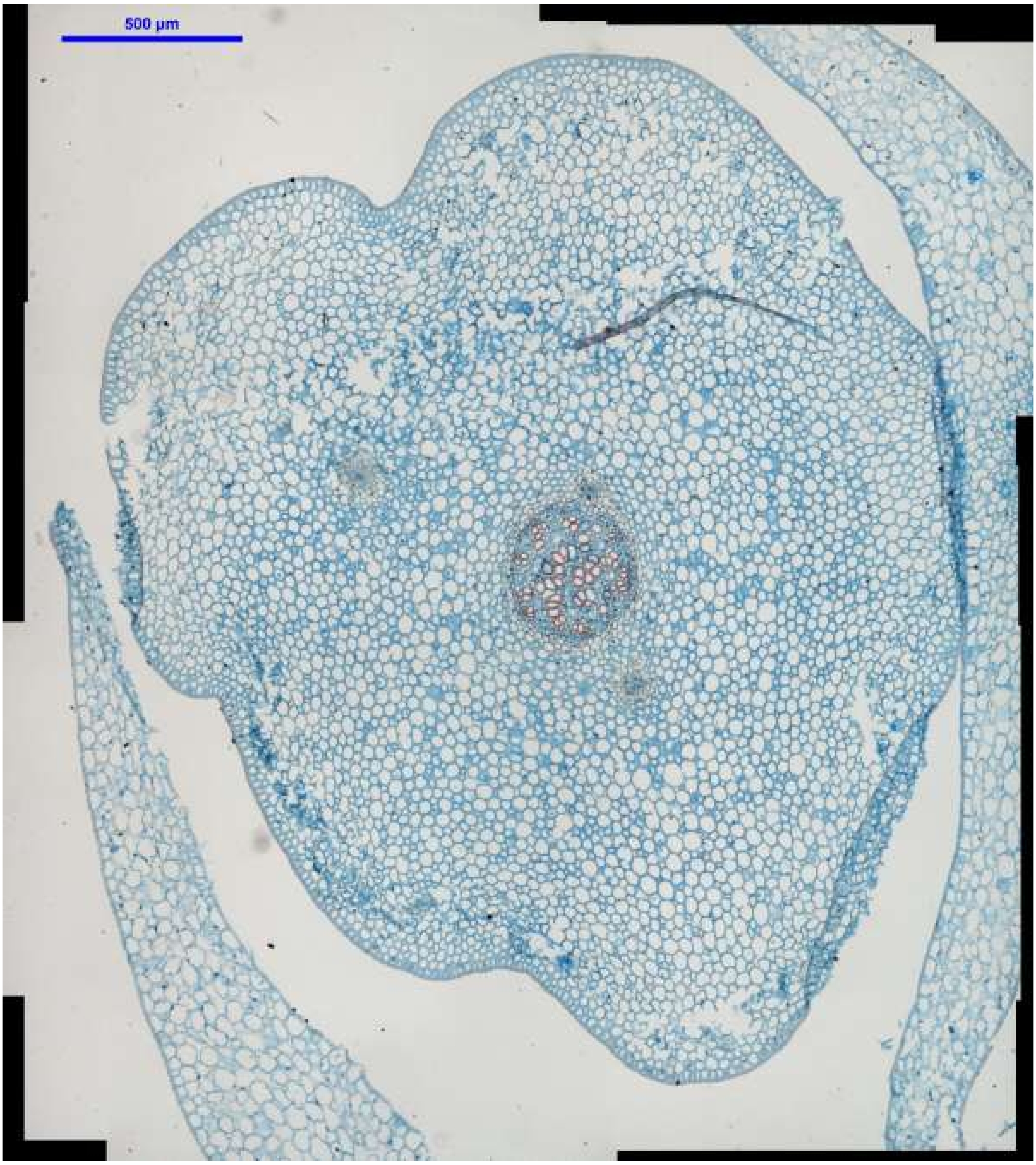
500 µm

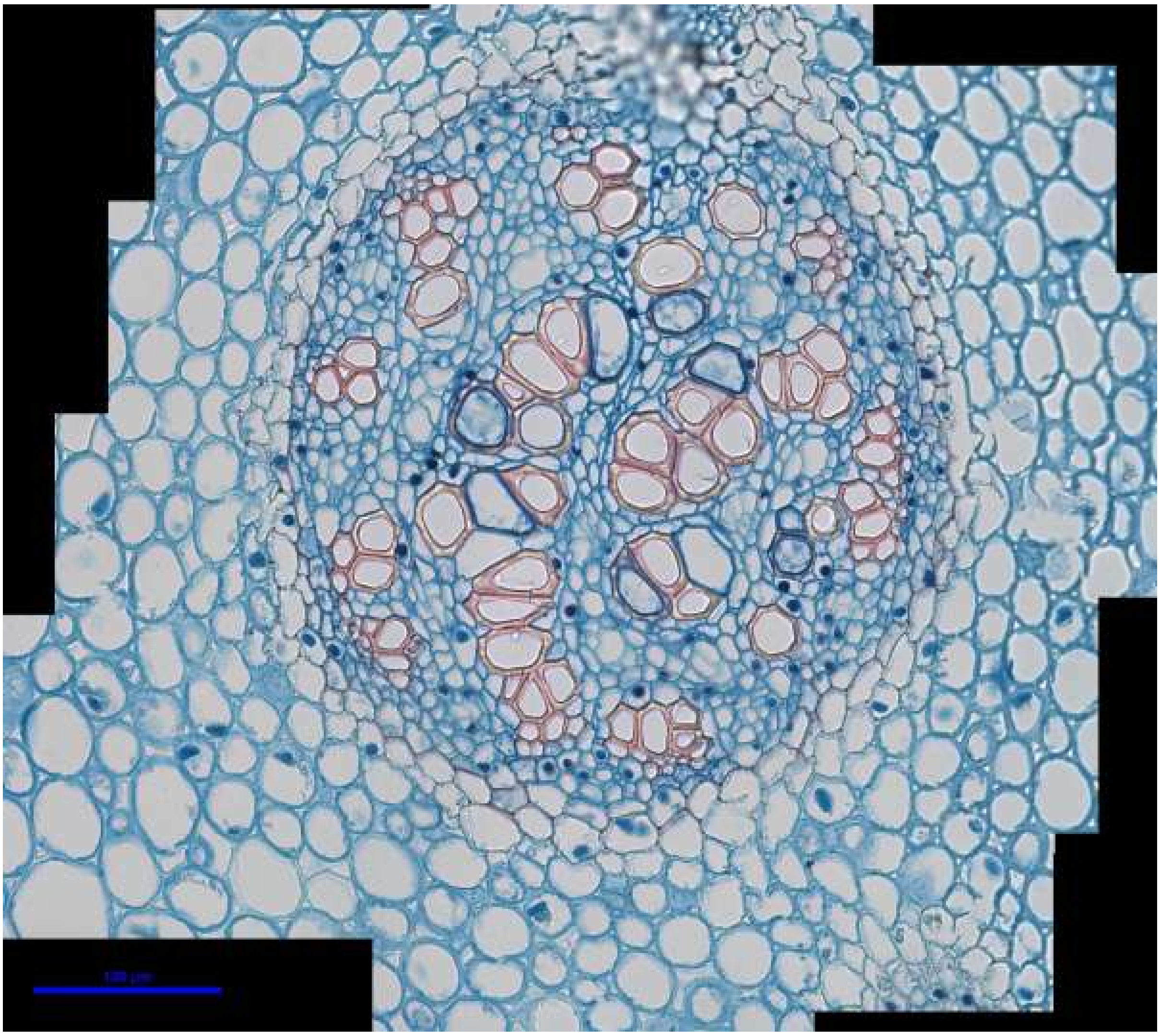

*Huperzia goebelii* stem: H_goeb_1

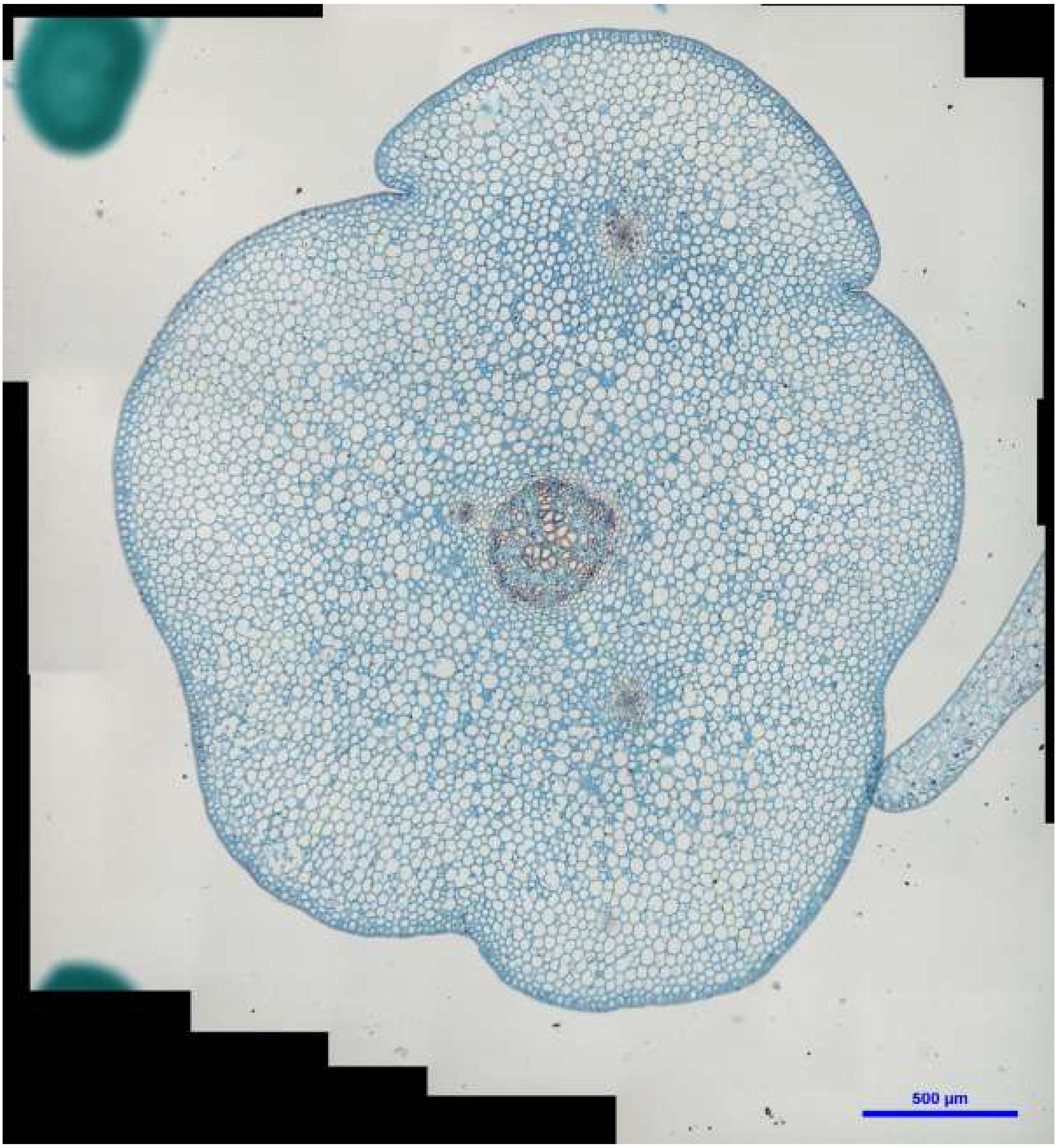
500 µm

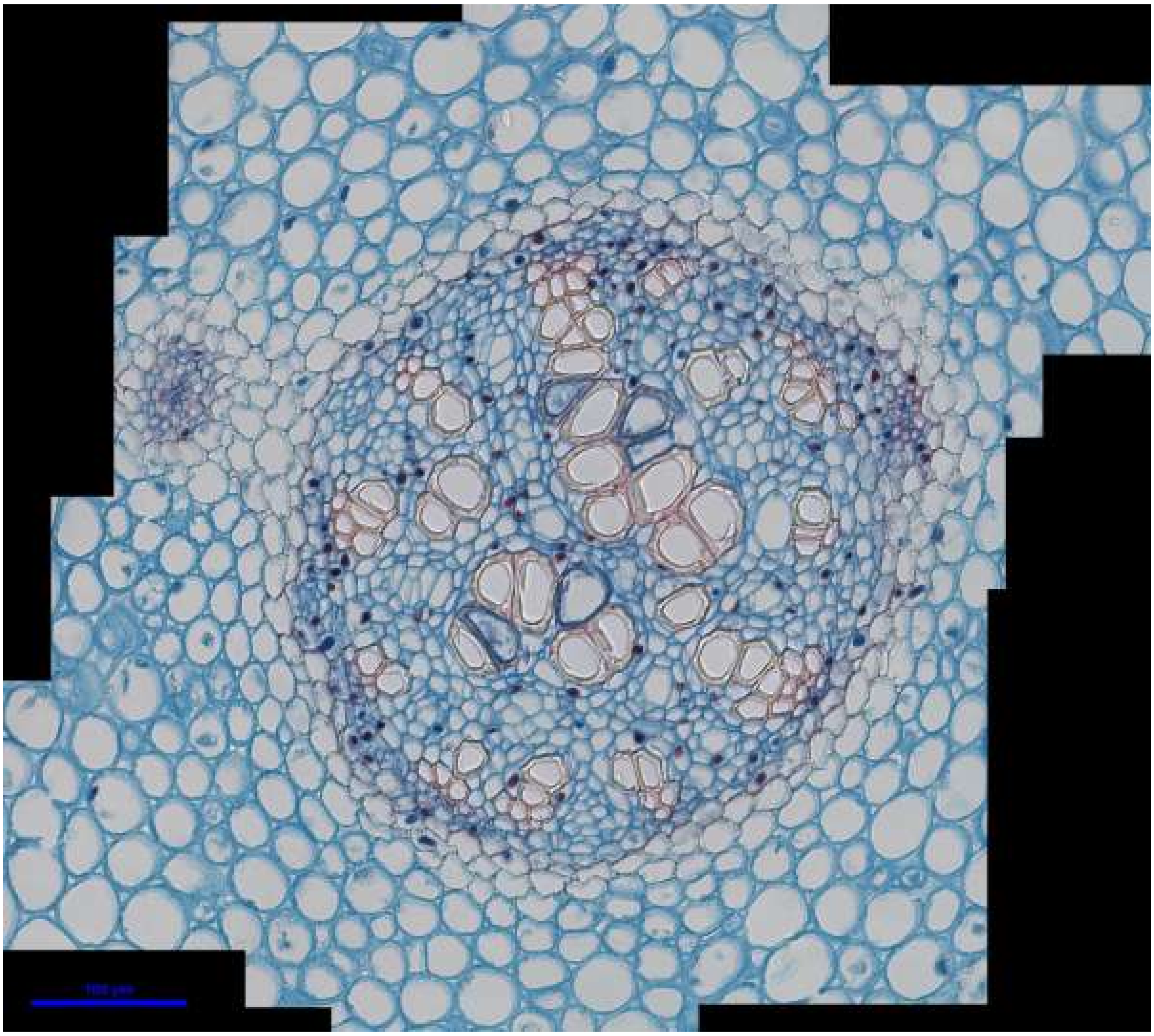

*Huperzia goebelii* stem: H_goeb_2

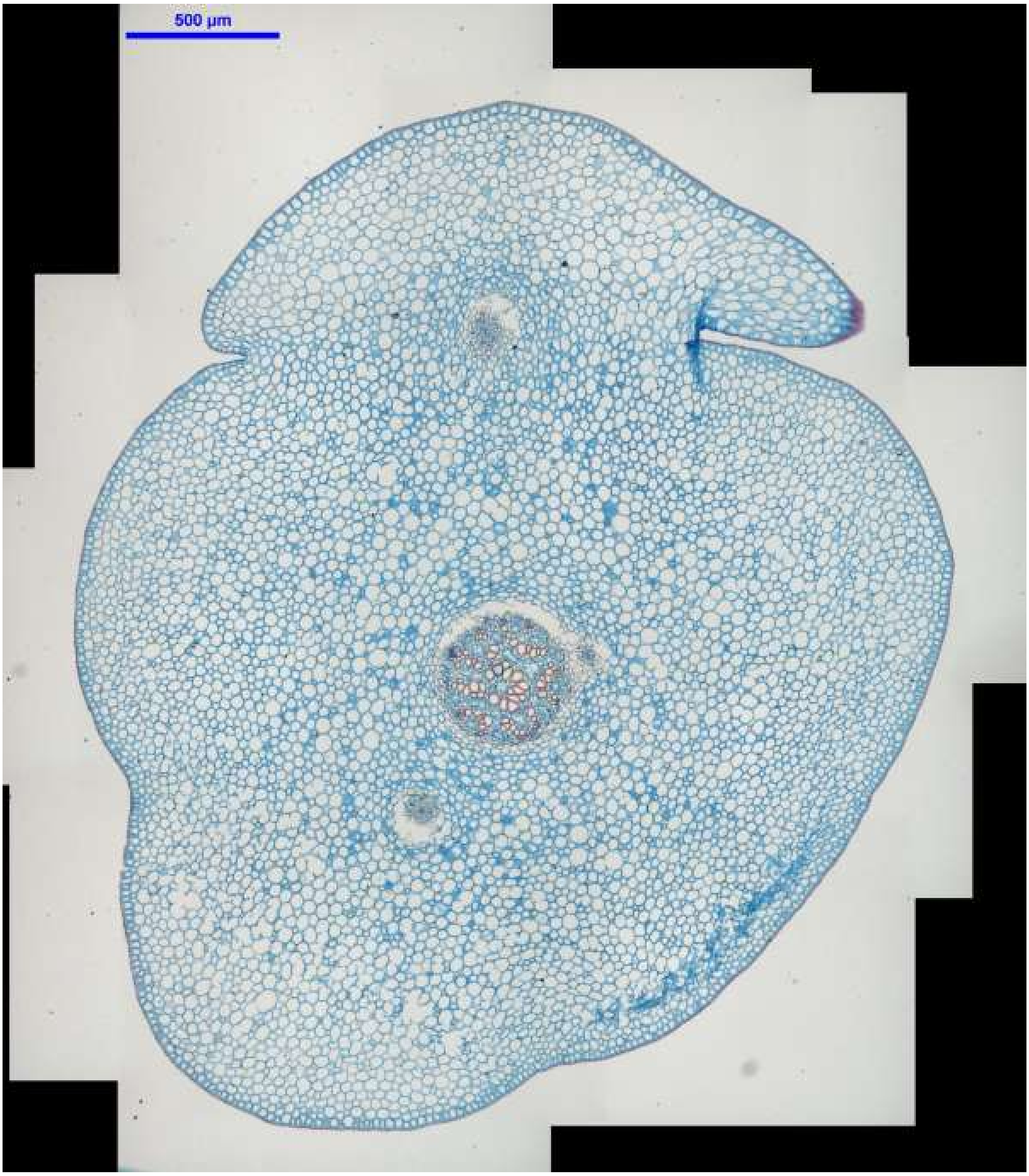
500 µm

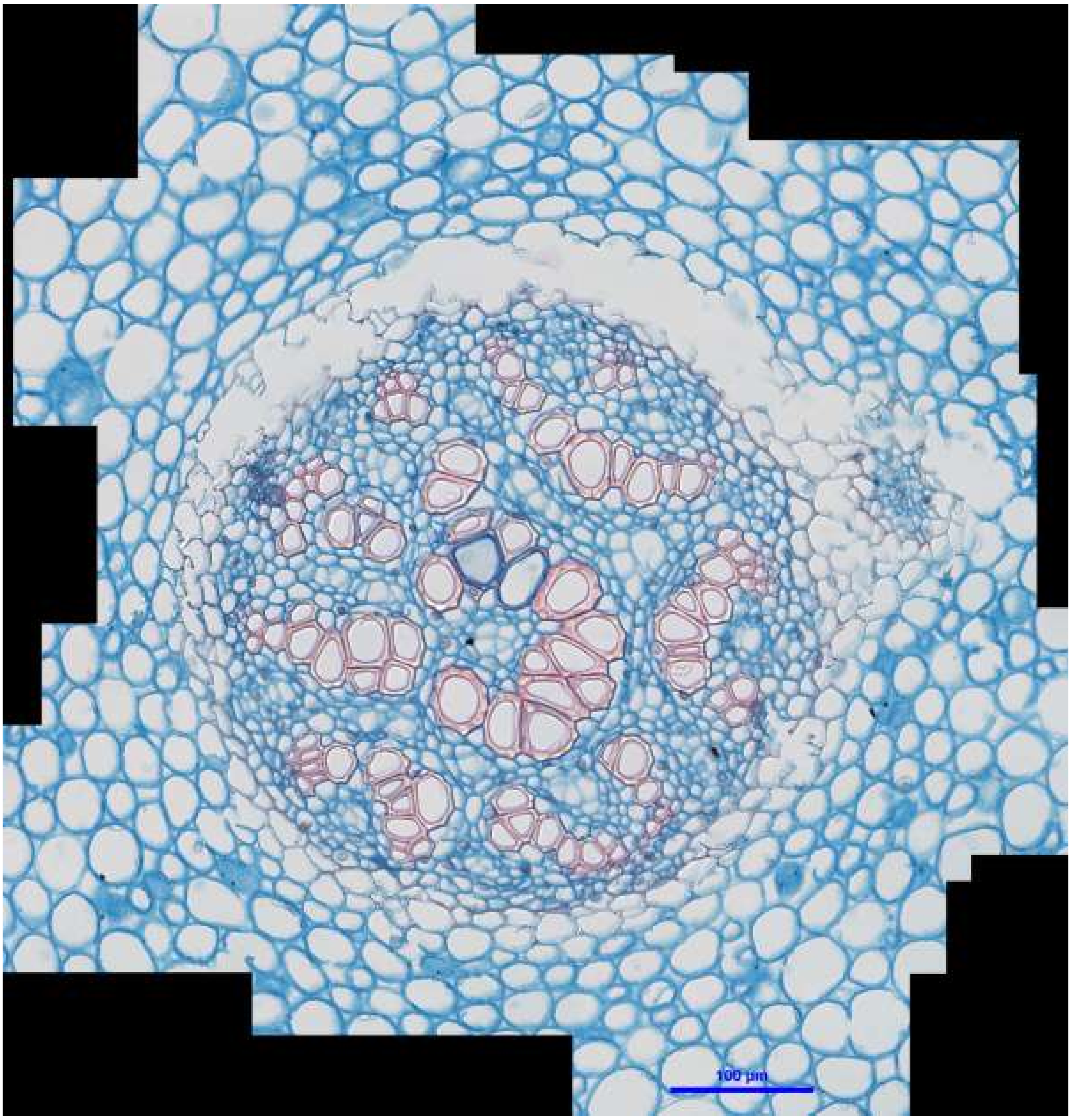


*Huperzia goebelii* stem: H_goeb_3

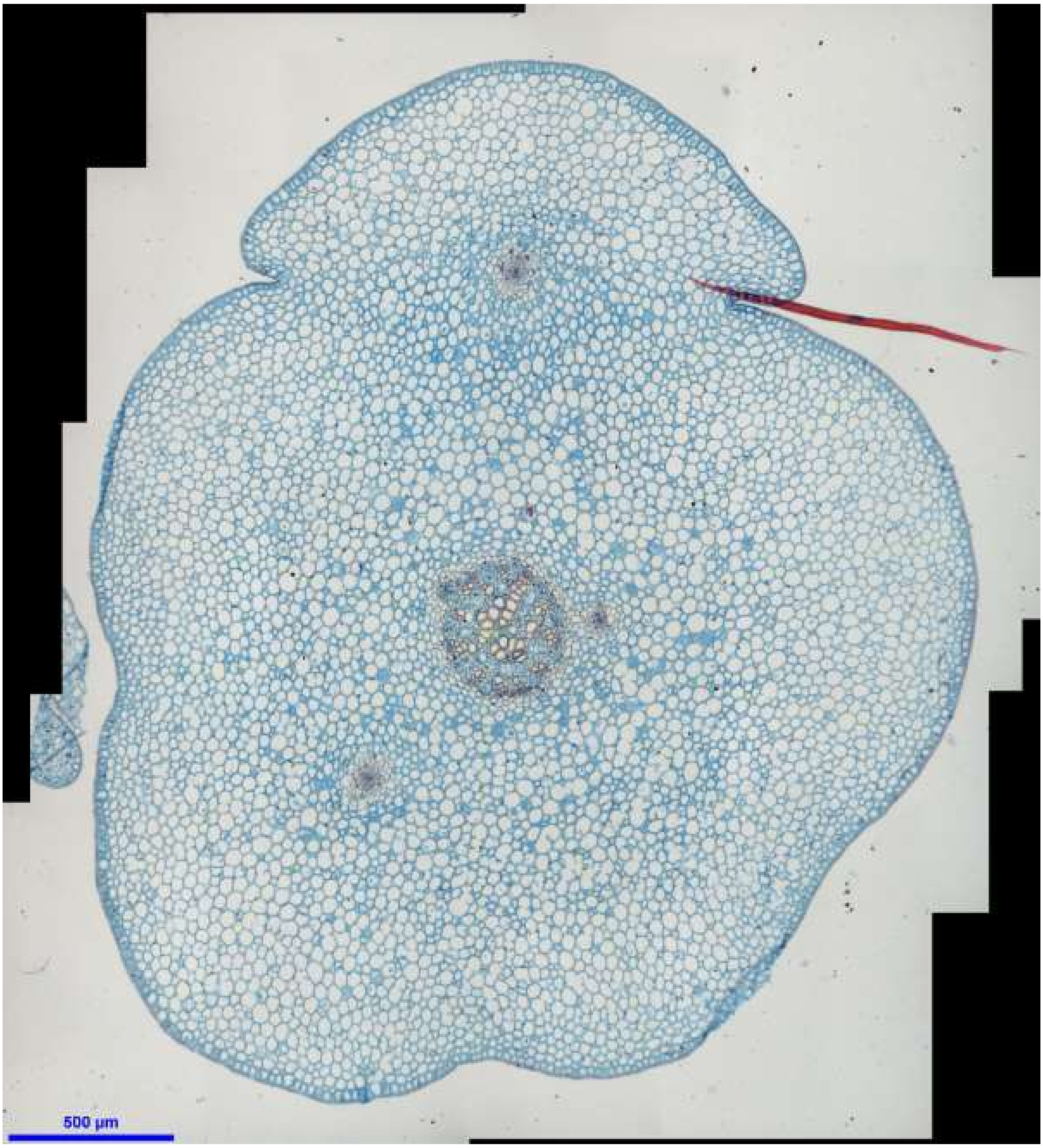
500 µm

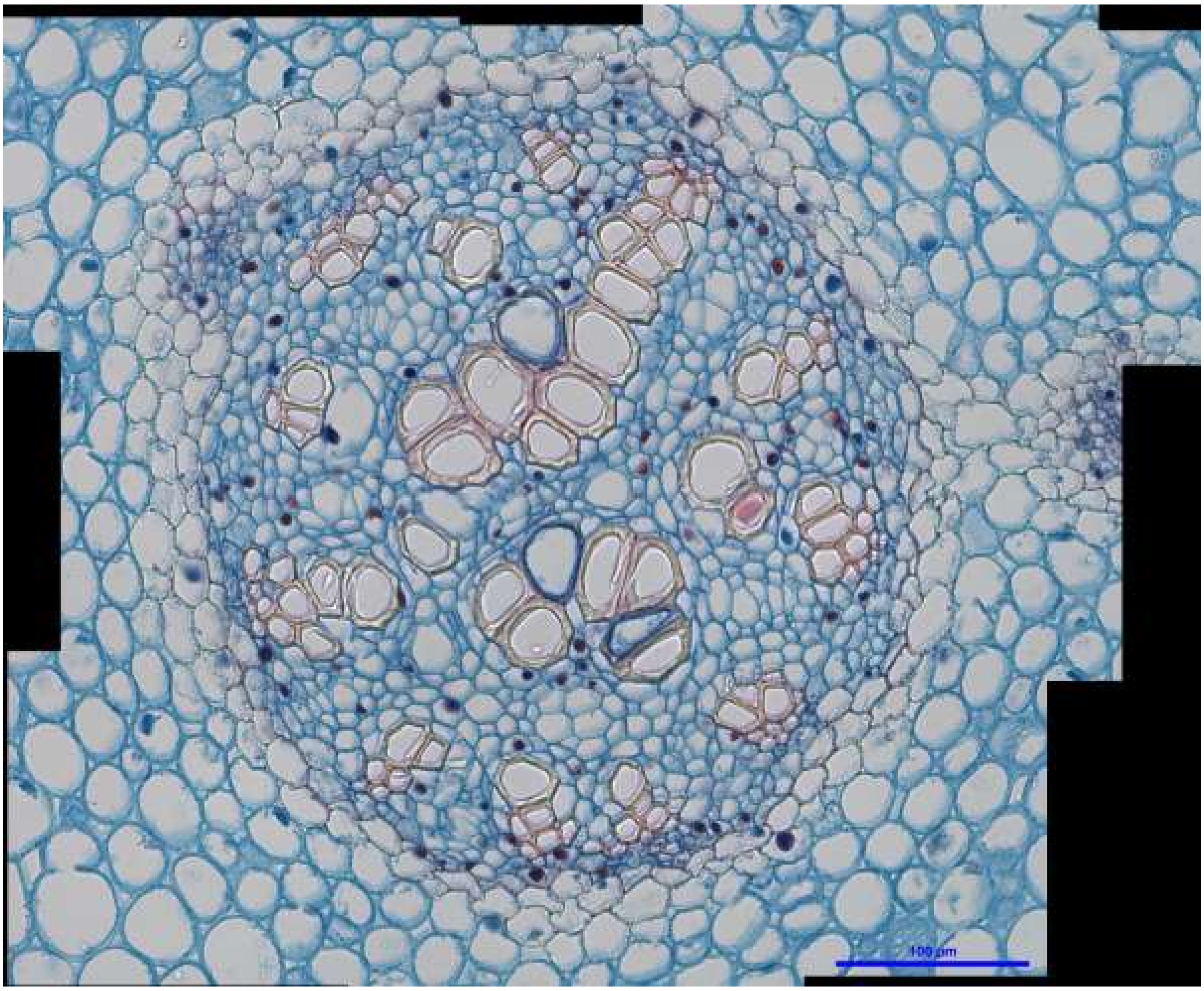


*Huperzia goebelii* stem: H_goeb_4

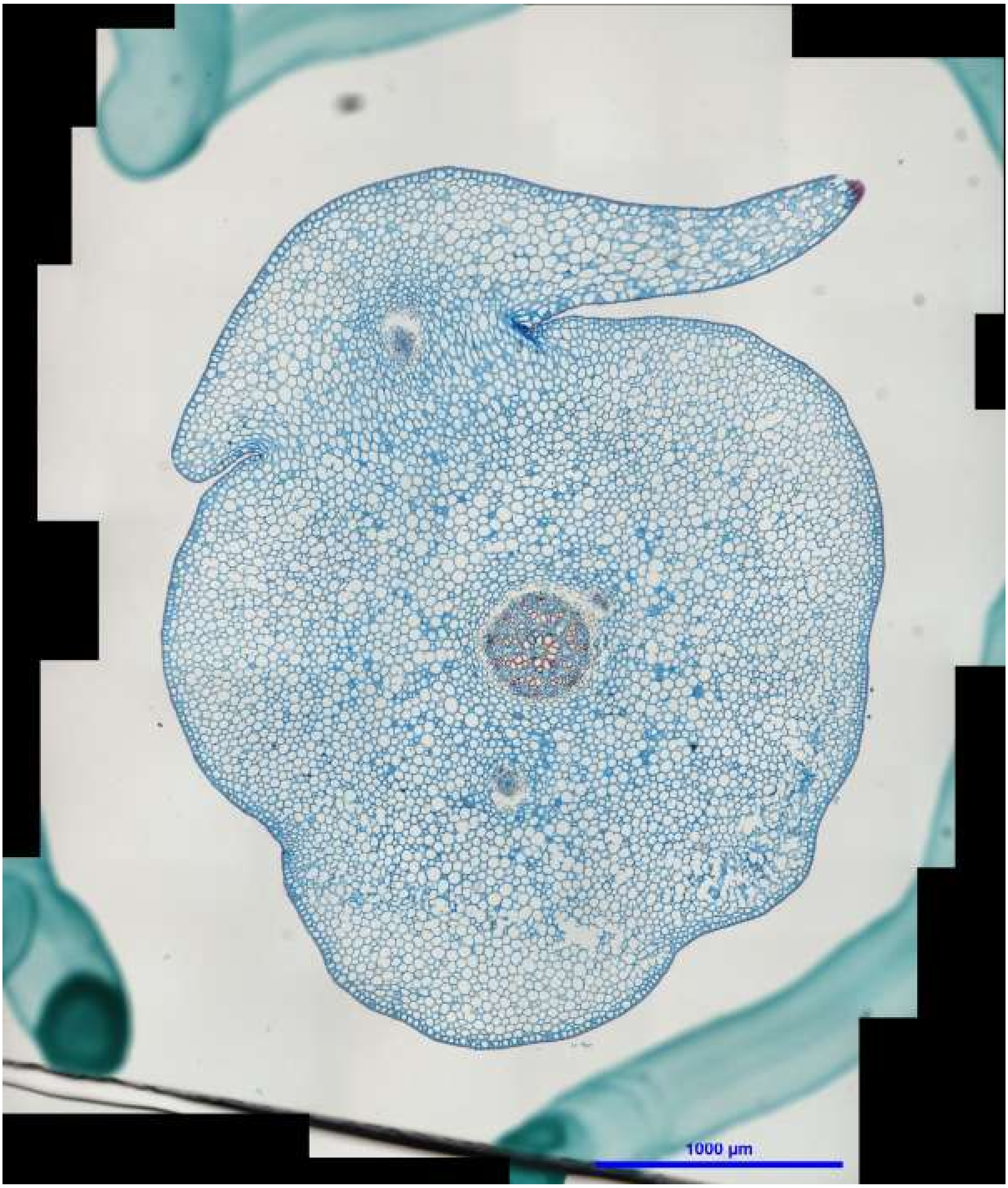
1000 µm

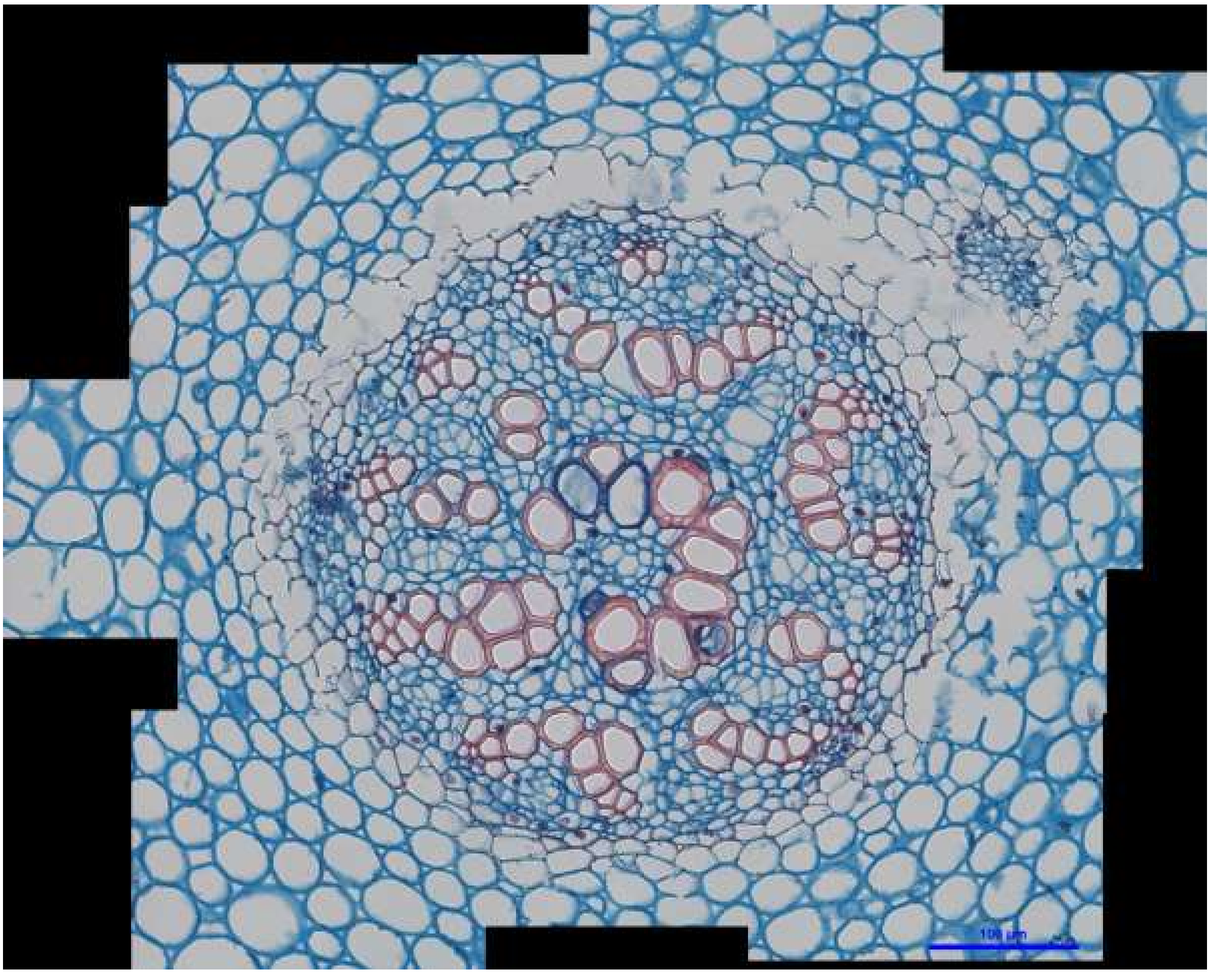


*Huperzia goebelii* stem: H_goeb_5

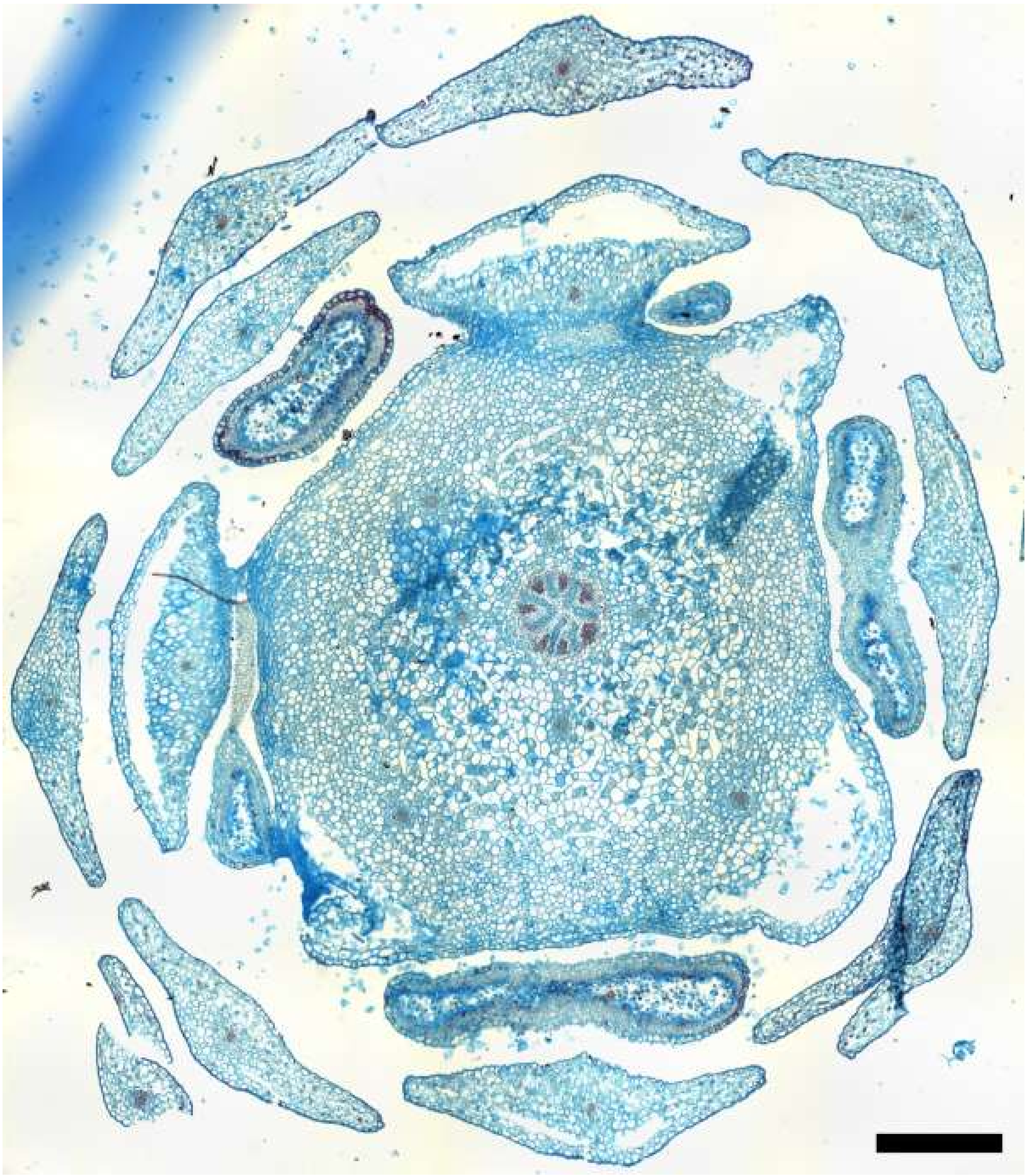

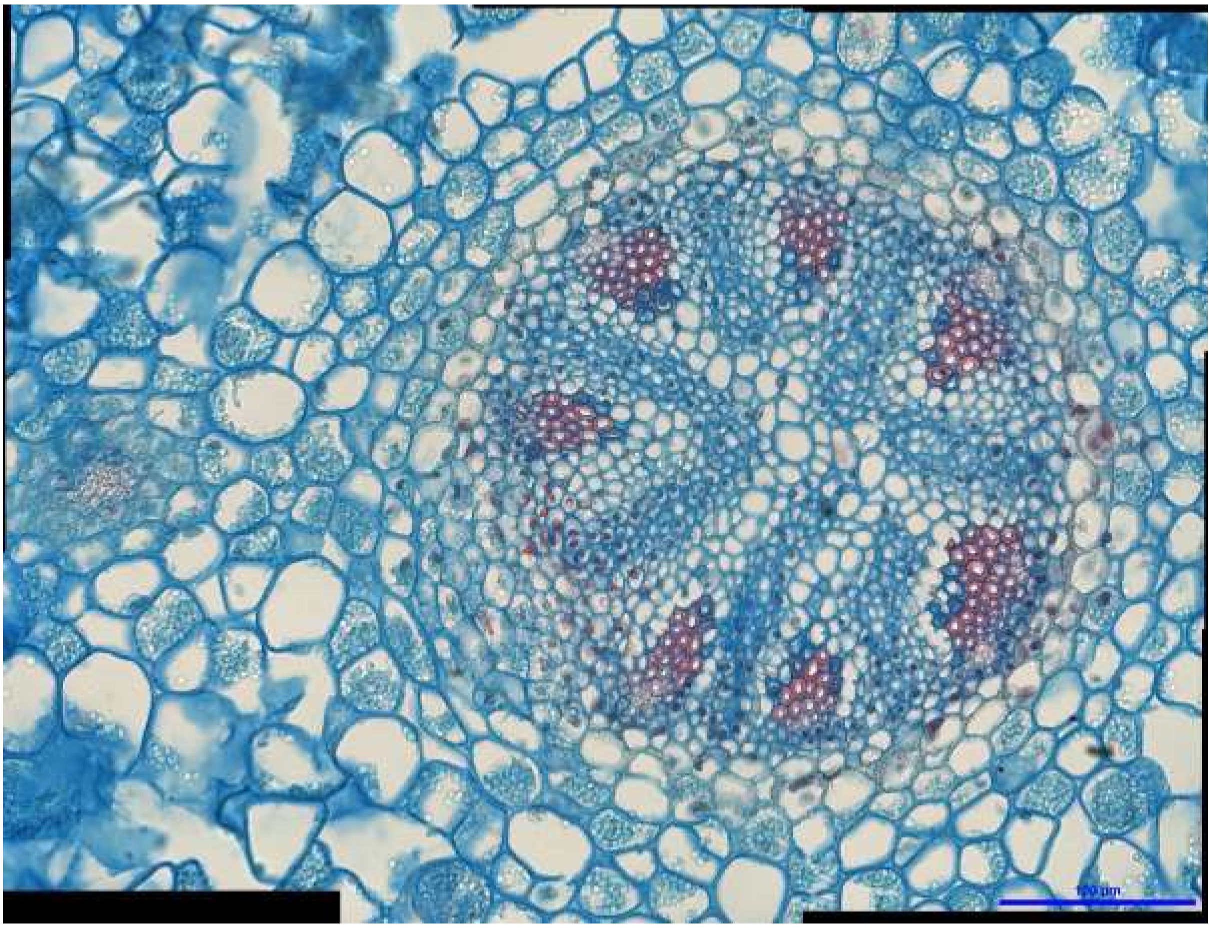

*Huperzia selago* stem: H_sel_1

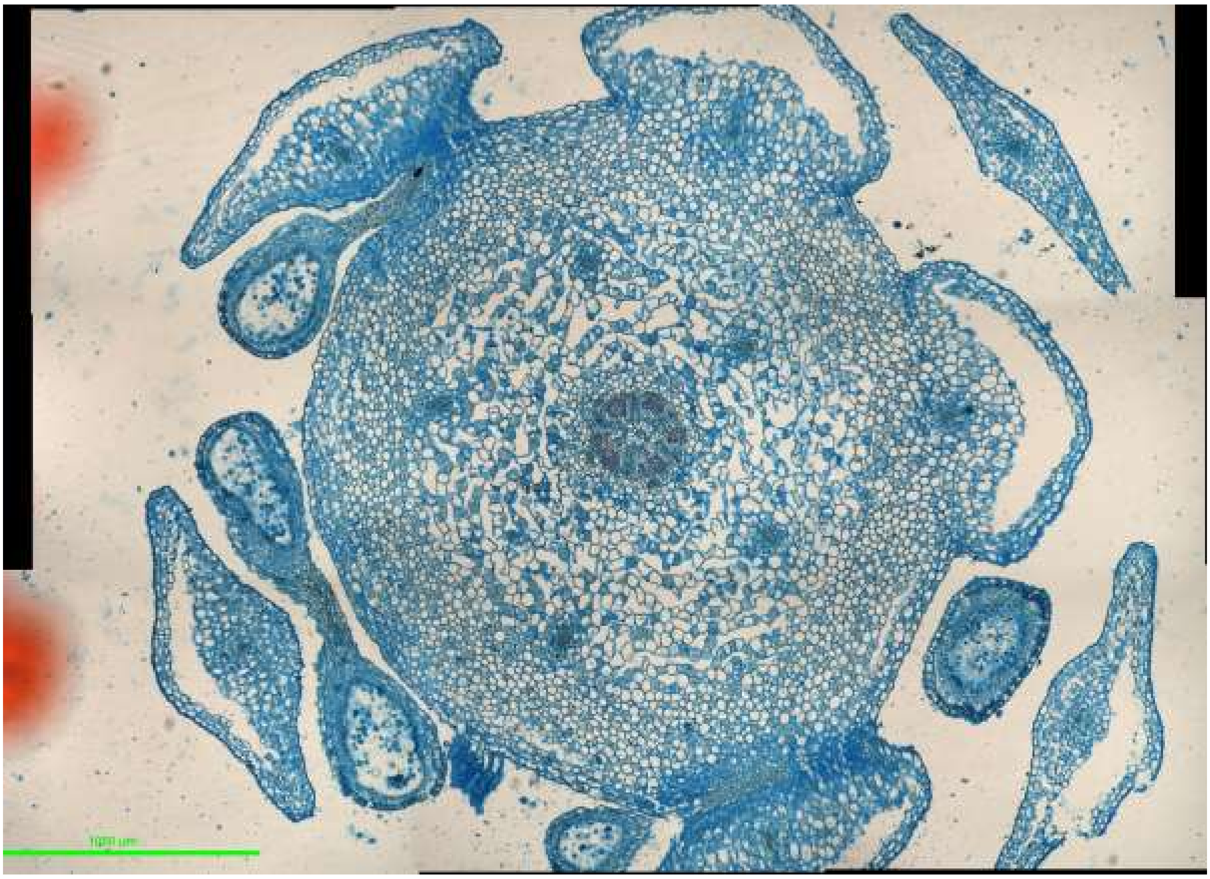

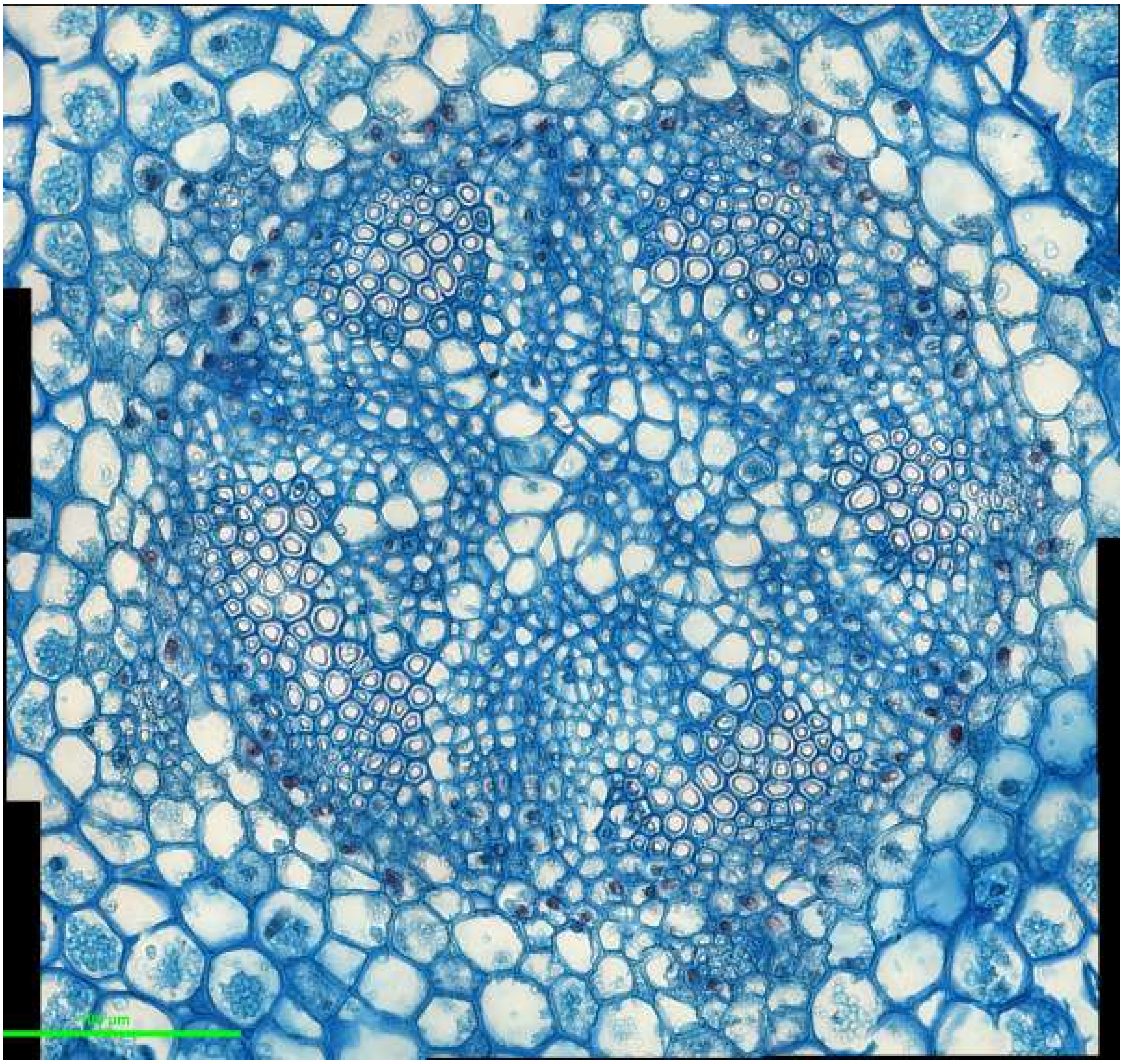

*Huperzia selago* stem: H_sel_2

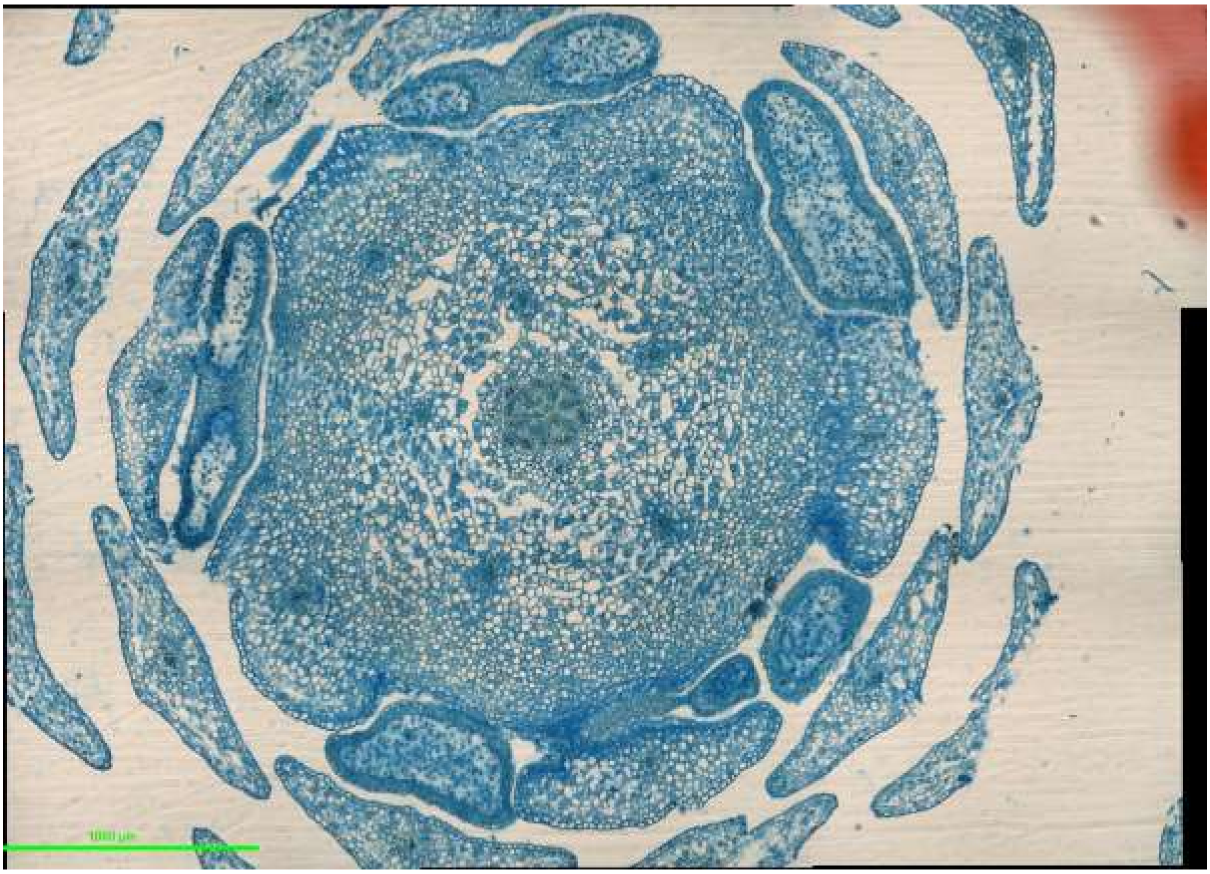


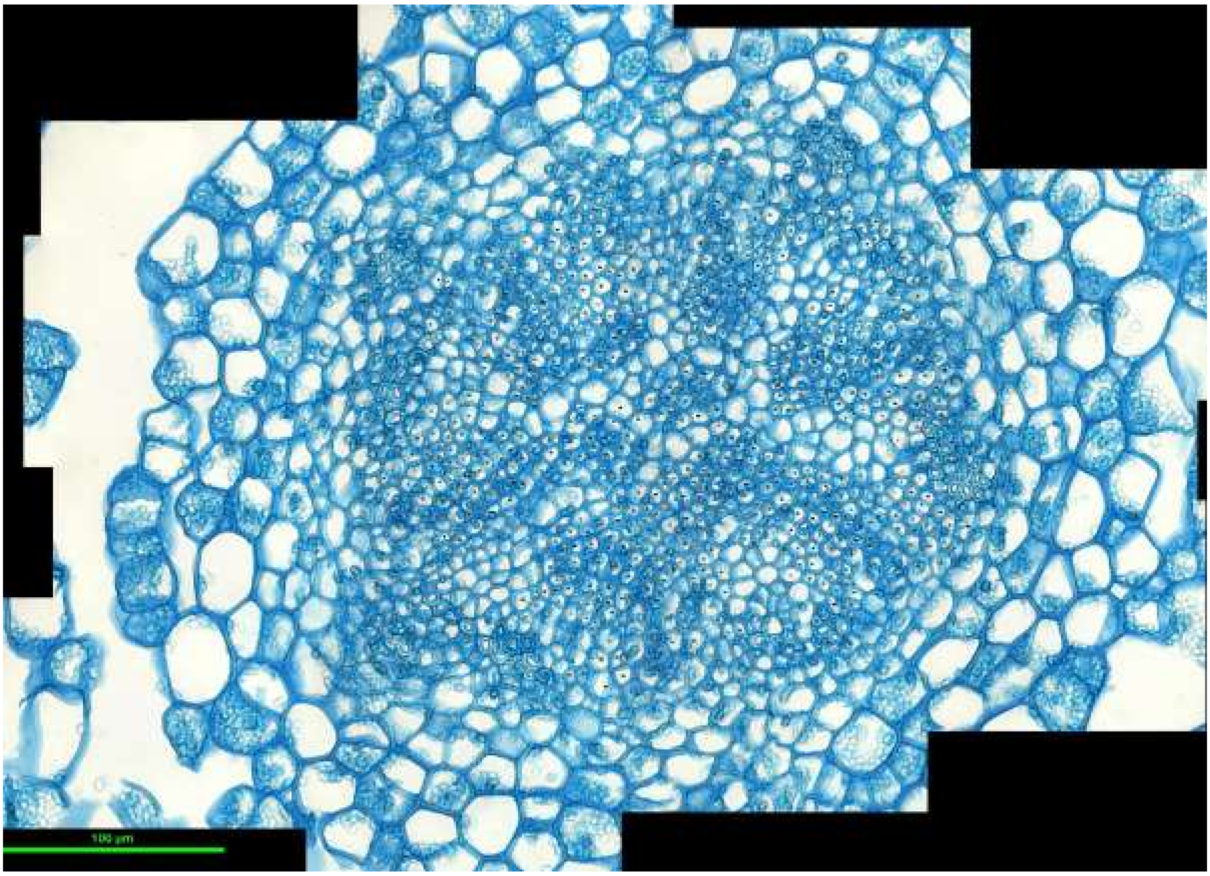


*Huperzia selago* stem: H_sel_3

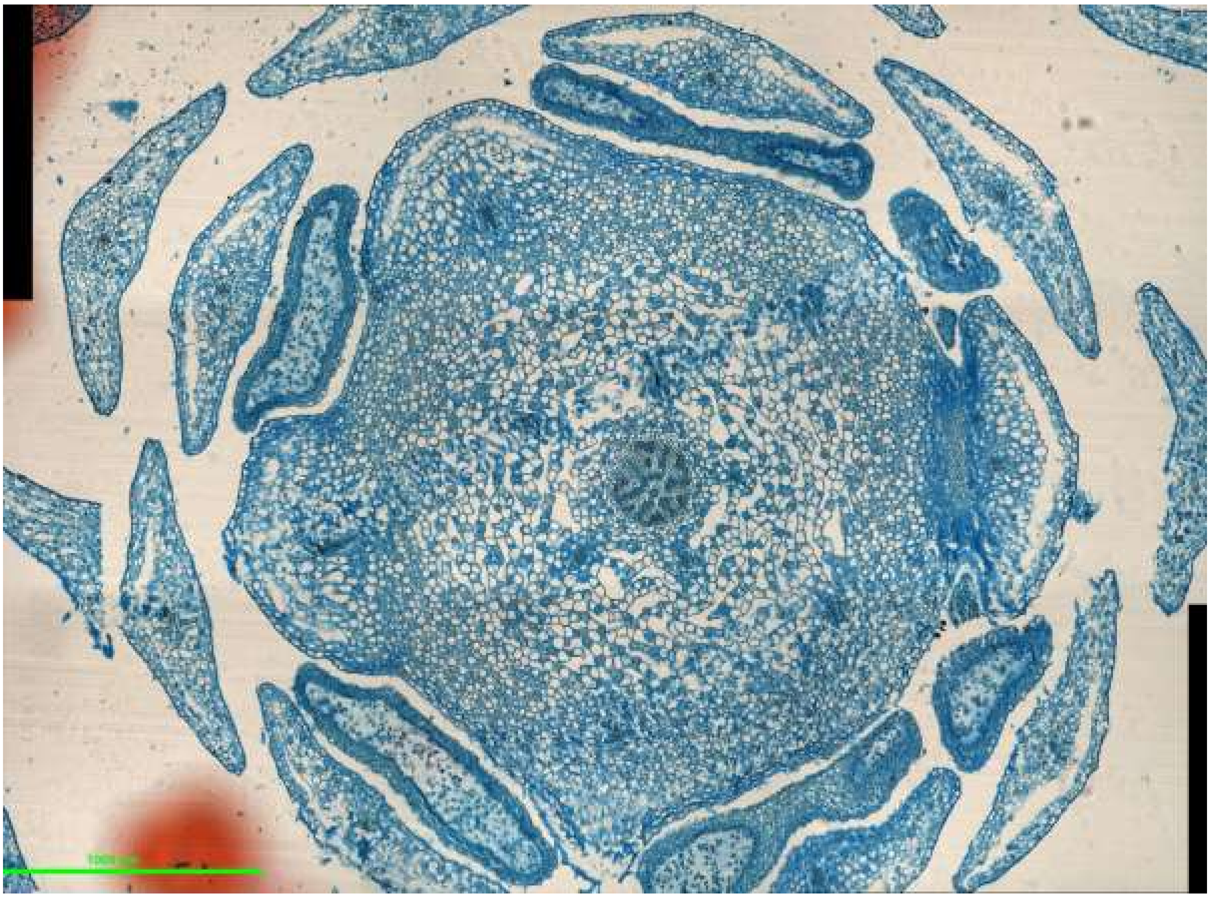

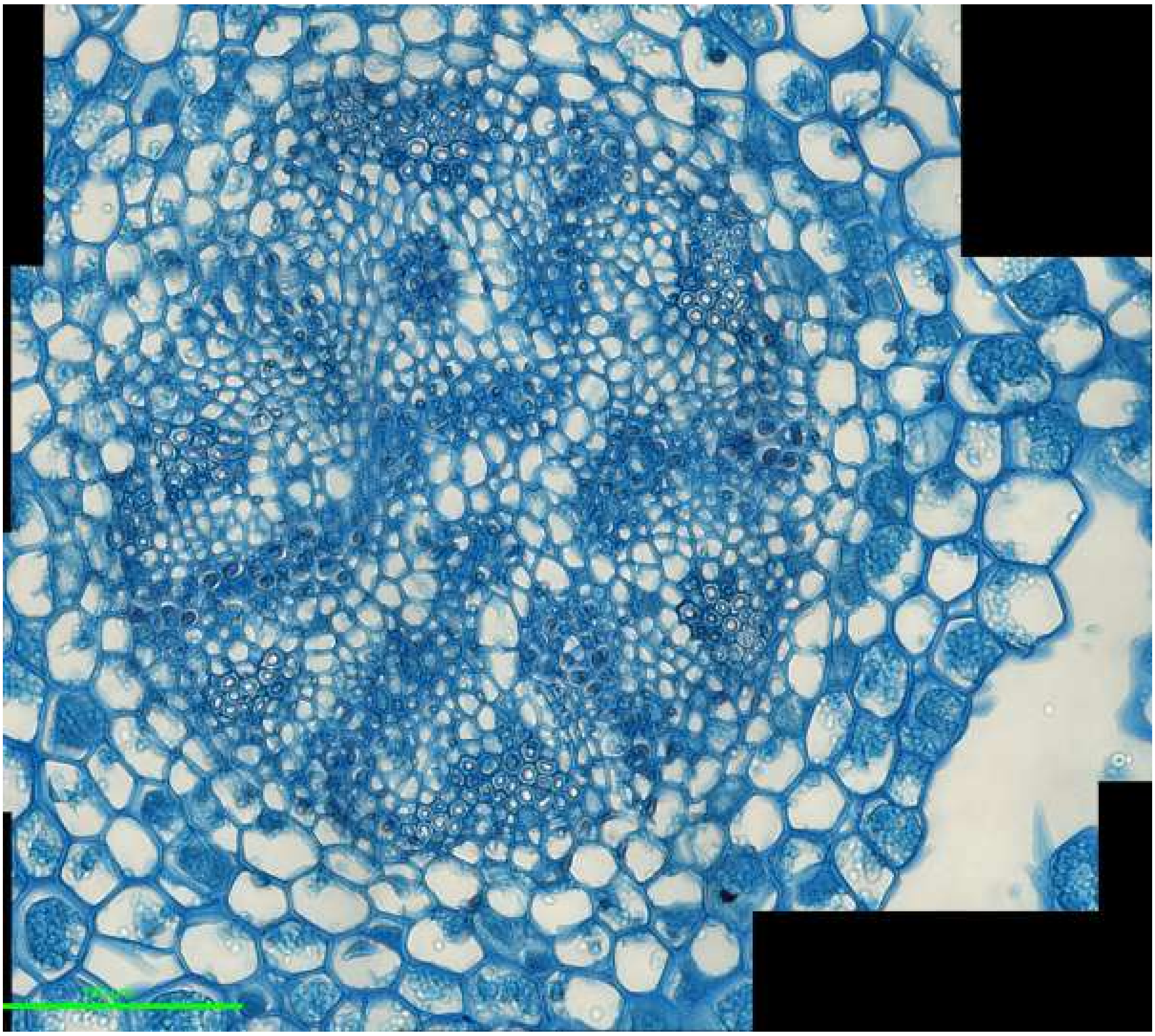

*Huperzia selago* stem: H_sel_4

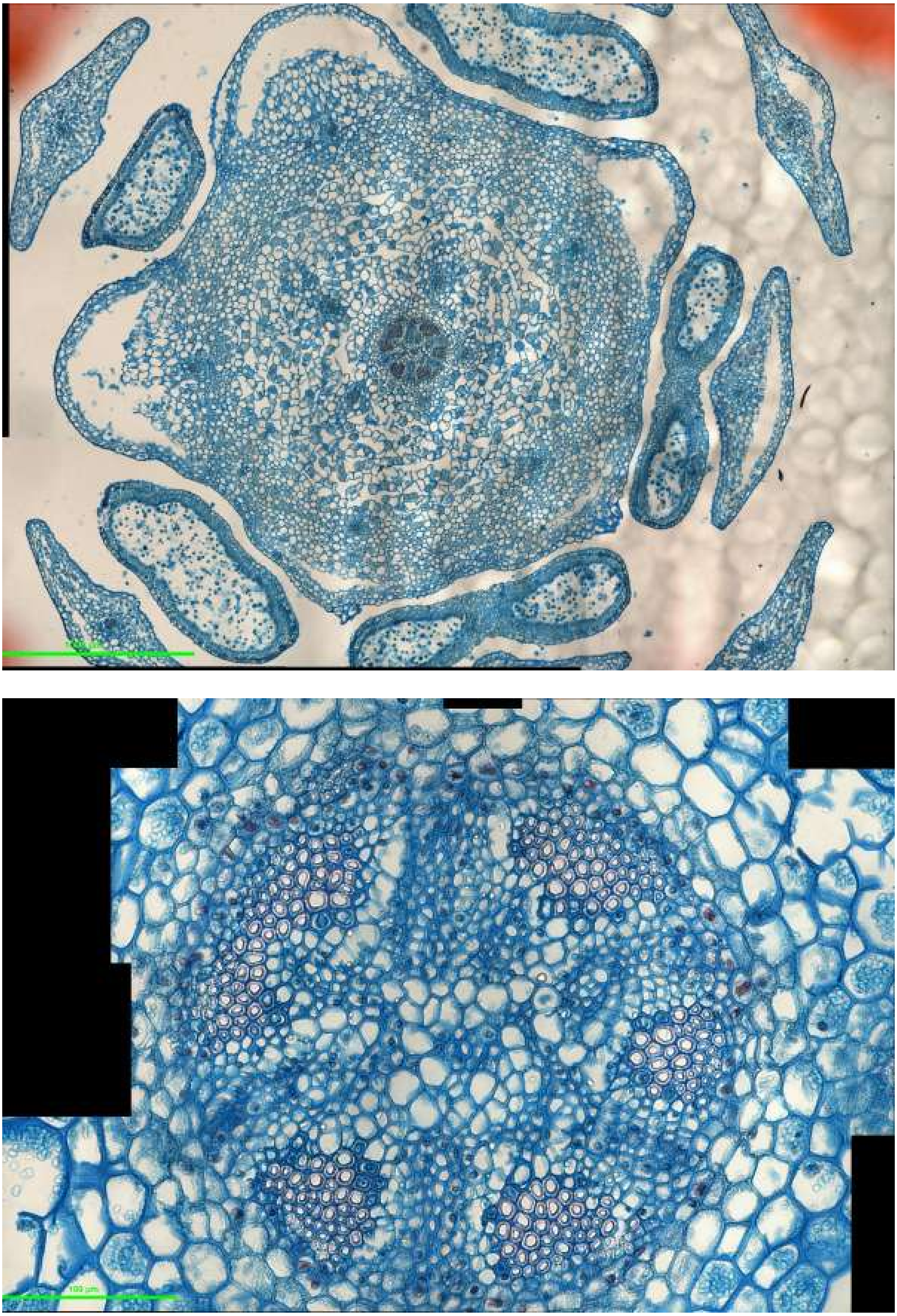

*Huperzia selago* stem: H_sel_5

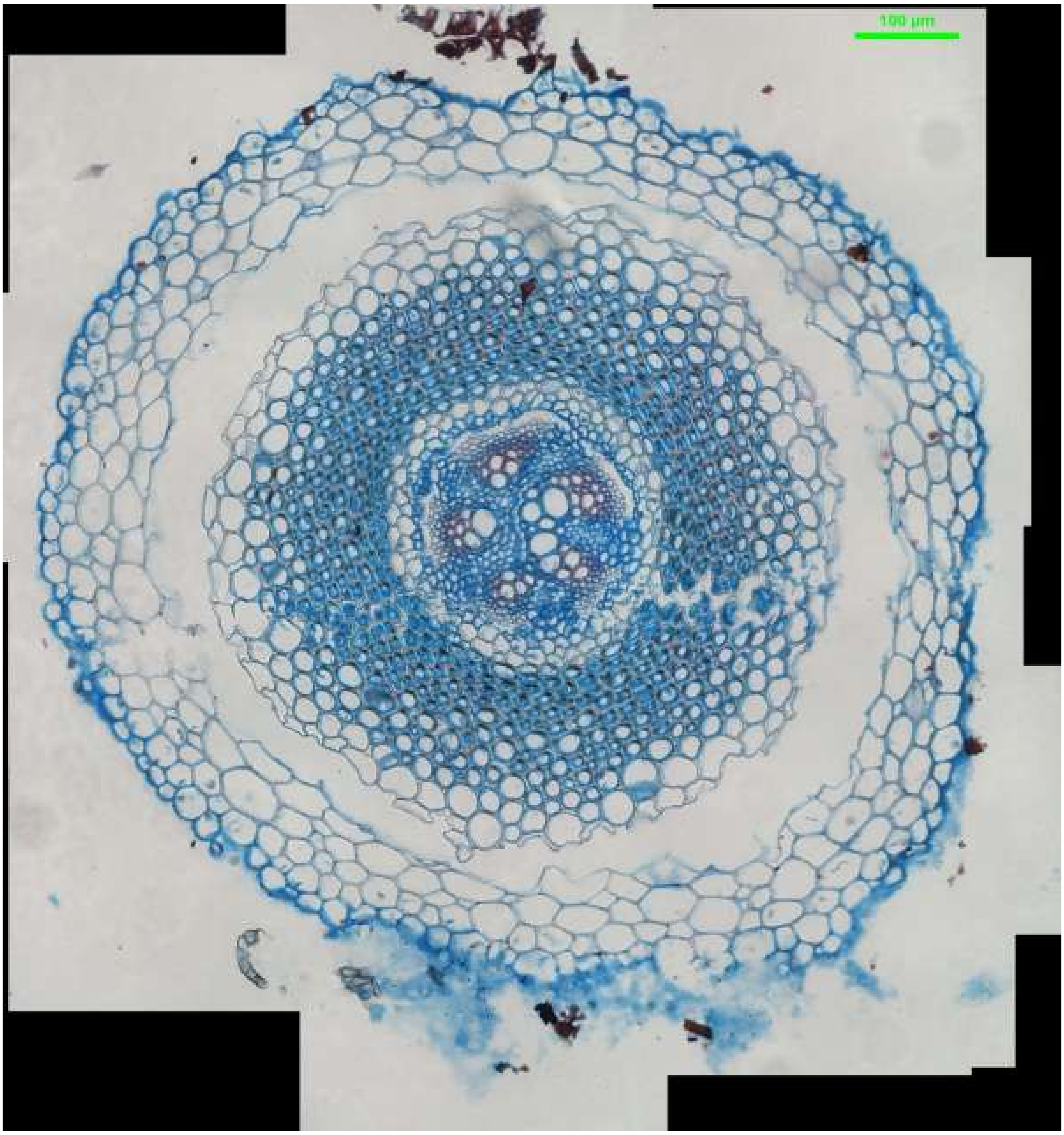
100 µm

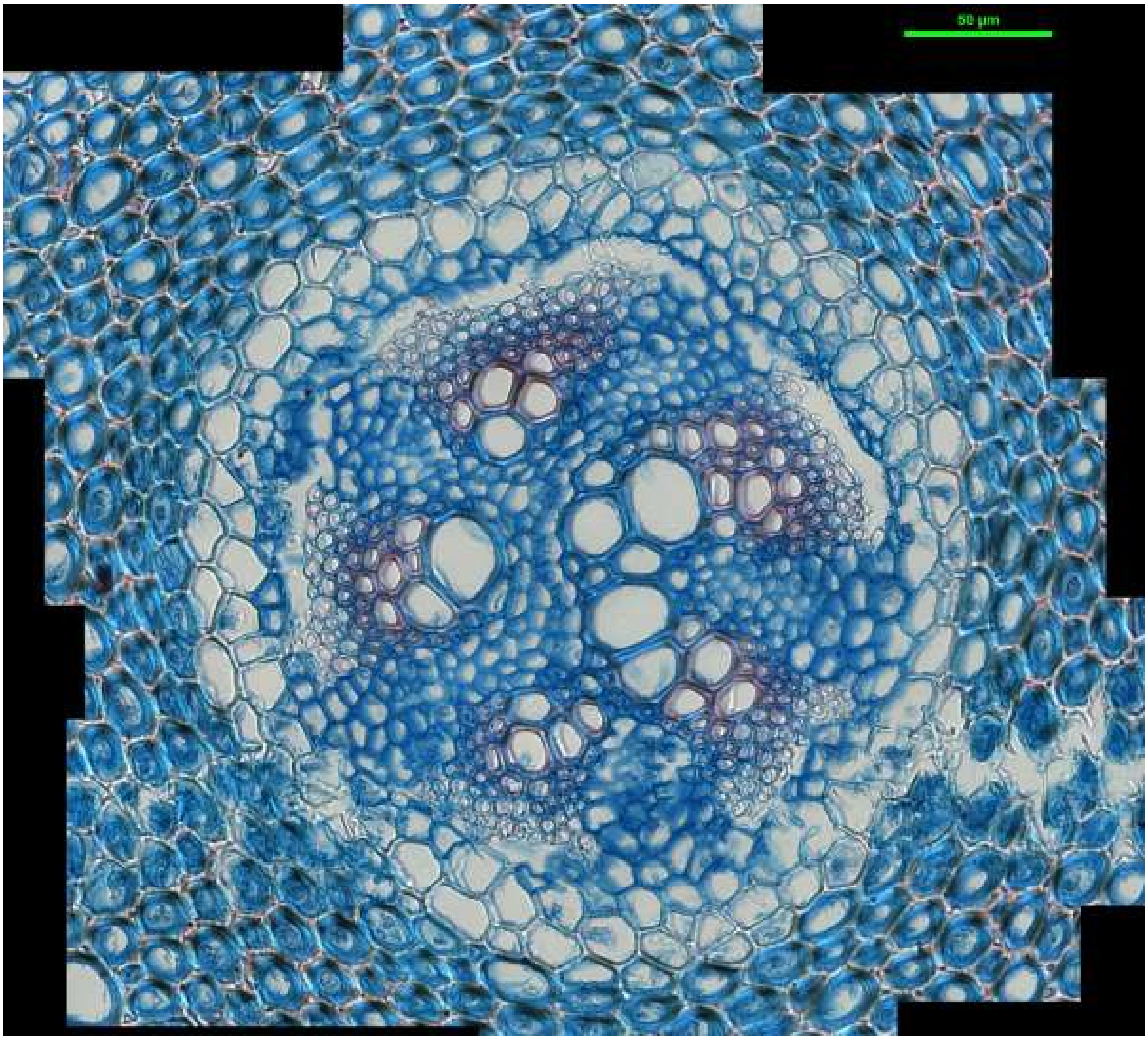


*Lycopodium clavatum* root: L_clav_root_1

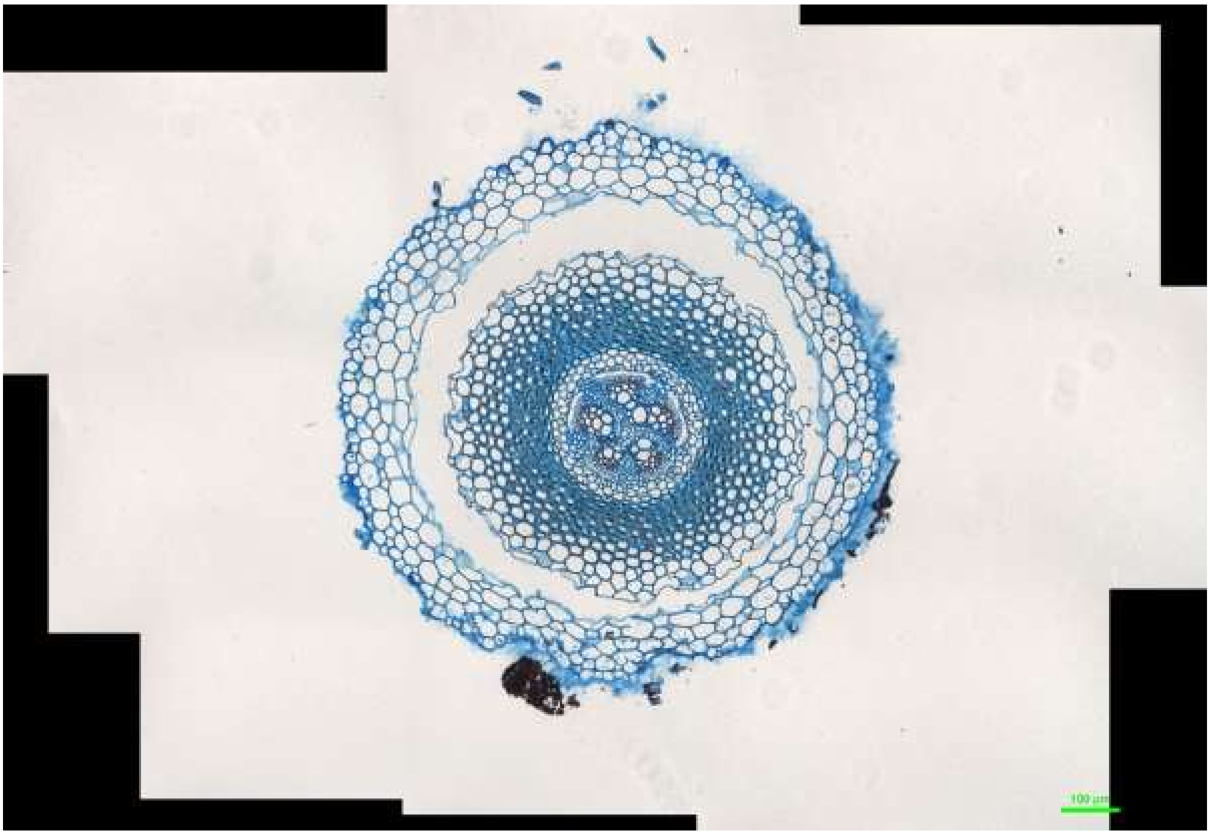


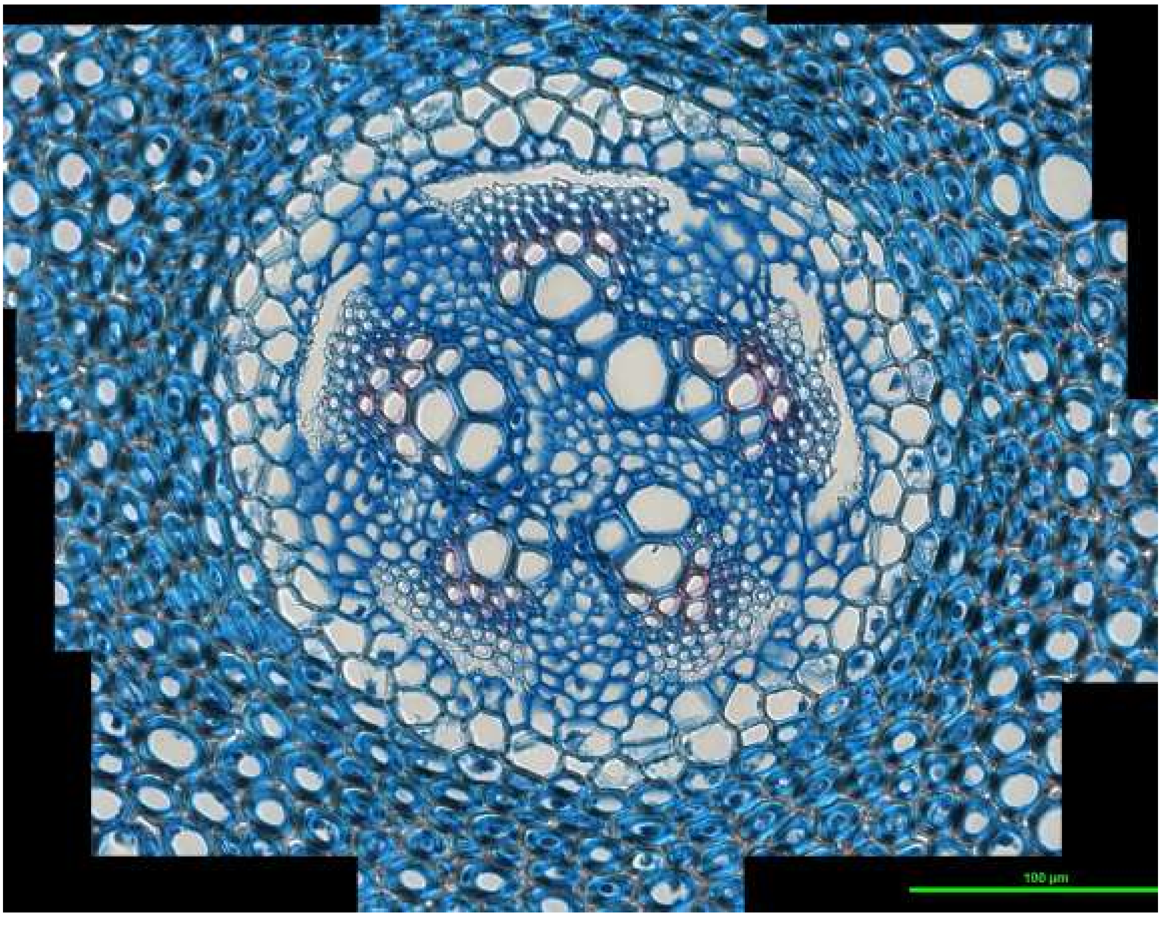


*Lycopodium clavatum* root: L_clav_root_2

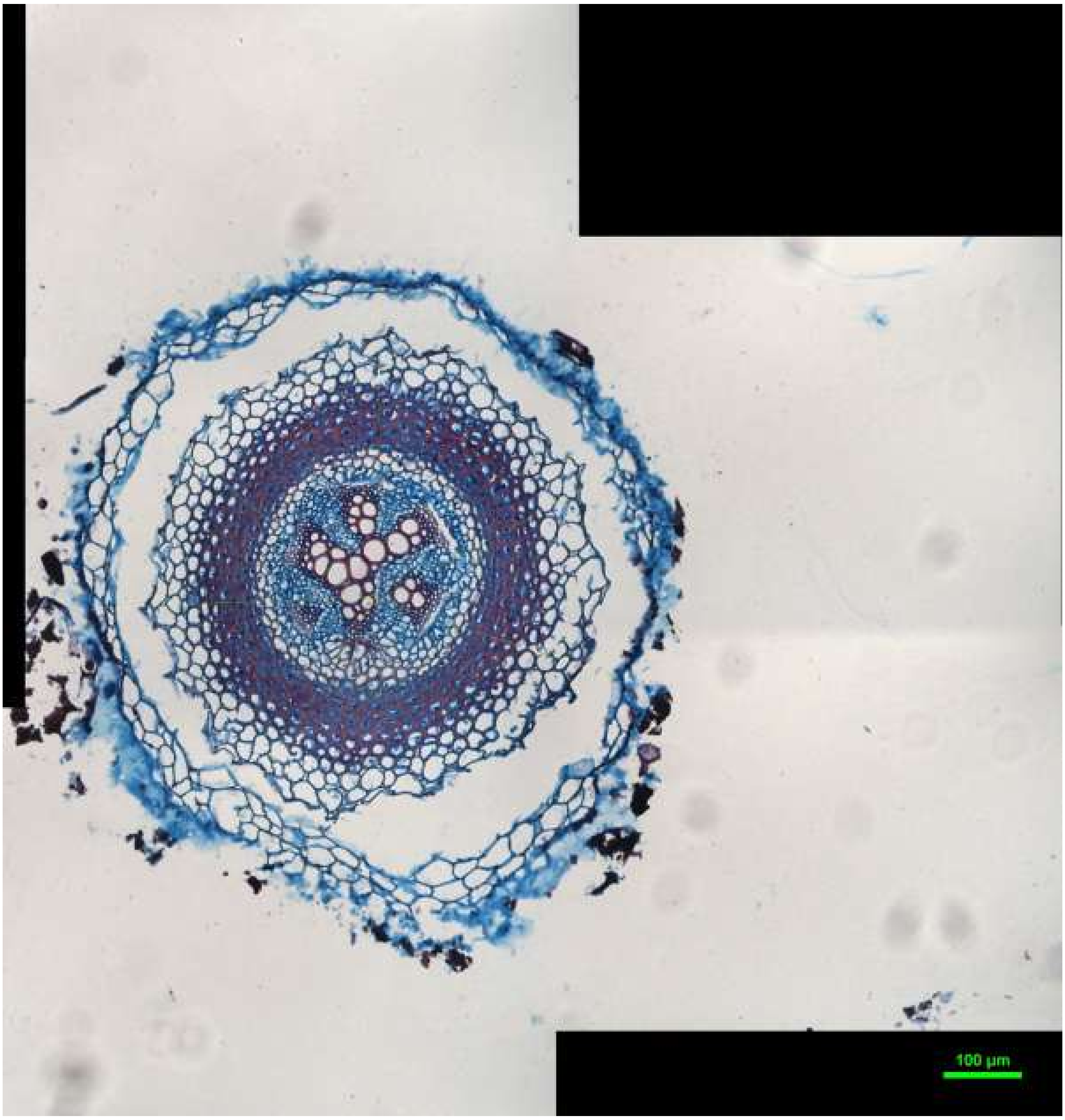
100 µm

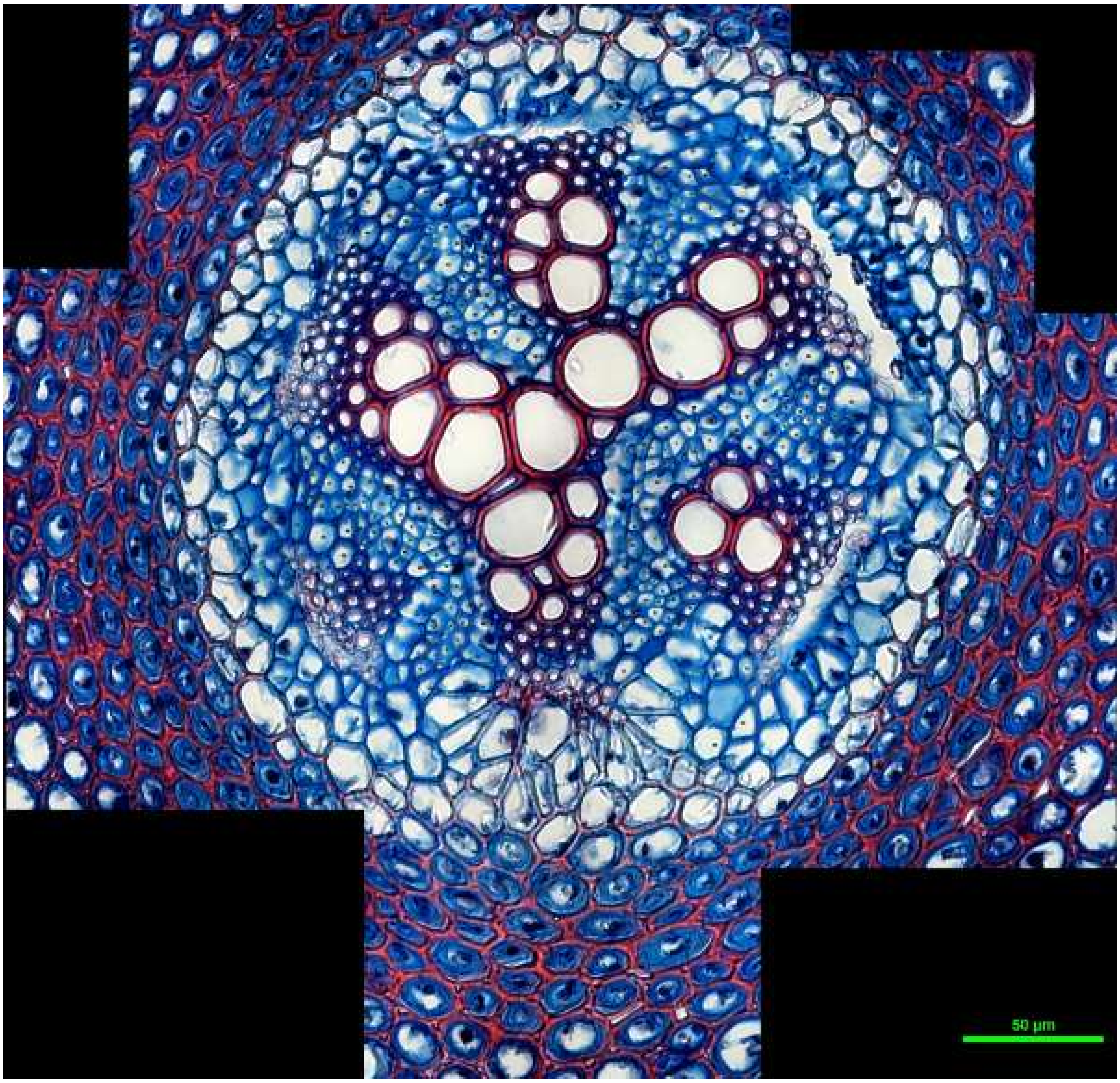


*Lycopodium clavatum* root: L_clav_root_3

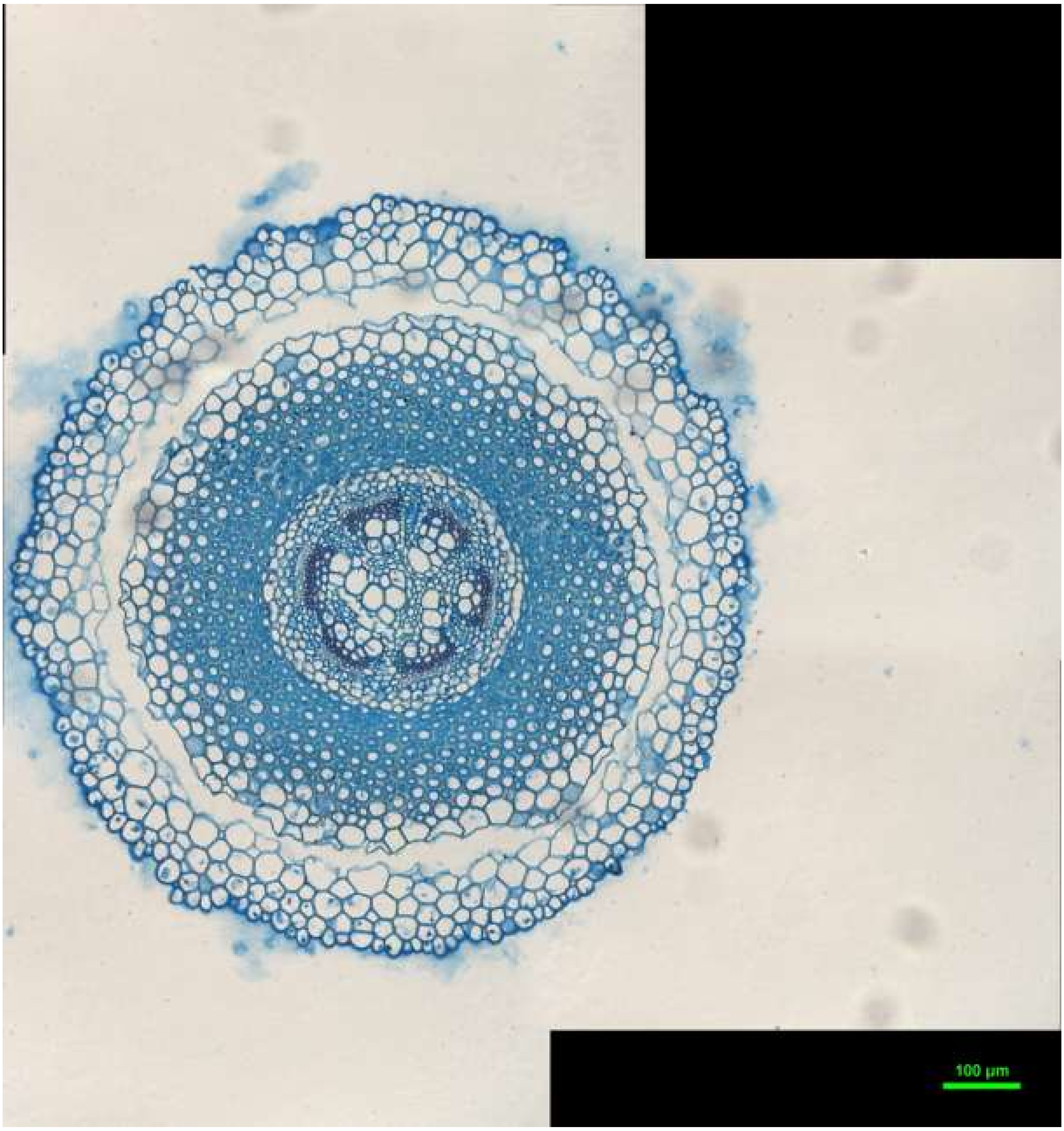
100 µm

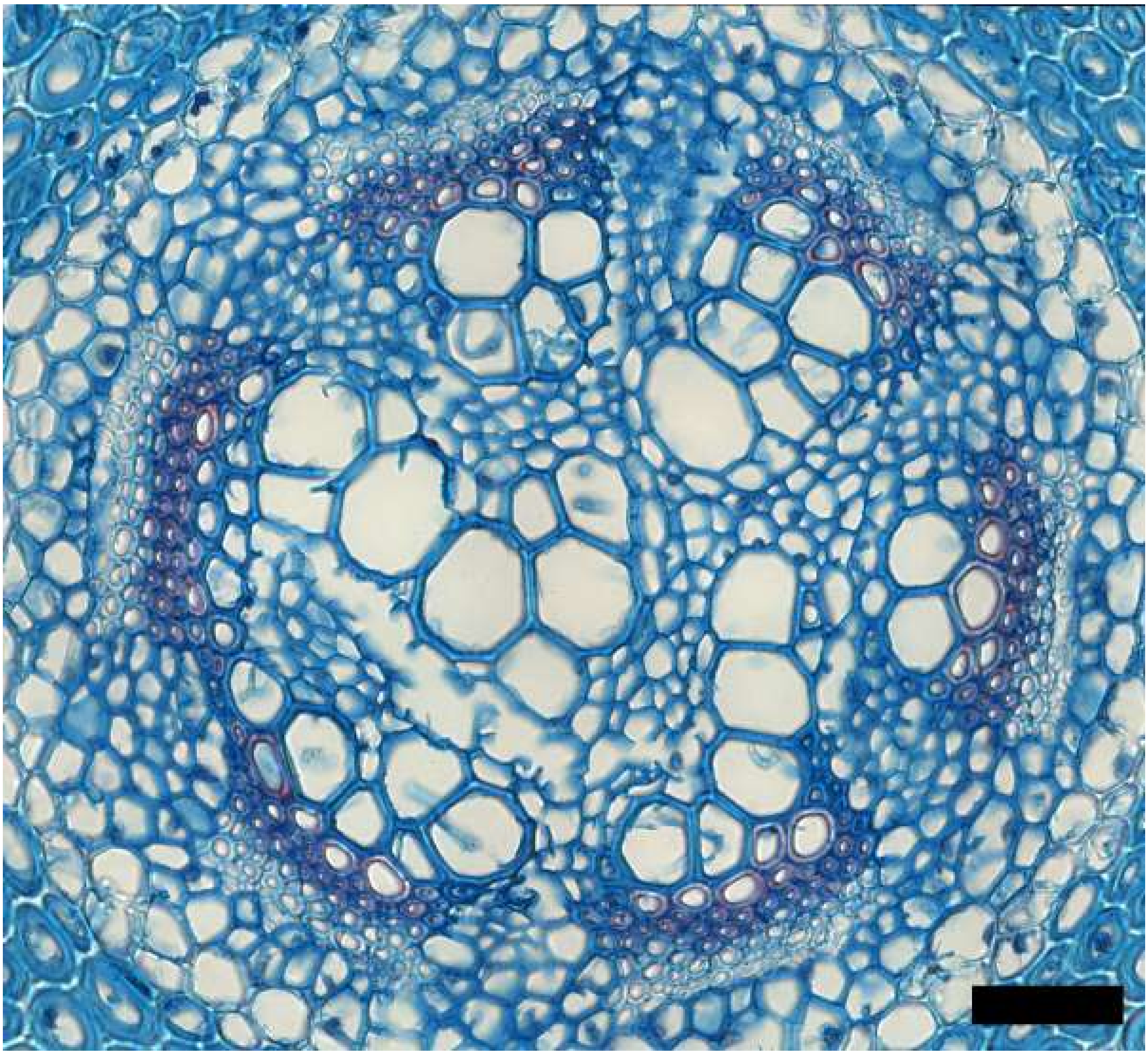

*Lycopodium clavatum* root: L_clav_root_4

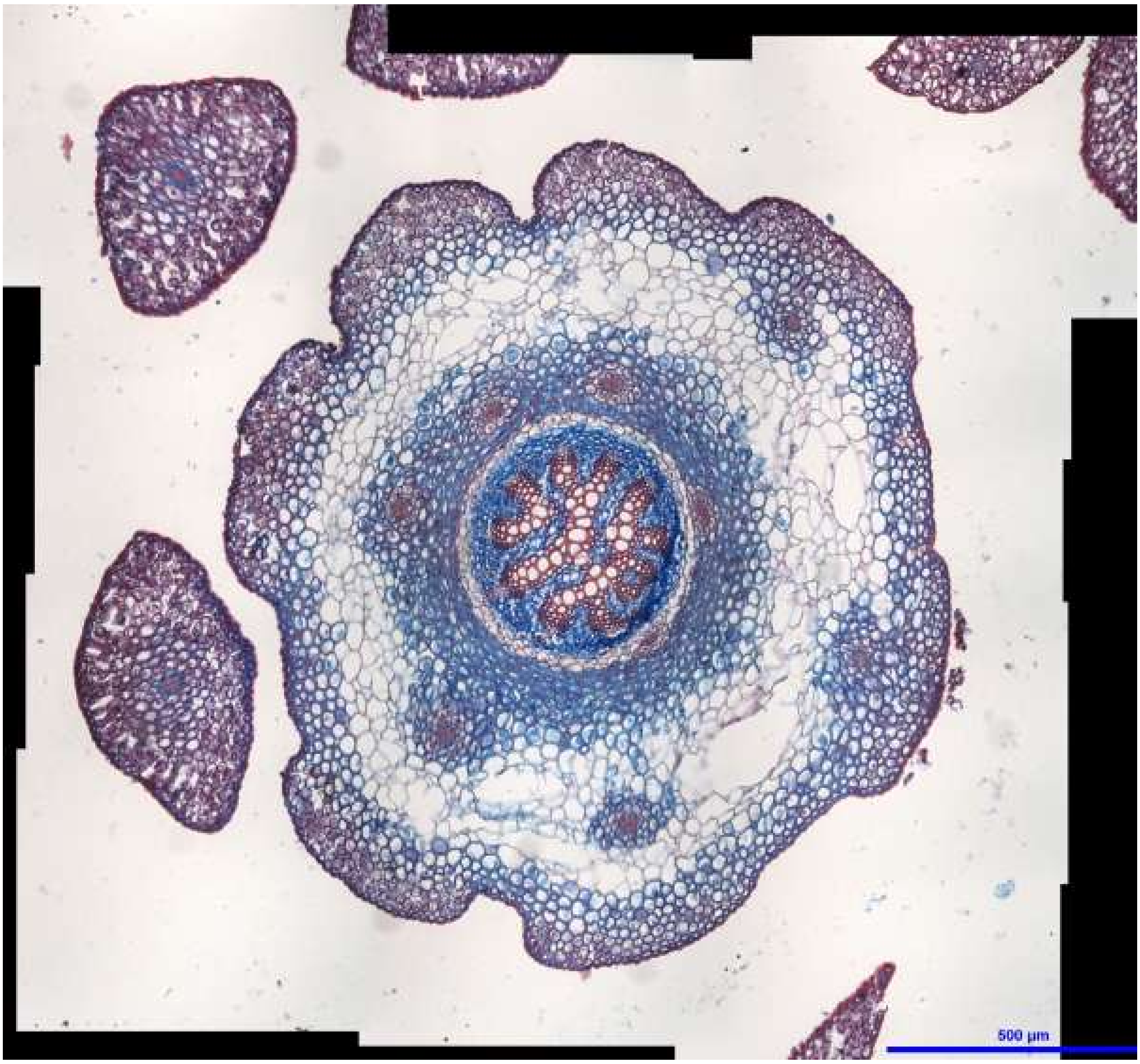
500 µm

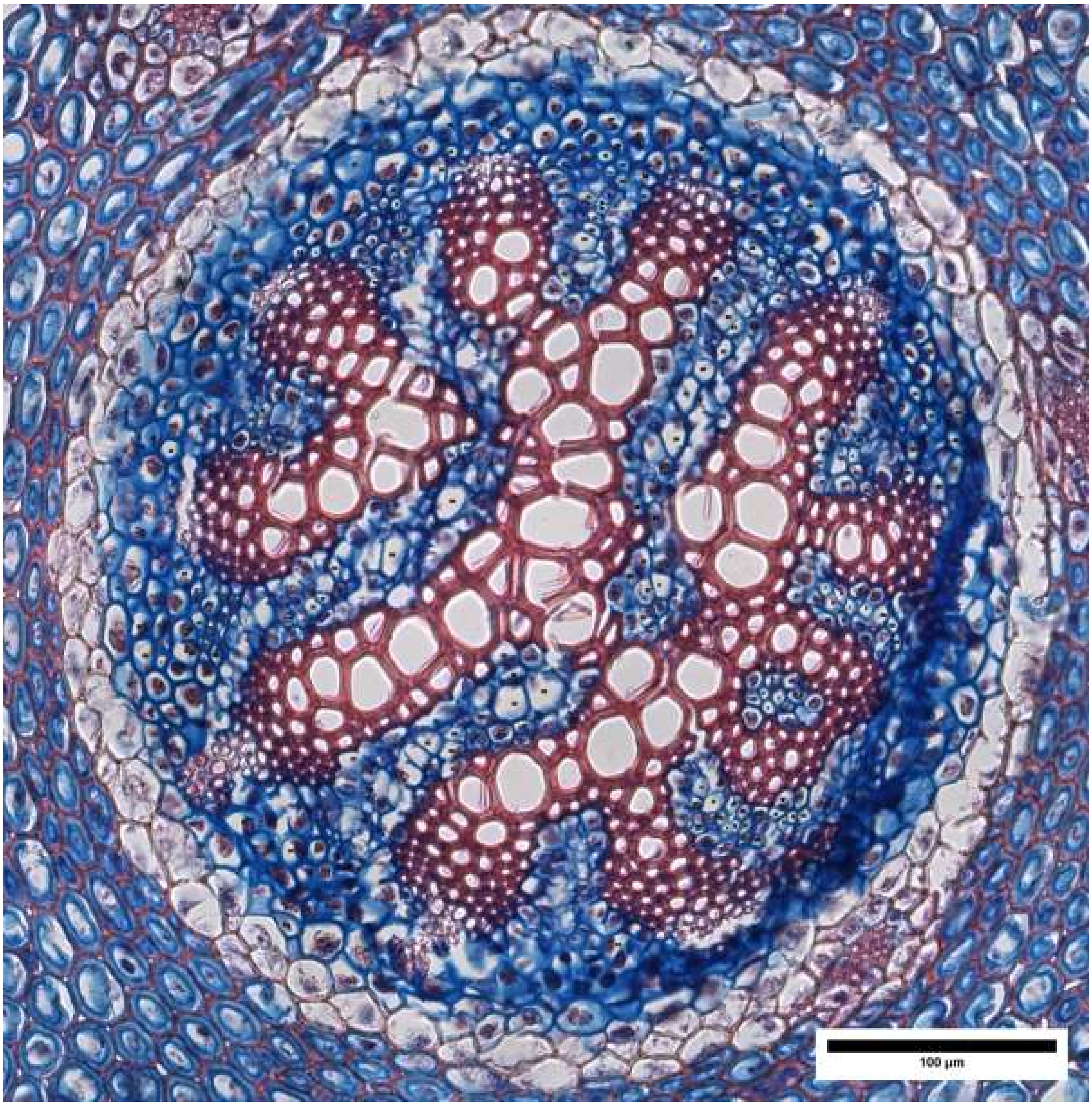
100 µm

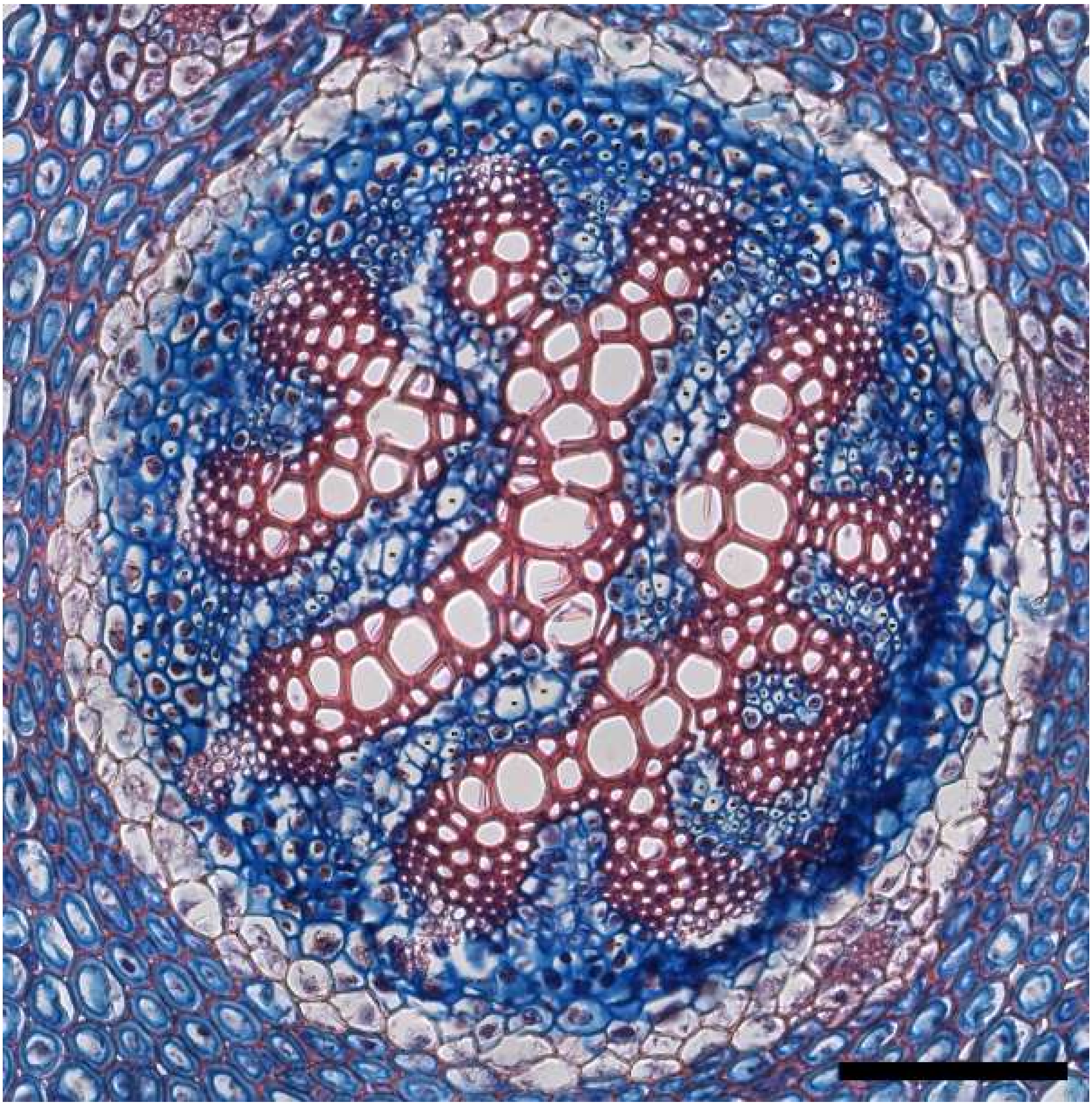

*Lycopodium clavatum* stem: L_clav_stem_1

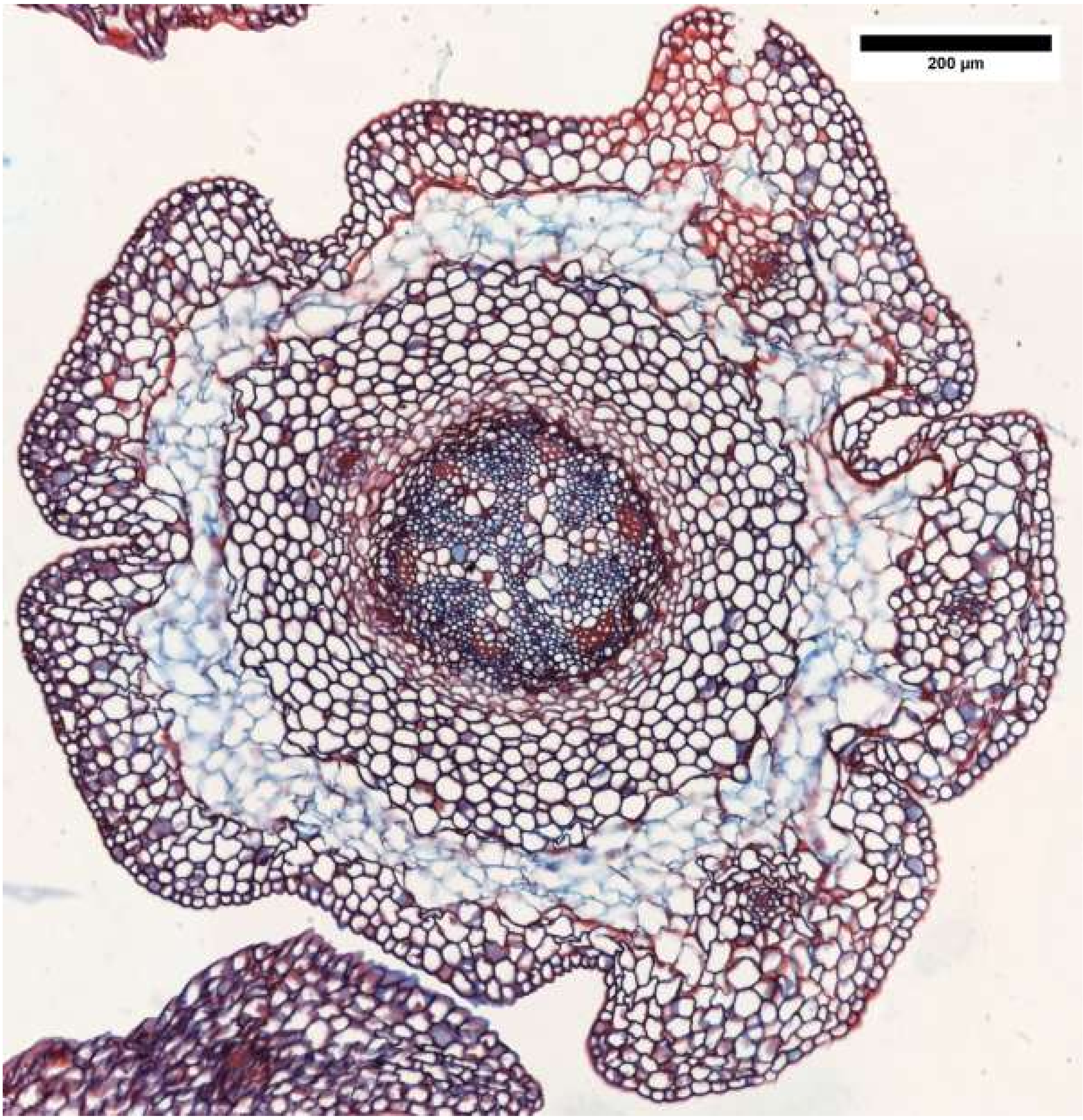
200 µm

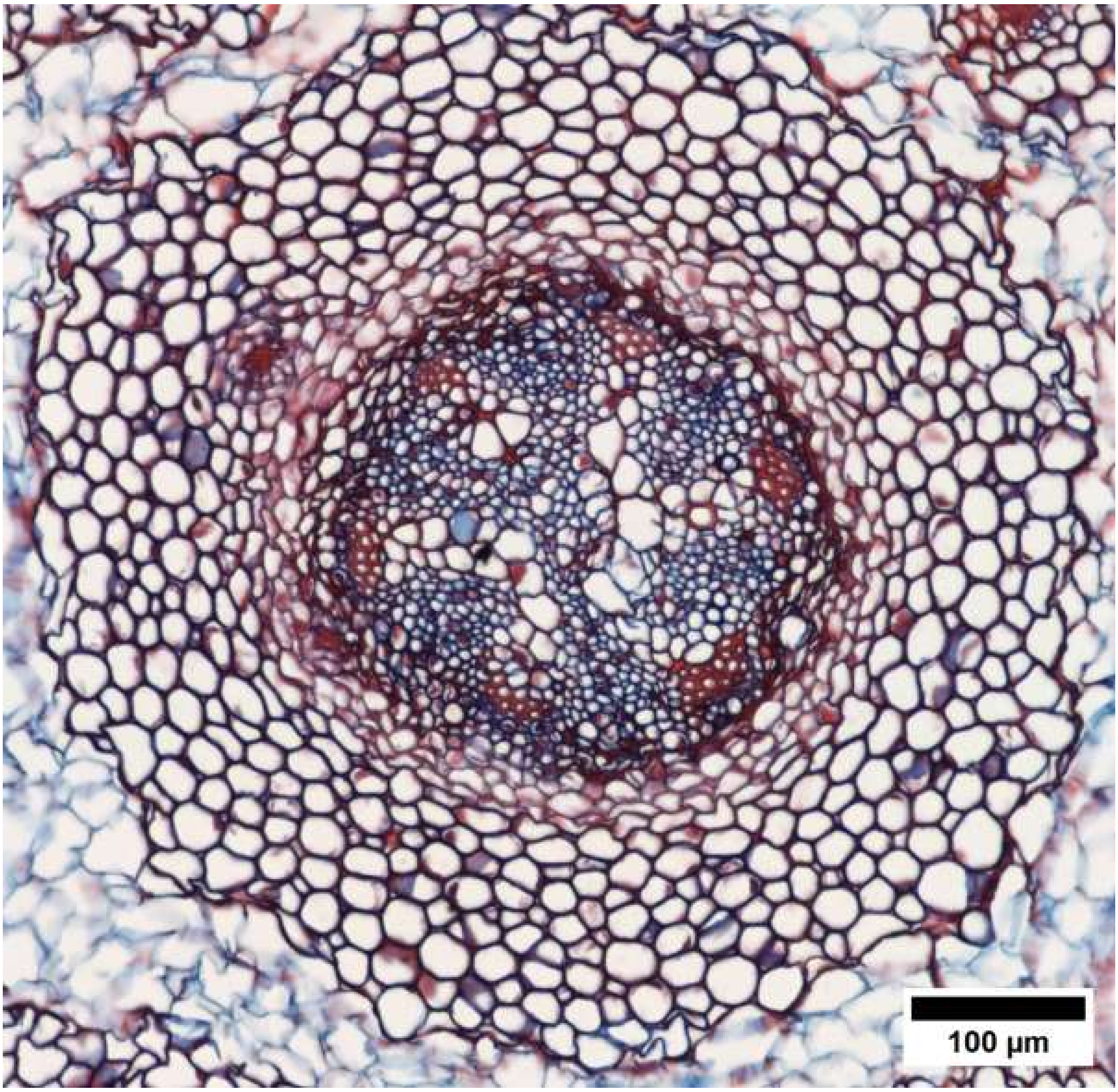


*Lycopodium clavatum* stem: L_clav_stem_2

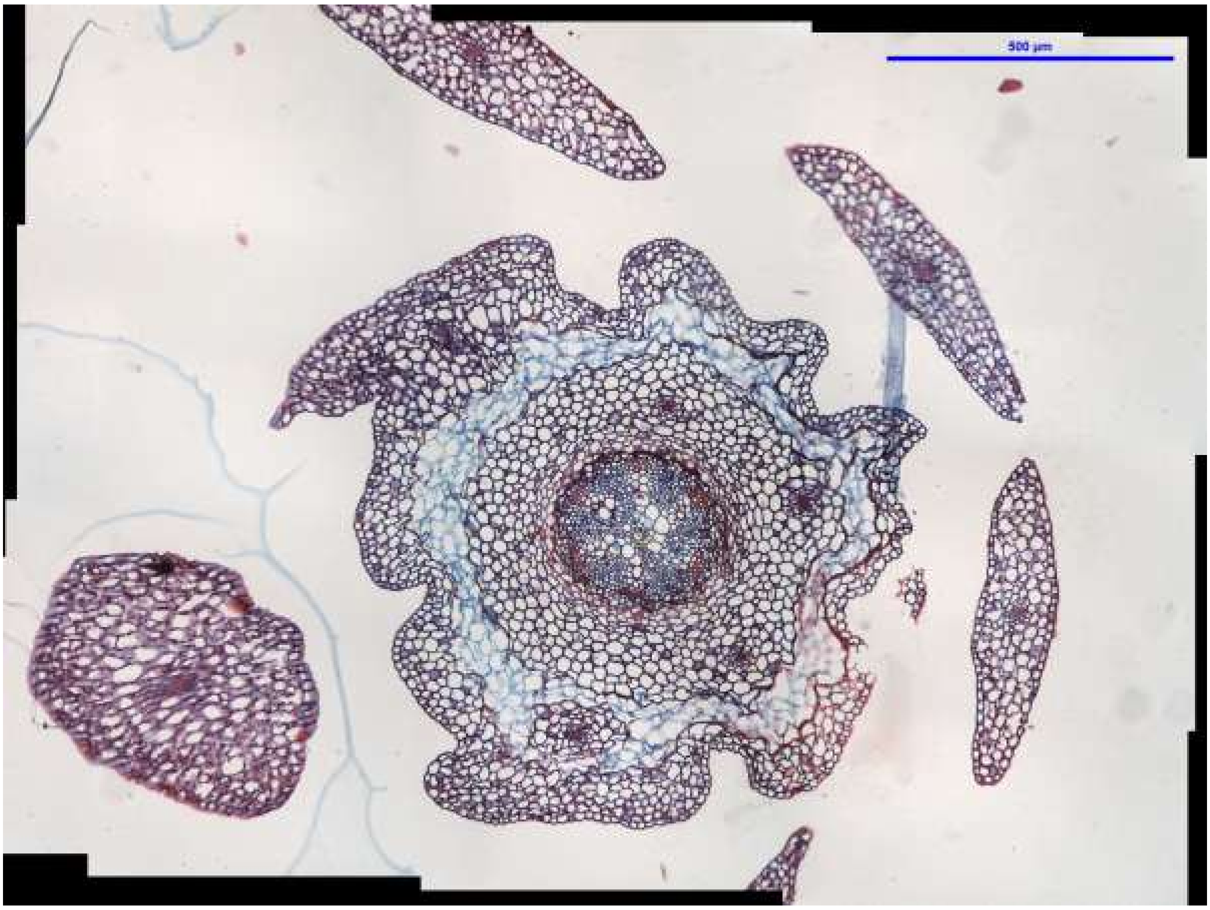
500 µm

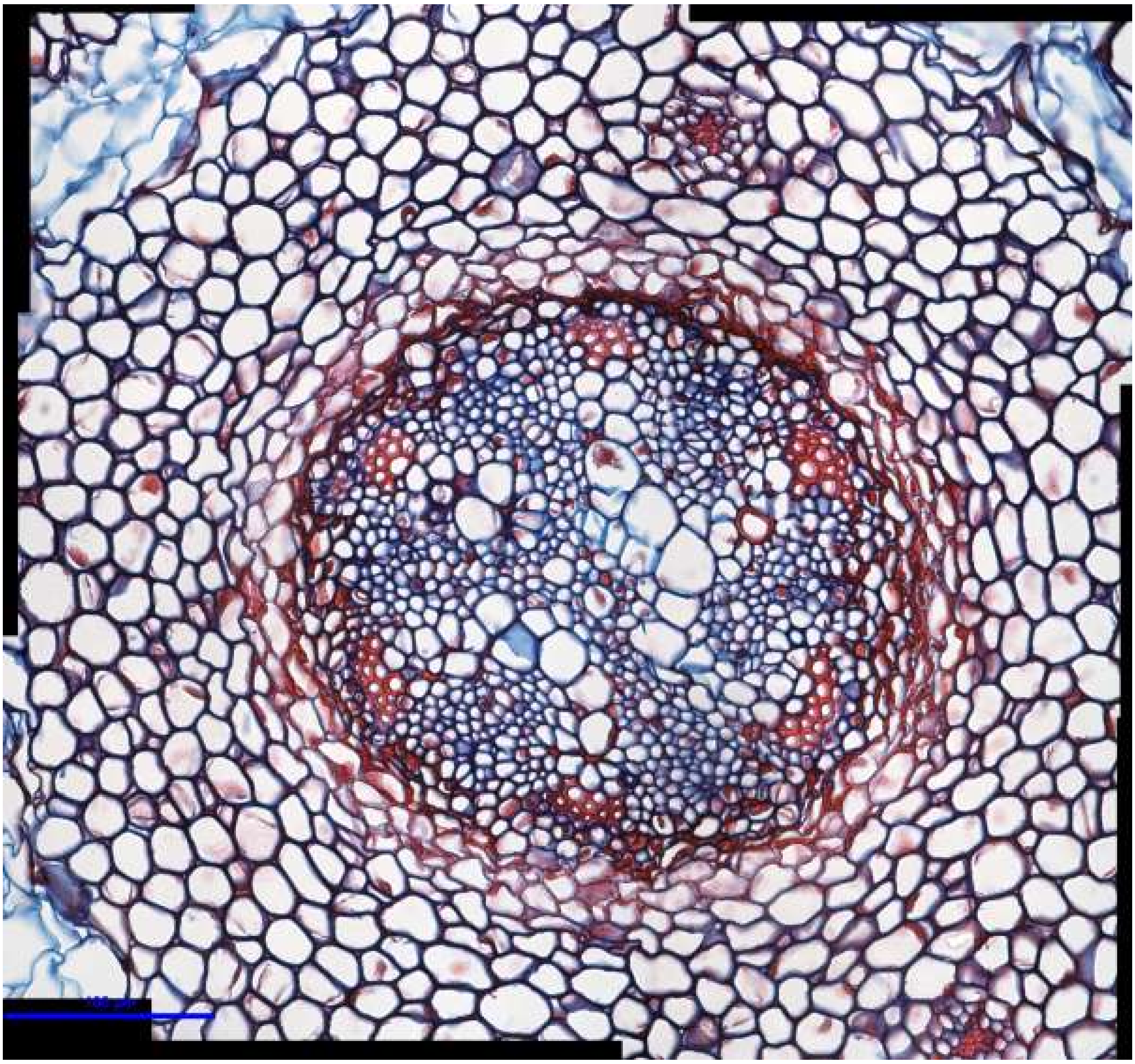

*Lycopodium clavatum* stem: L_clav_stem_3

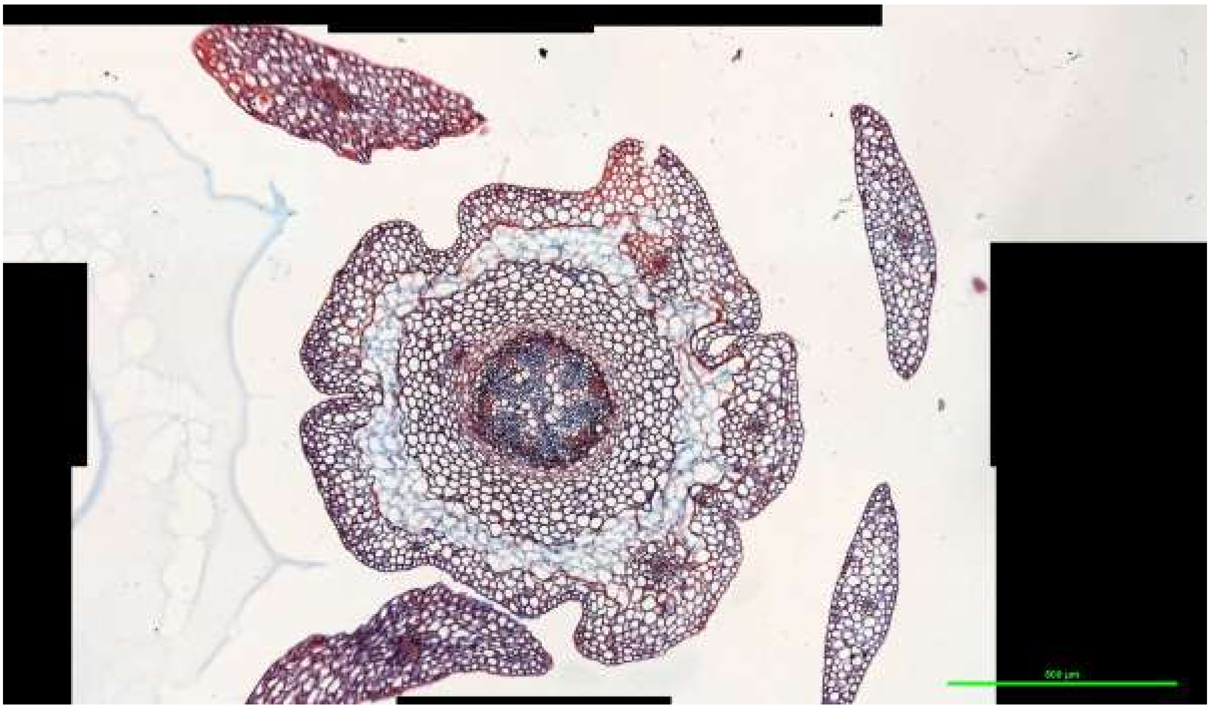

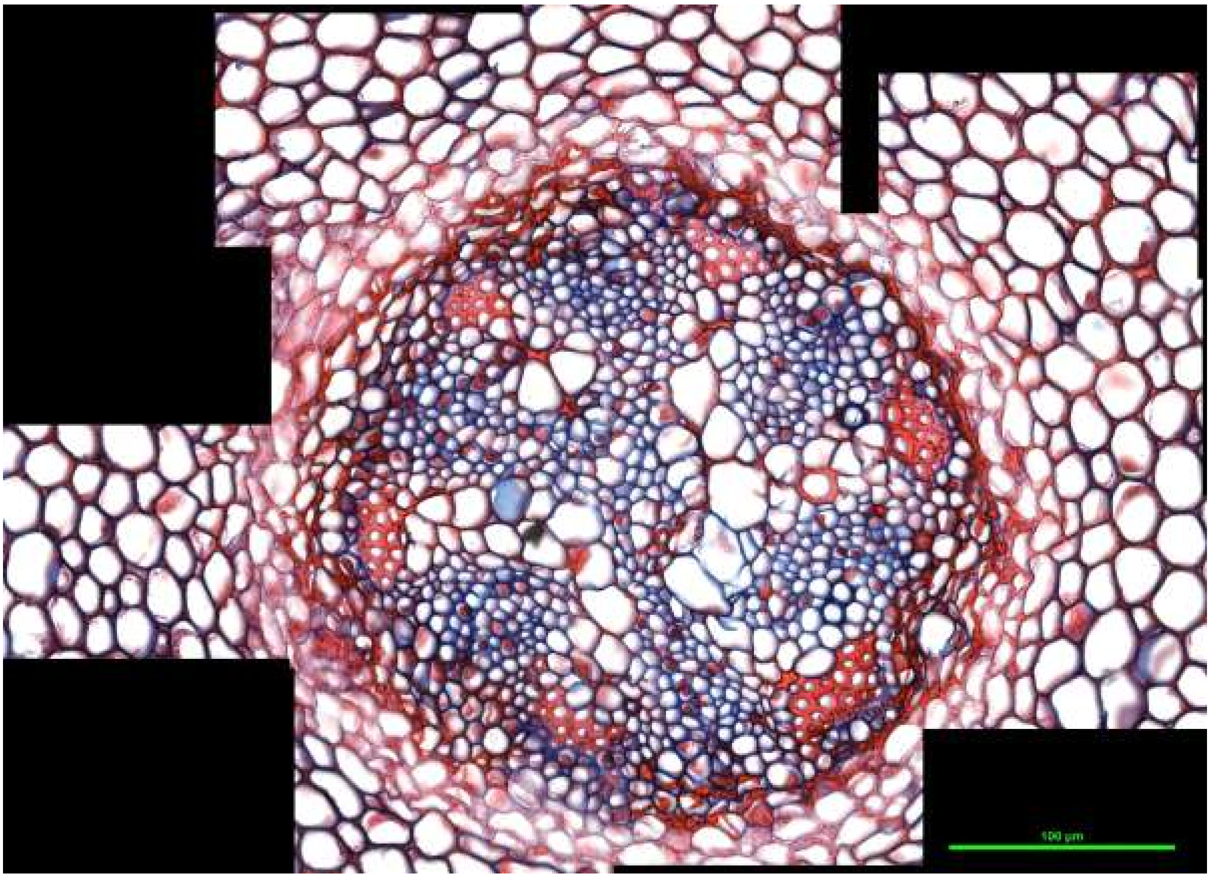

*Lycopodium clavatum* stem: L_clav_stem_4

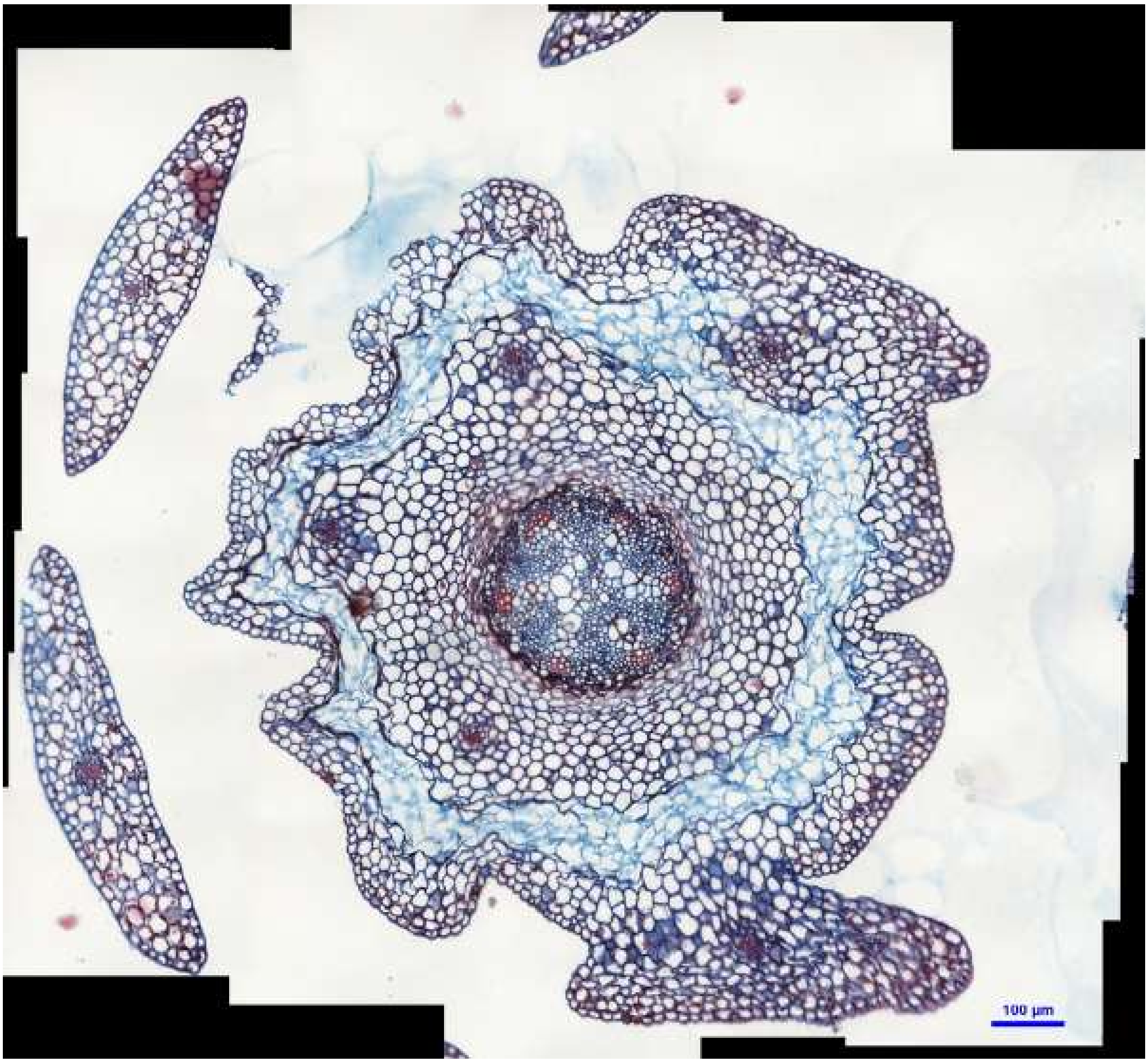
100 µm

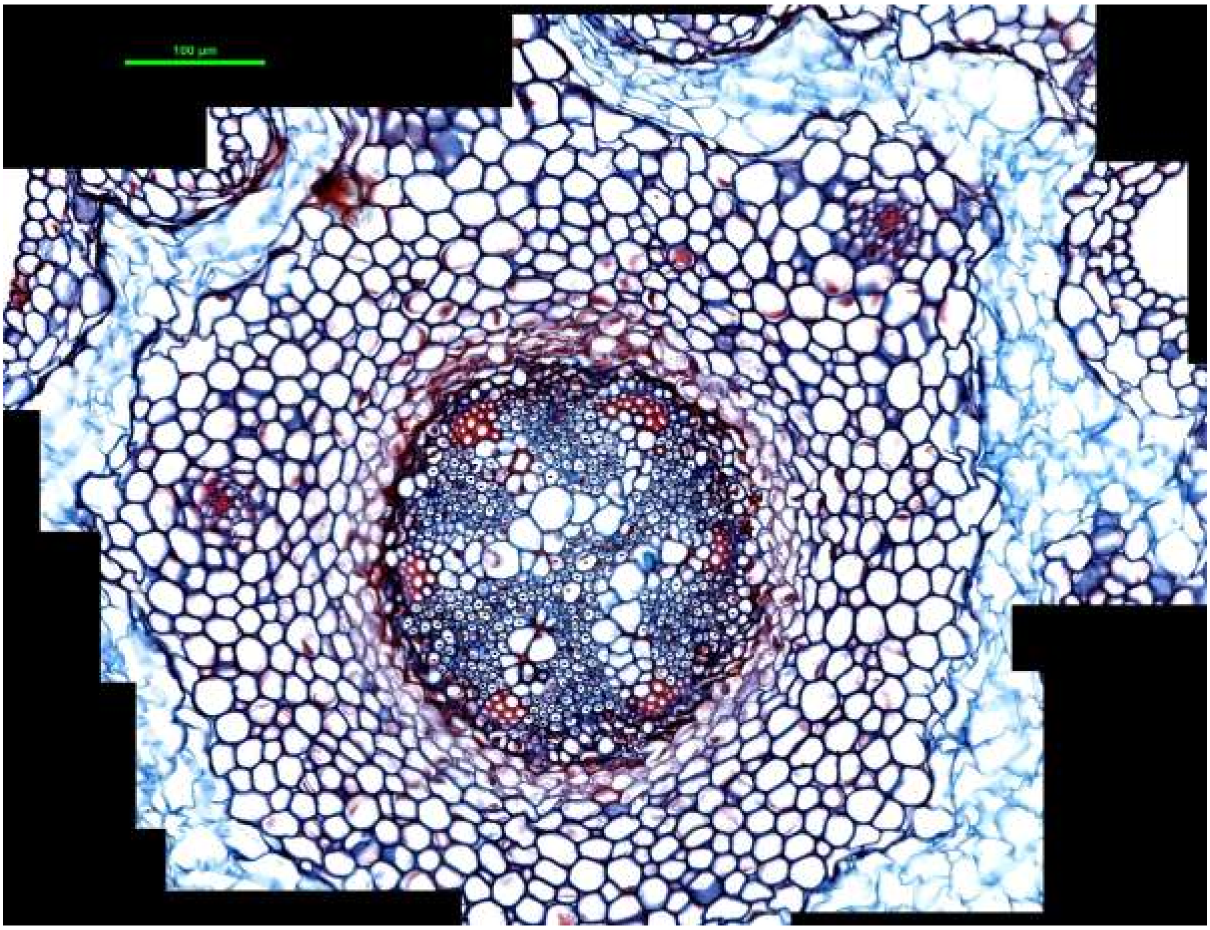


*Lycopodium clavatum* stem: L_clav_stem_5

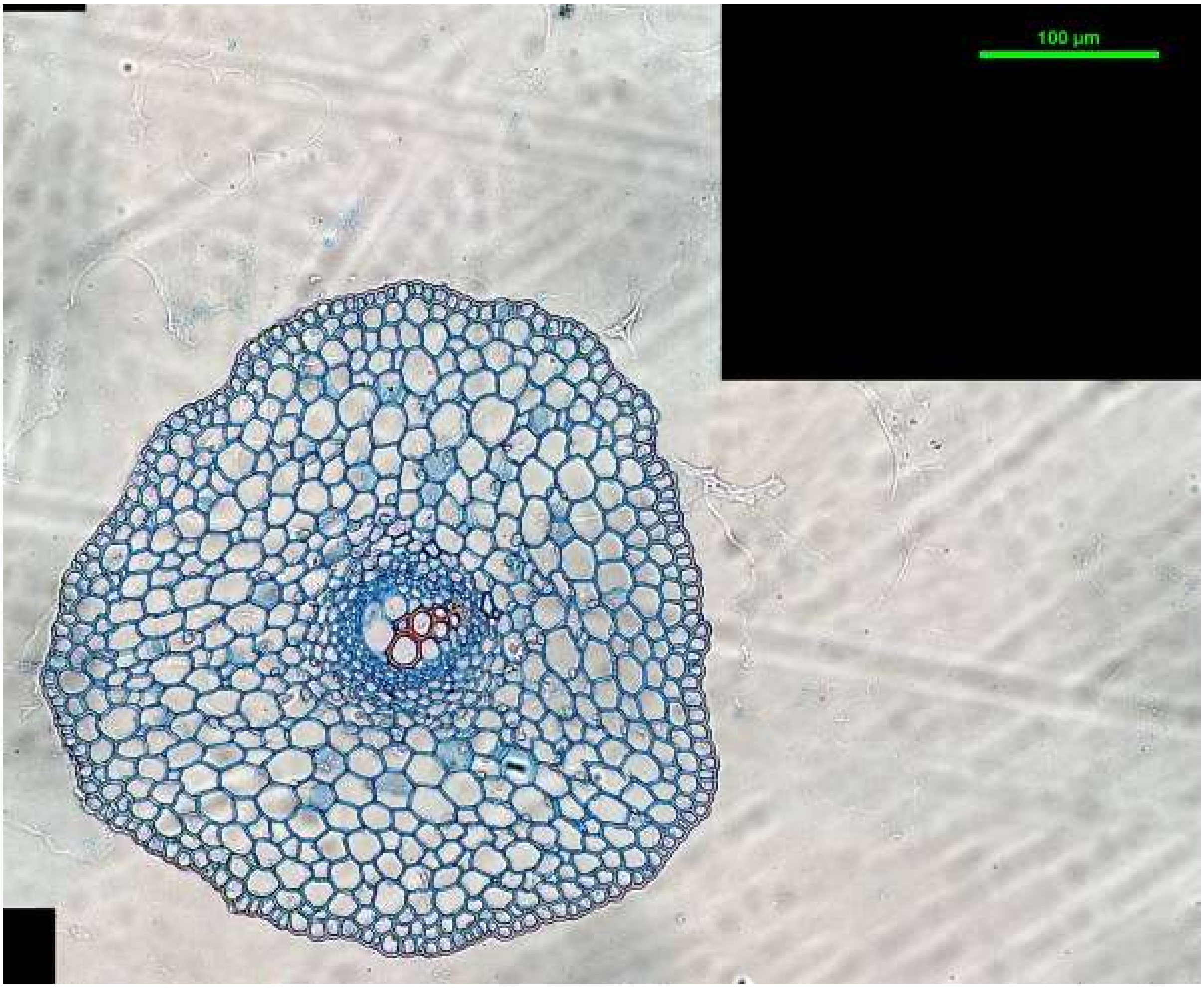
100 µm

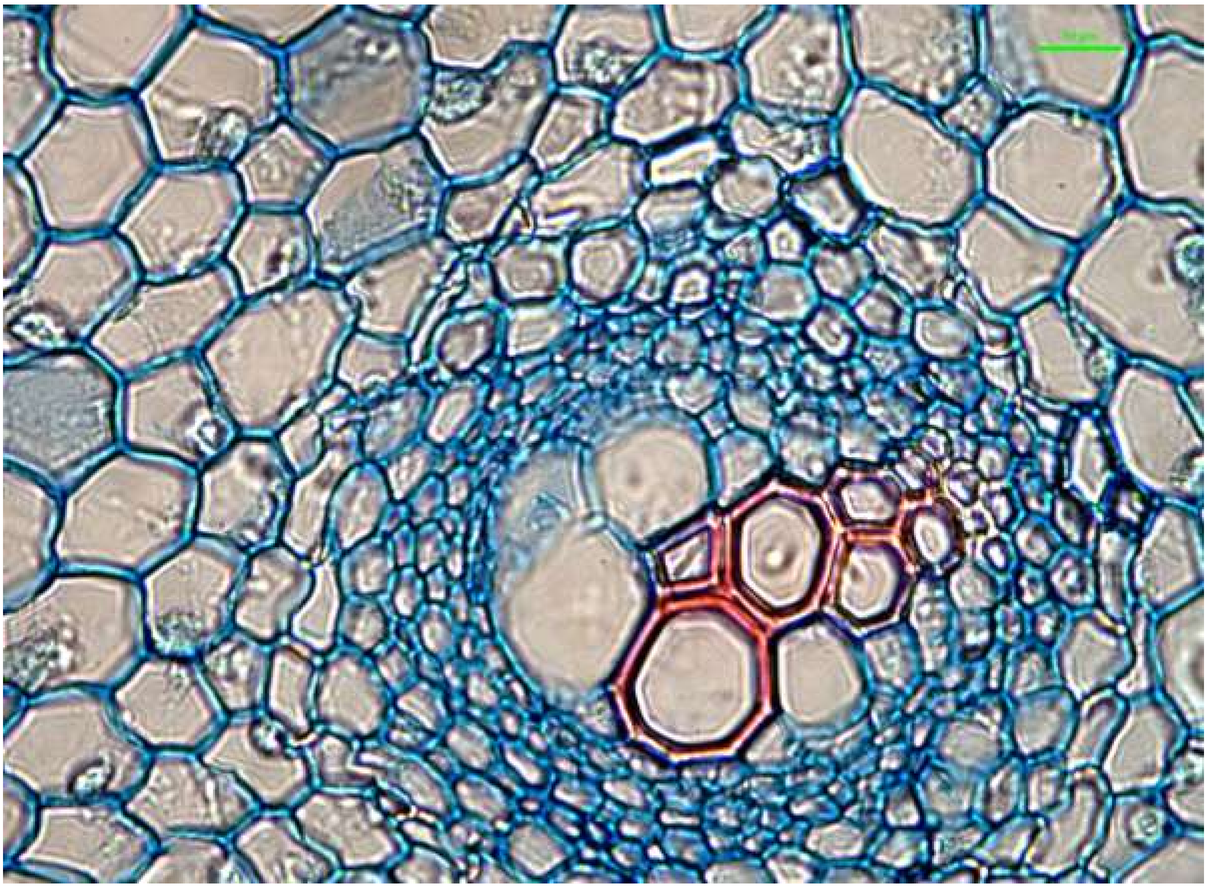

*Selaginella uncinata* root: S_unc_root_1

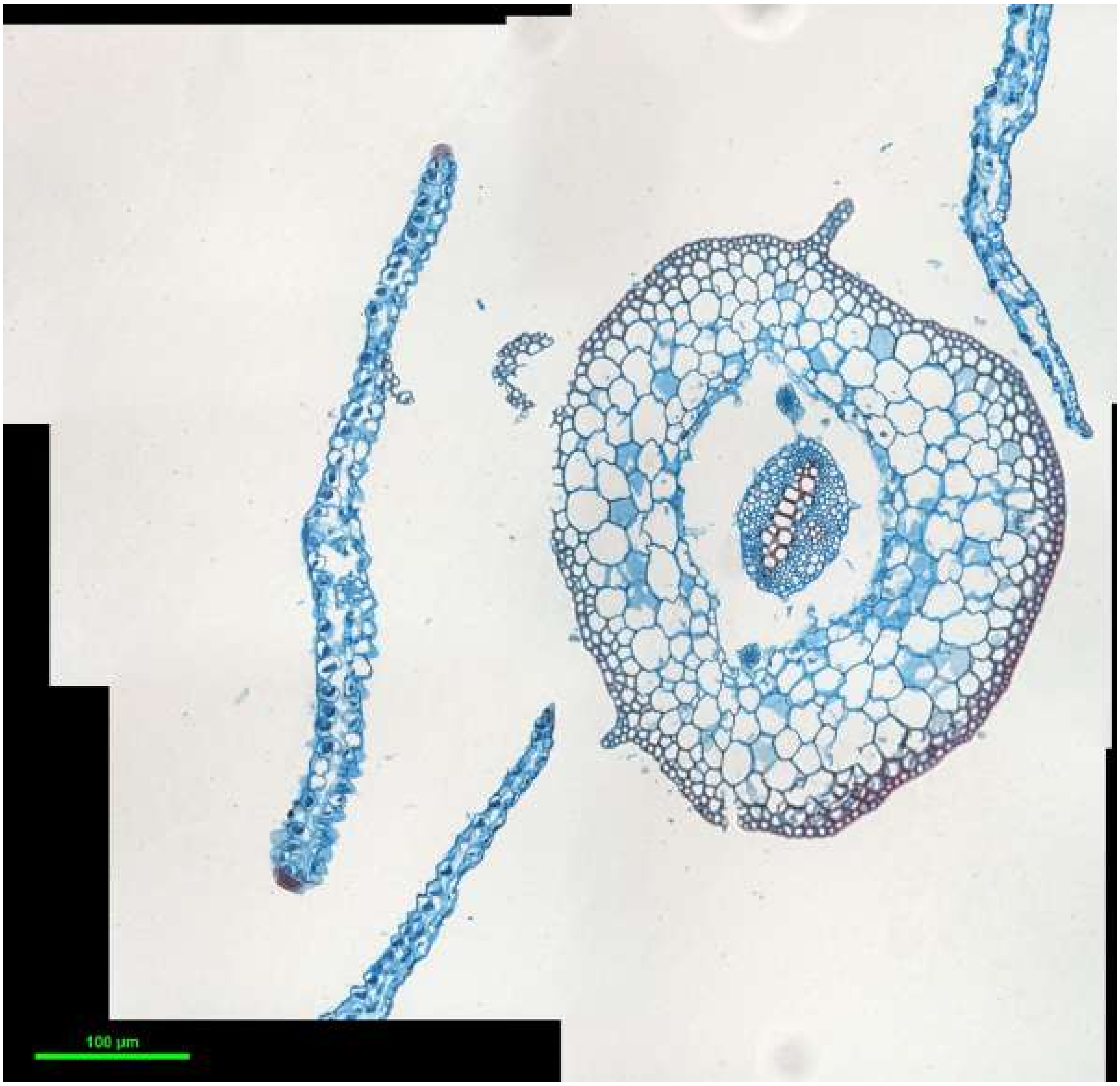
100 µm

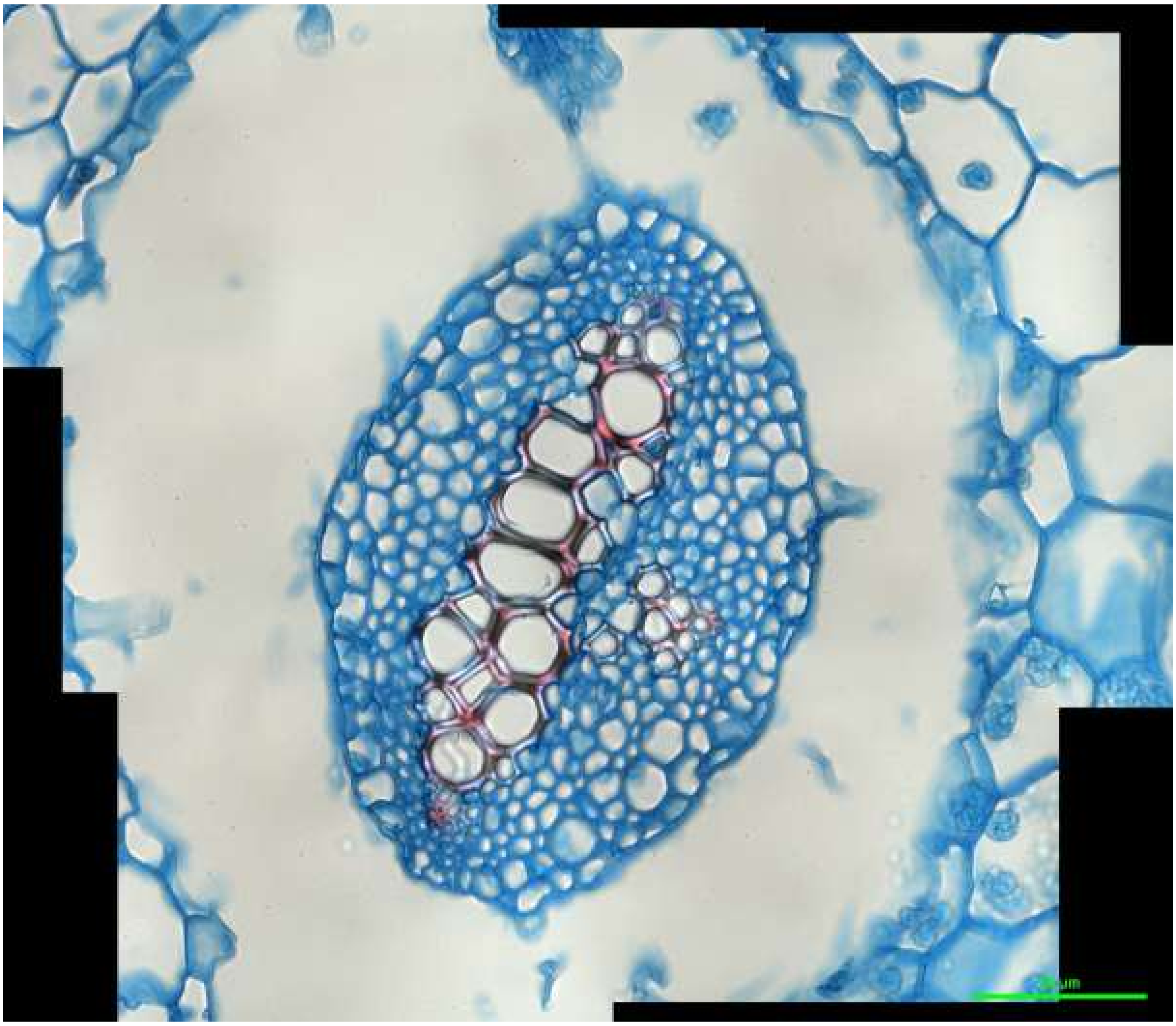

*Selaginella uncinata* stem: S_unc_stem_1

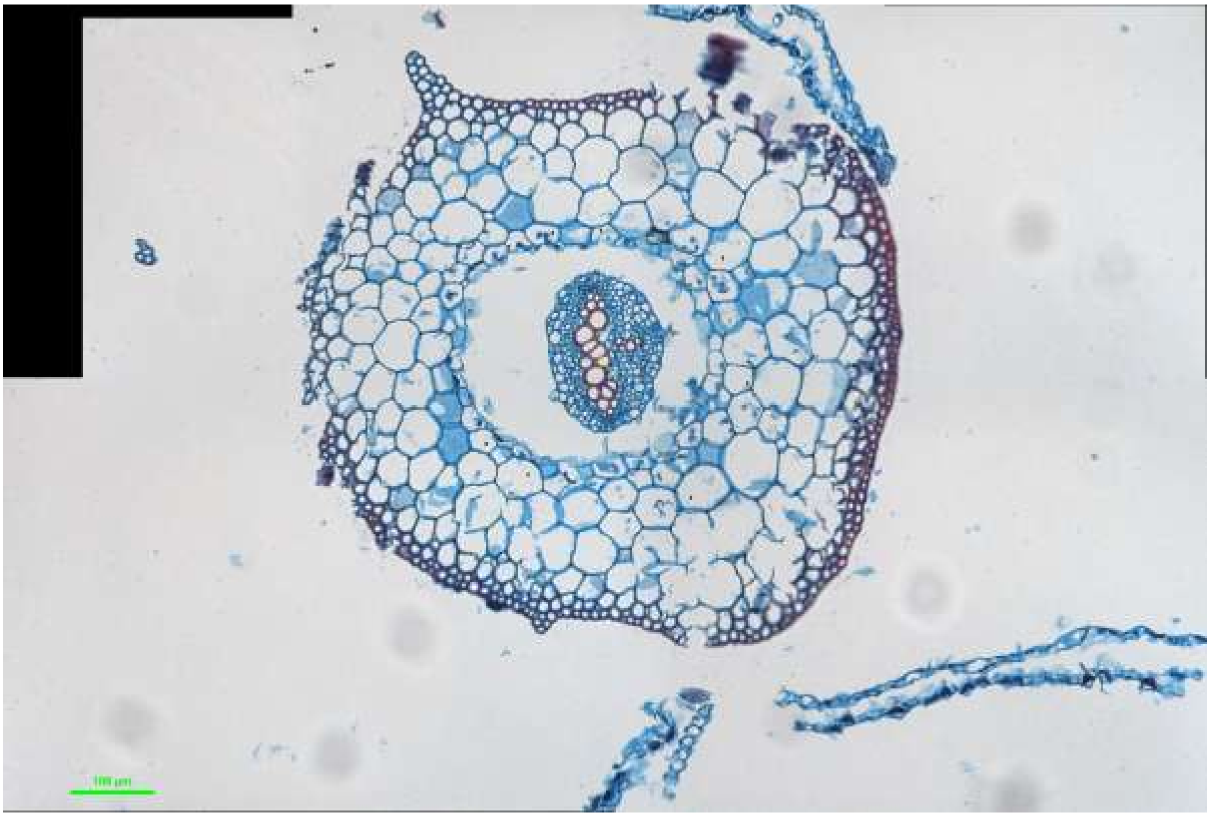

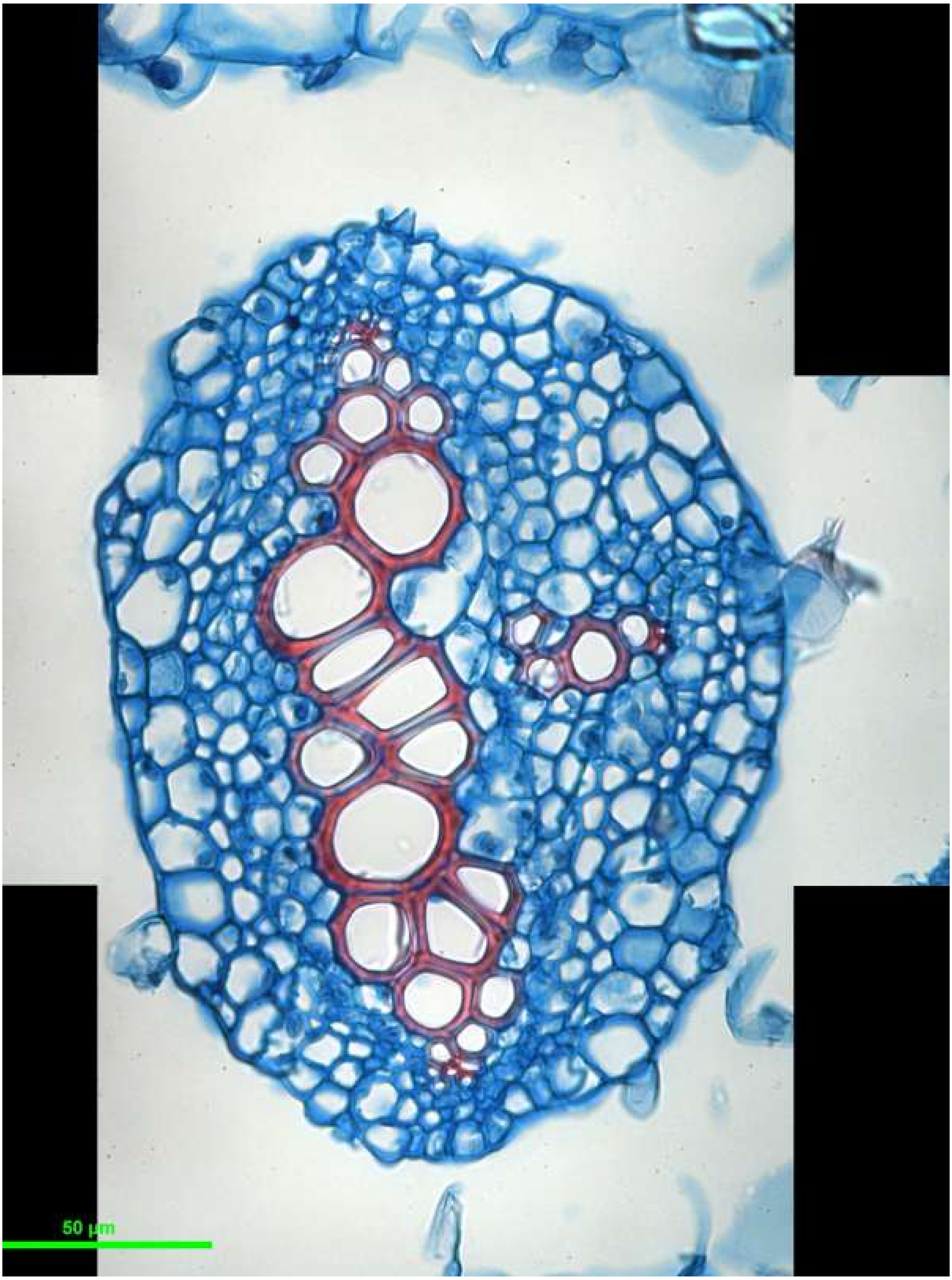


*Selaginella uncinata* stem: S_unc_stem_2

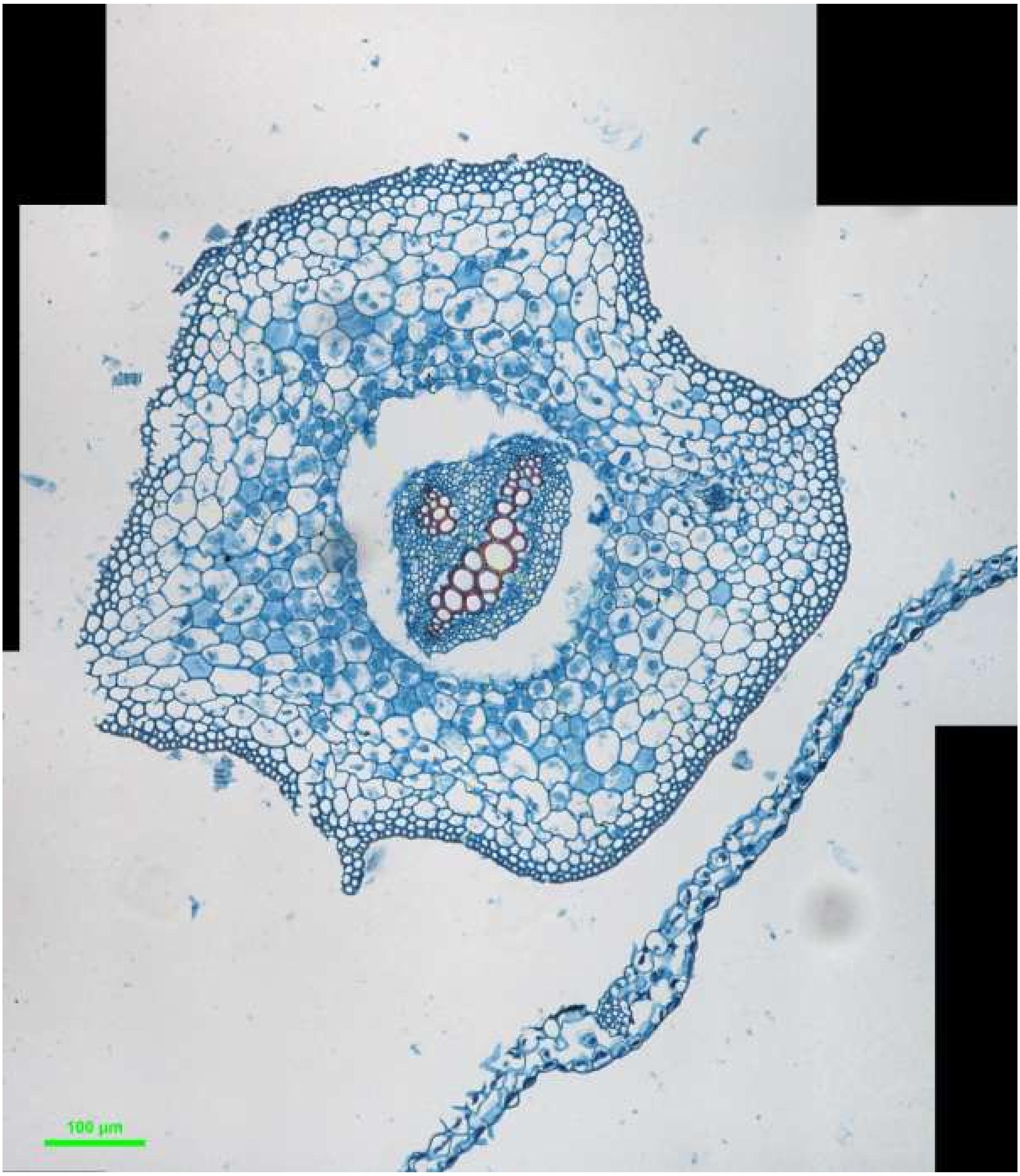
100 µm

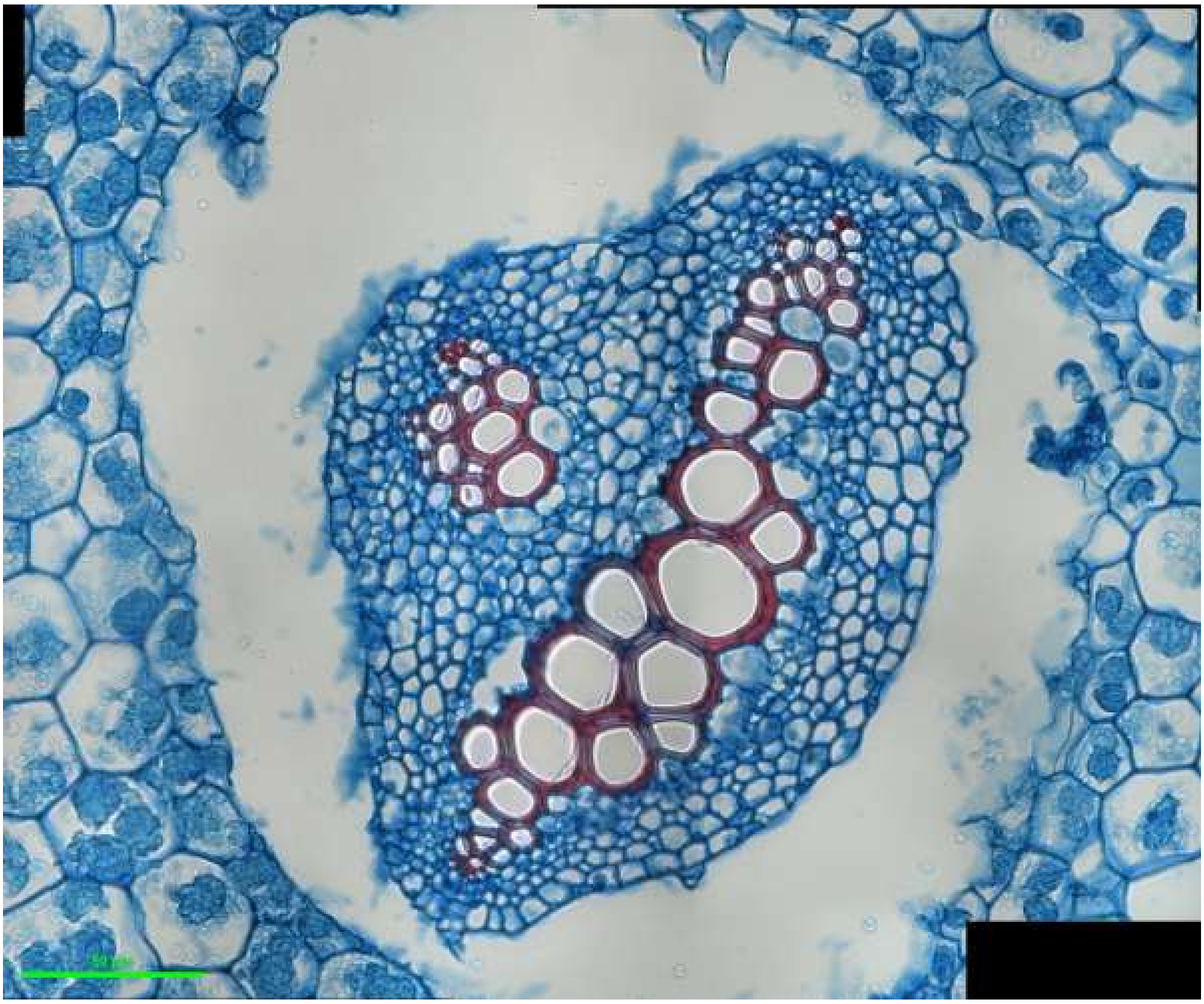

*Selaginella uncinata* stem: S_unc_stem_3

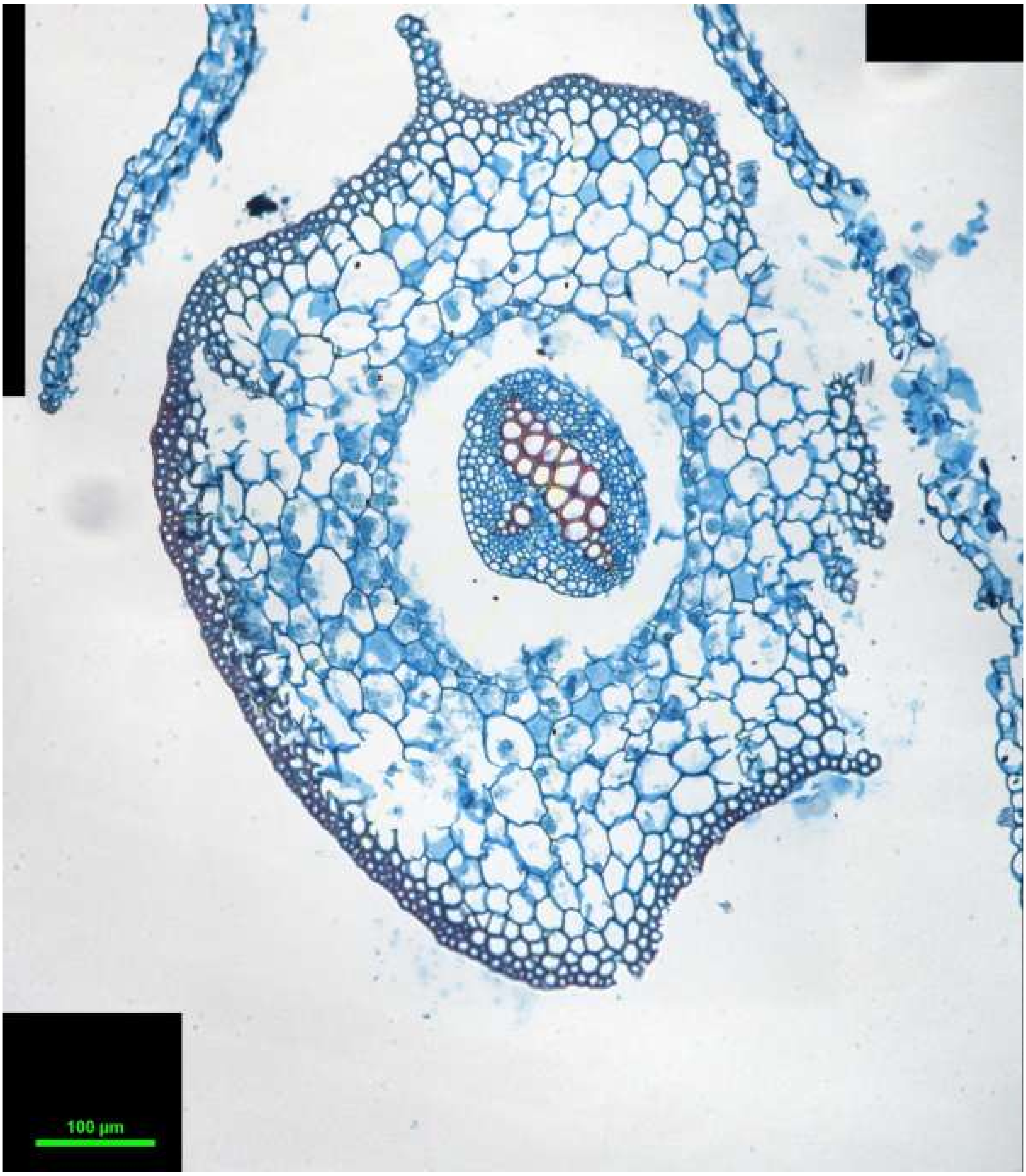
100 µm

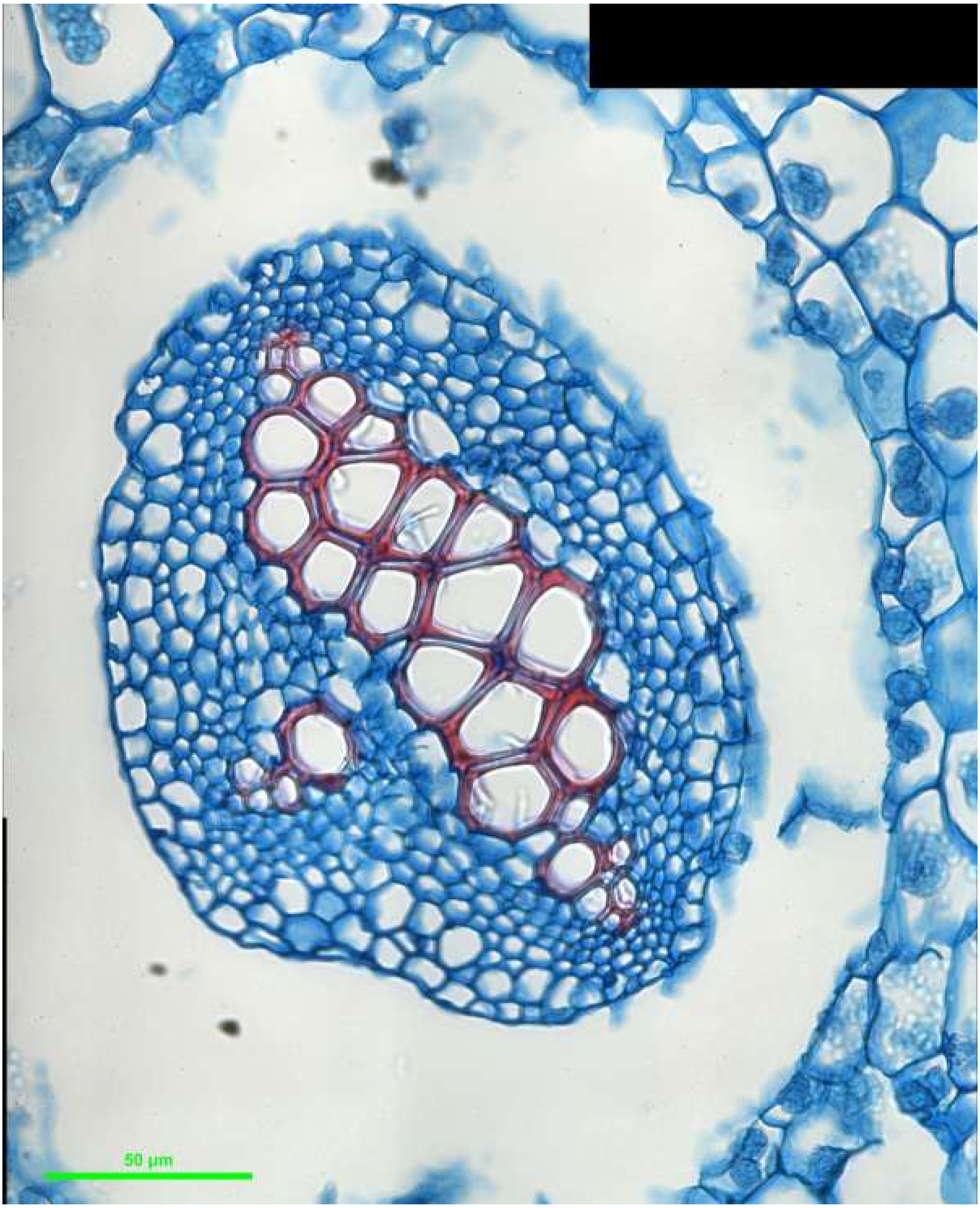


*Selaginella uncinata* stem: S_unc_stem_4

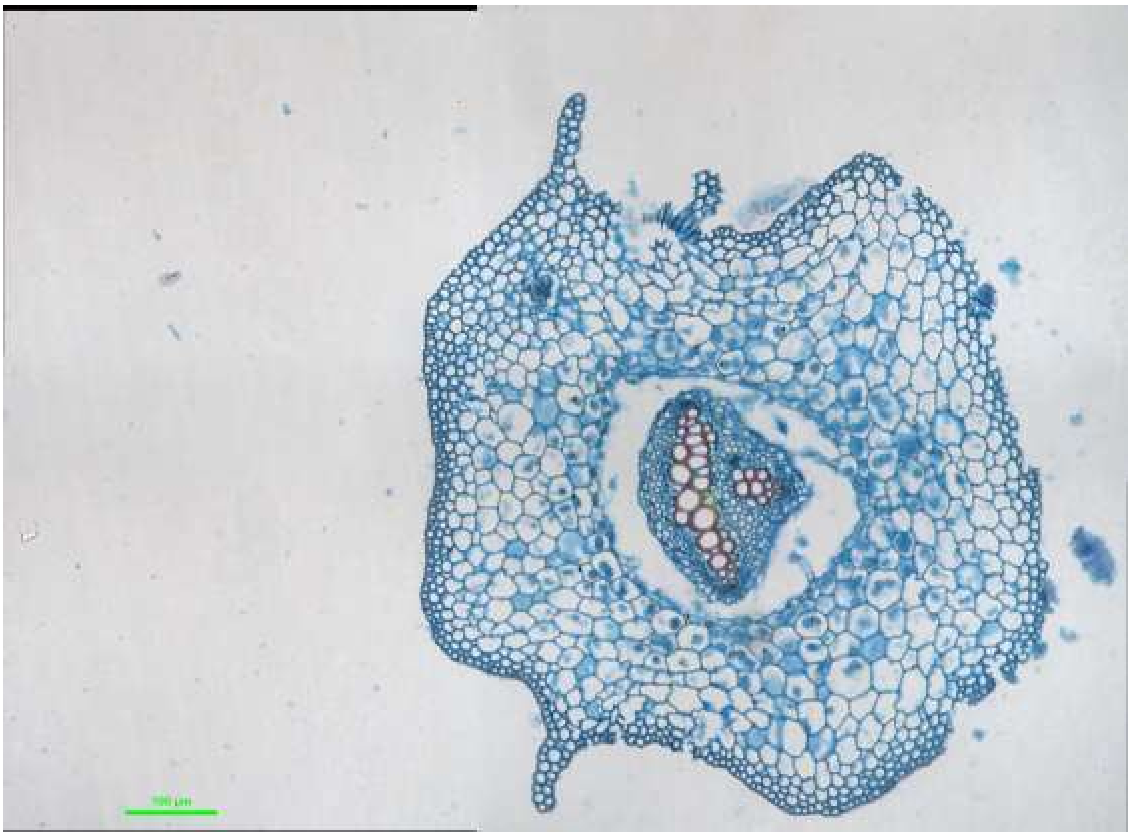

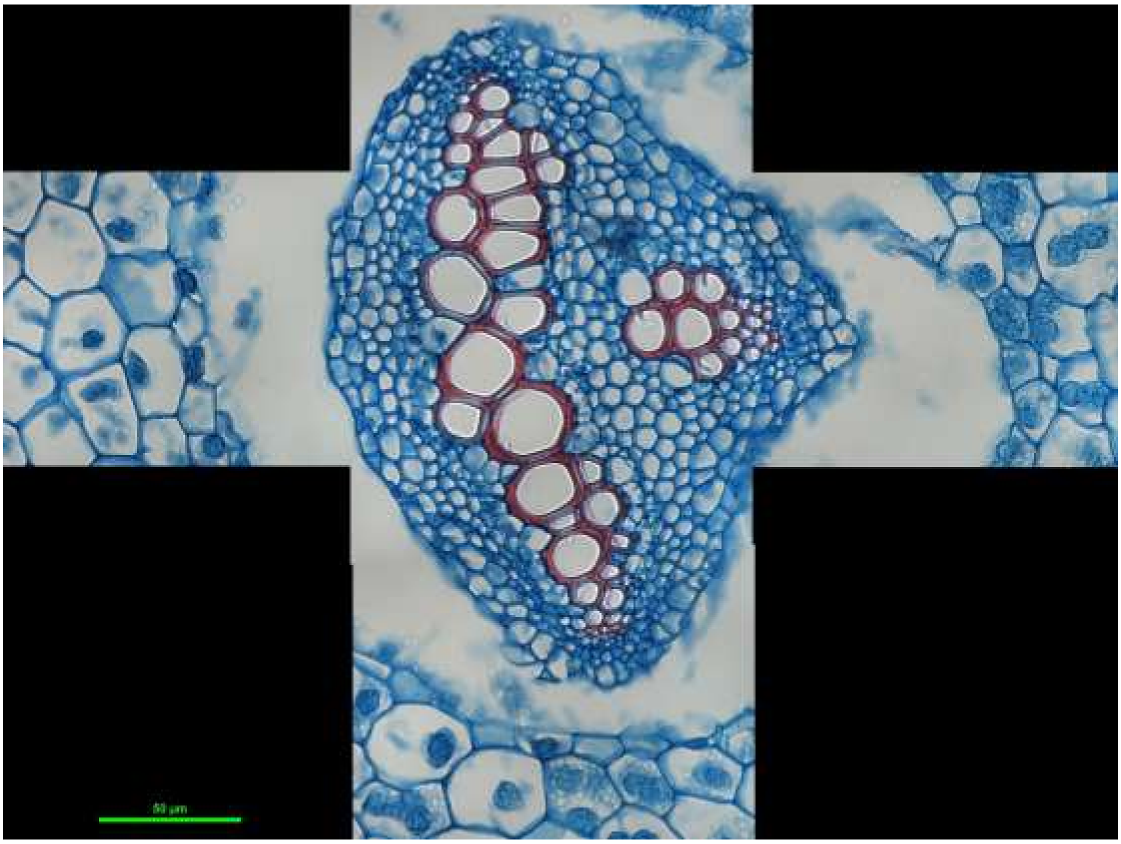

*Selaginella uncinata* stem: S_unc_stem_5

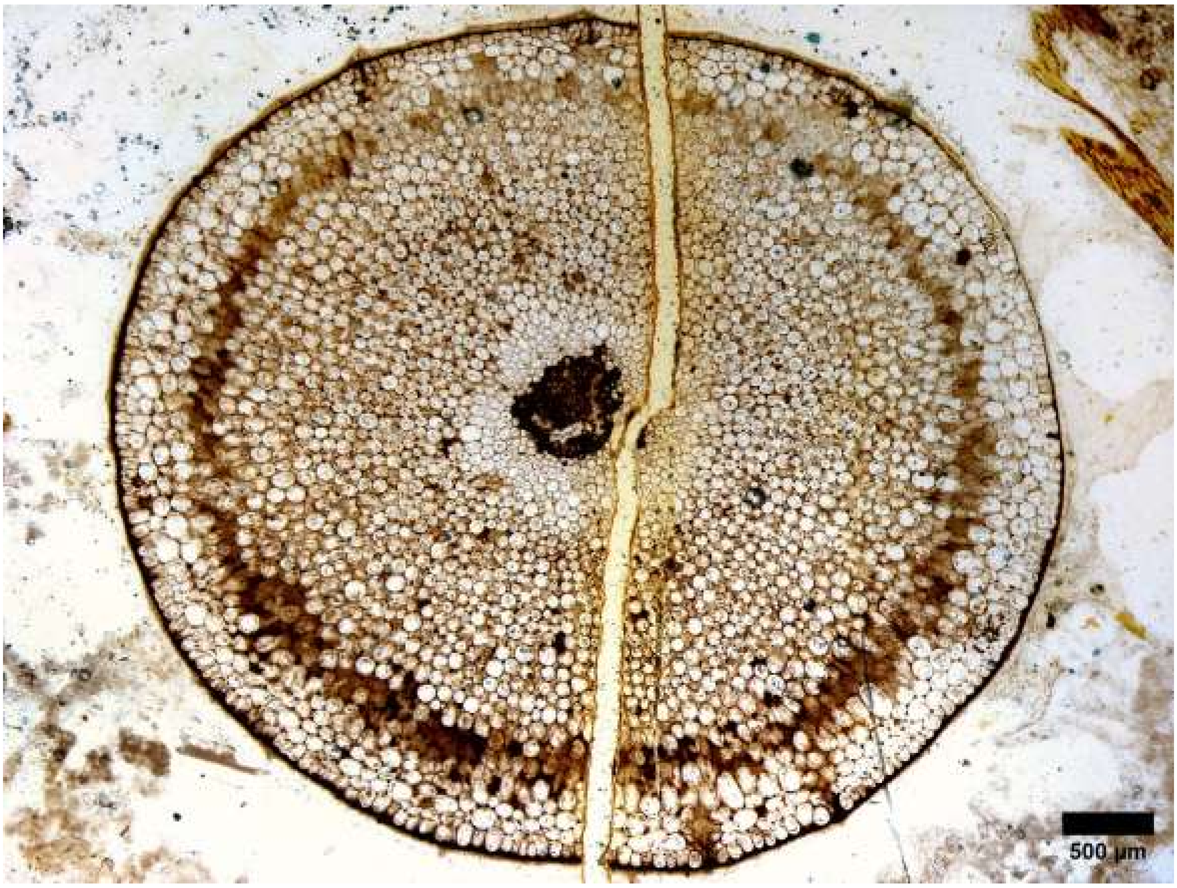


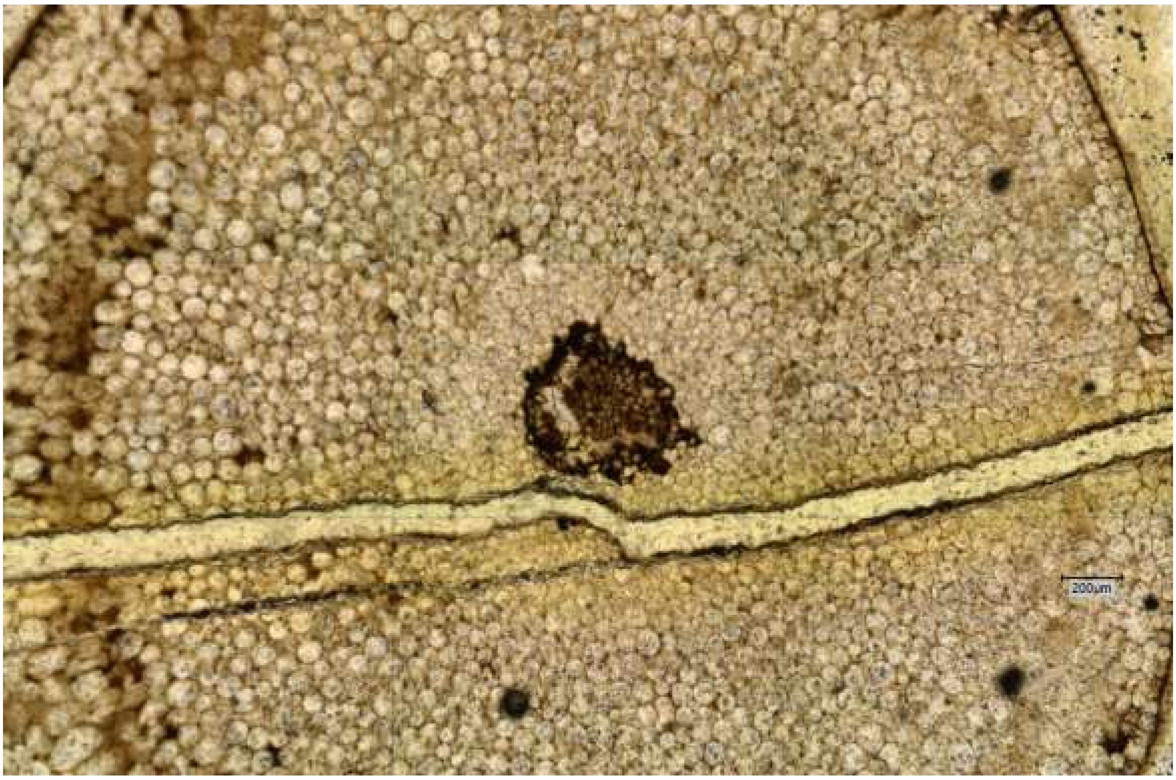


*Aglaophyton majus* (Pb1870): Aglao_1

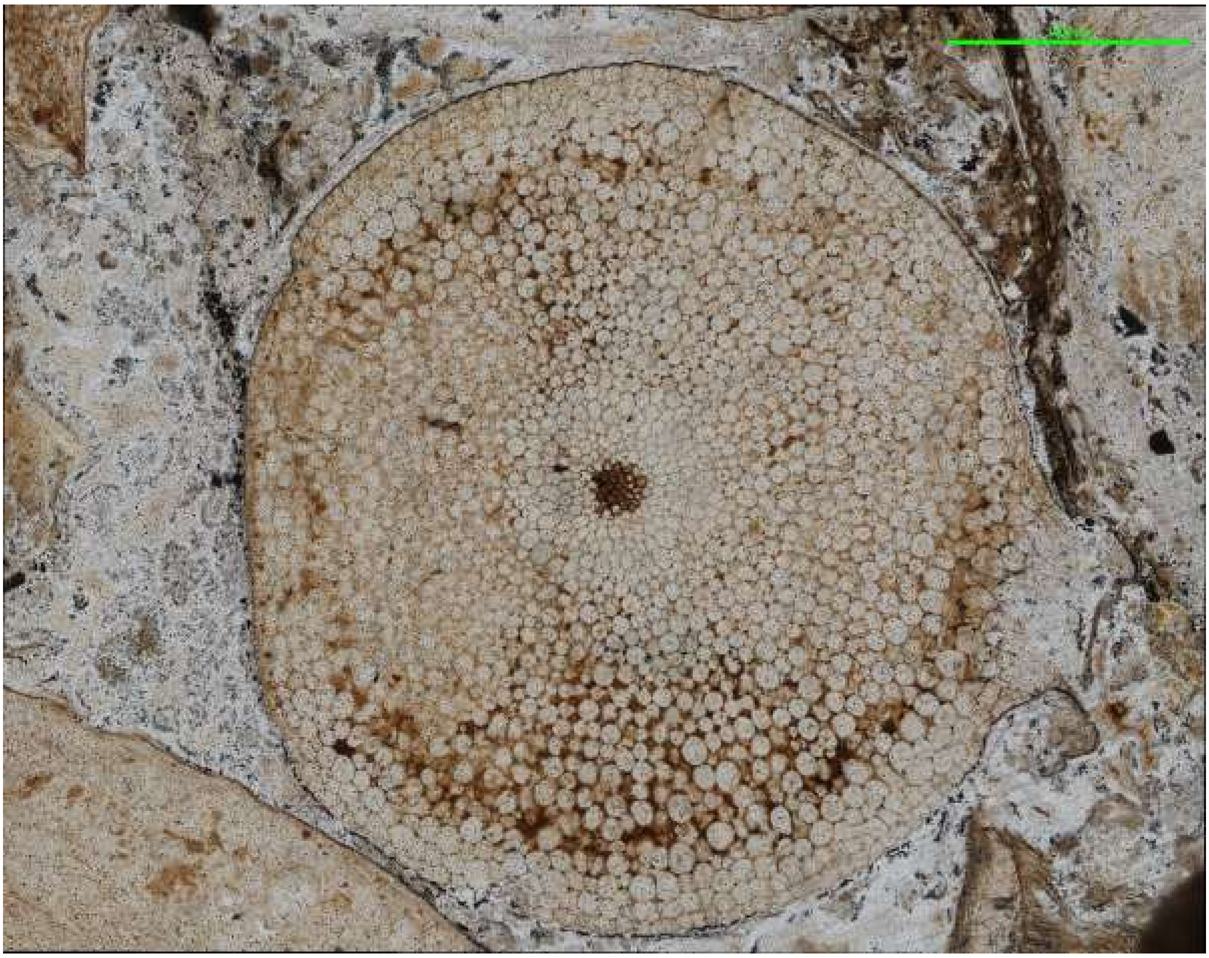

*Aglaophyton majus* (SCOTT RC59): Aglao_2

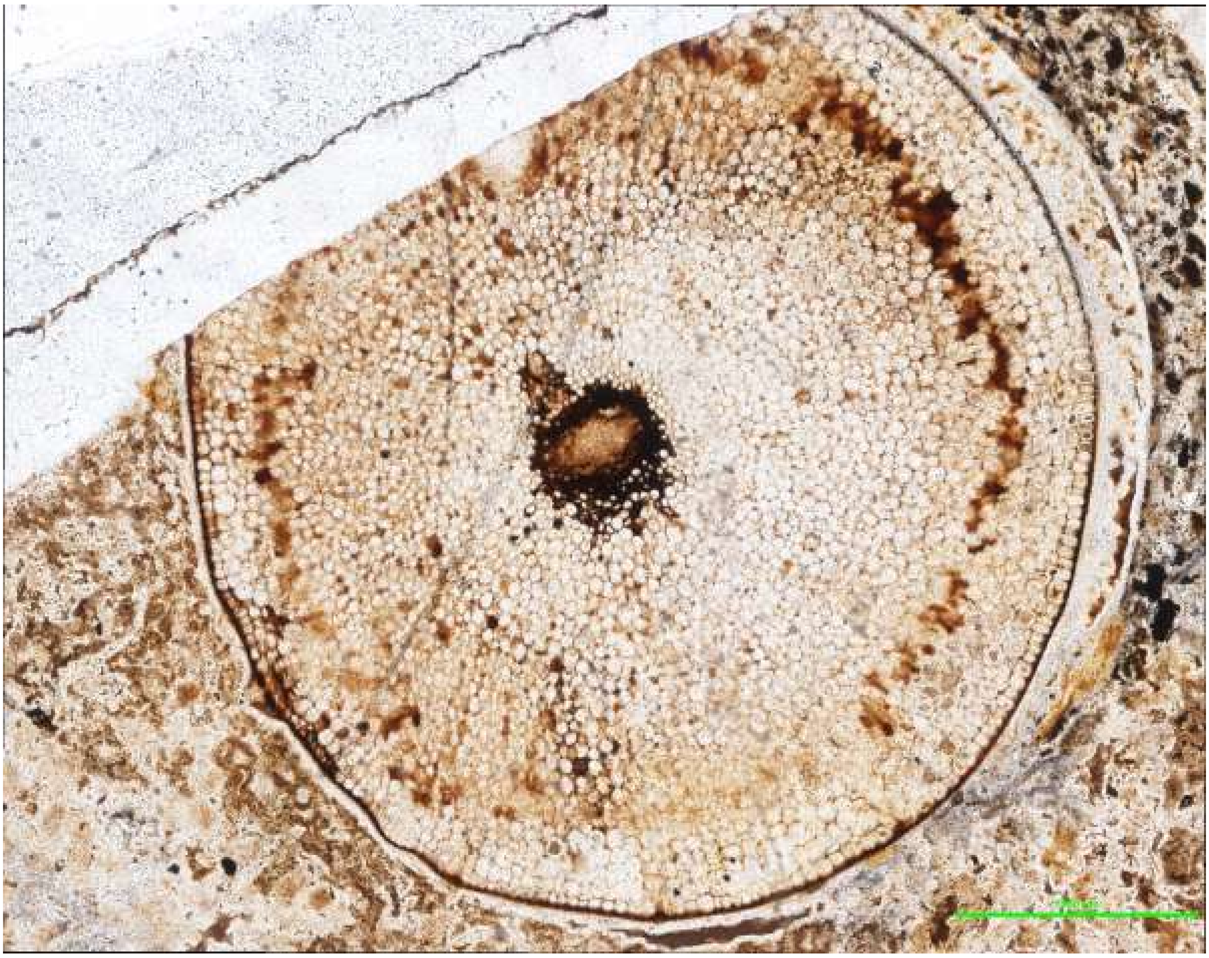

*Aglaophyton majus* (SCOTT RY36): Aglao_3

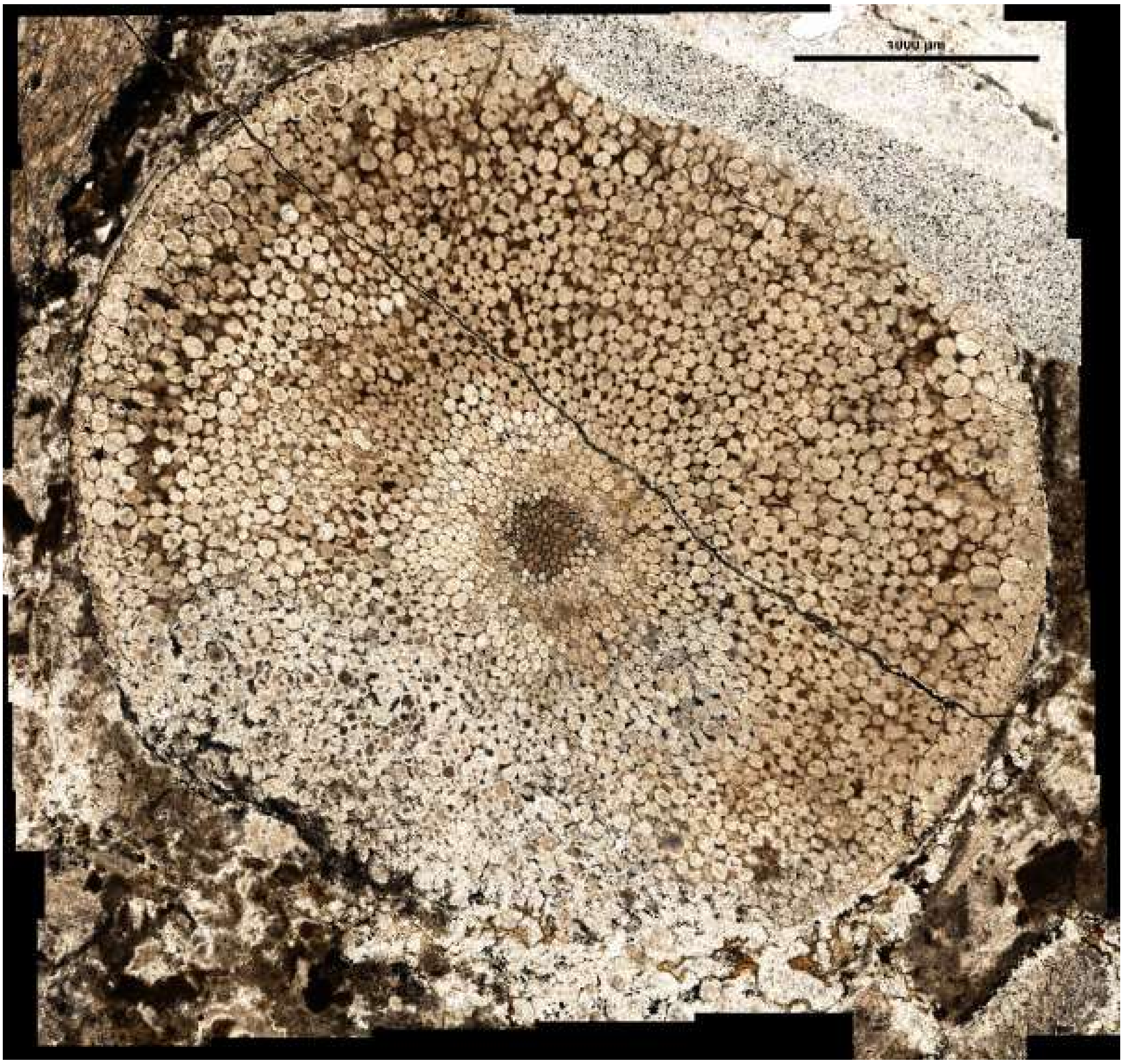

*Aglaophyton majus* (MPEG007): Aglao_4

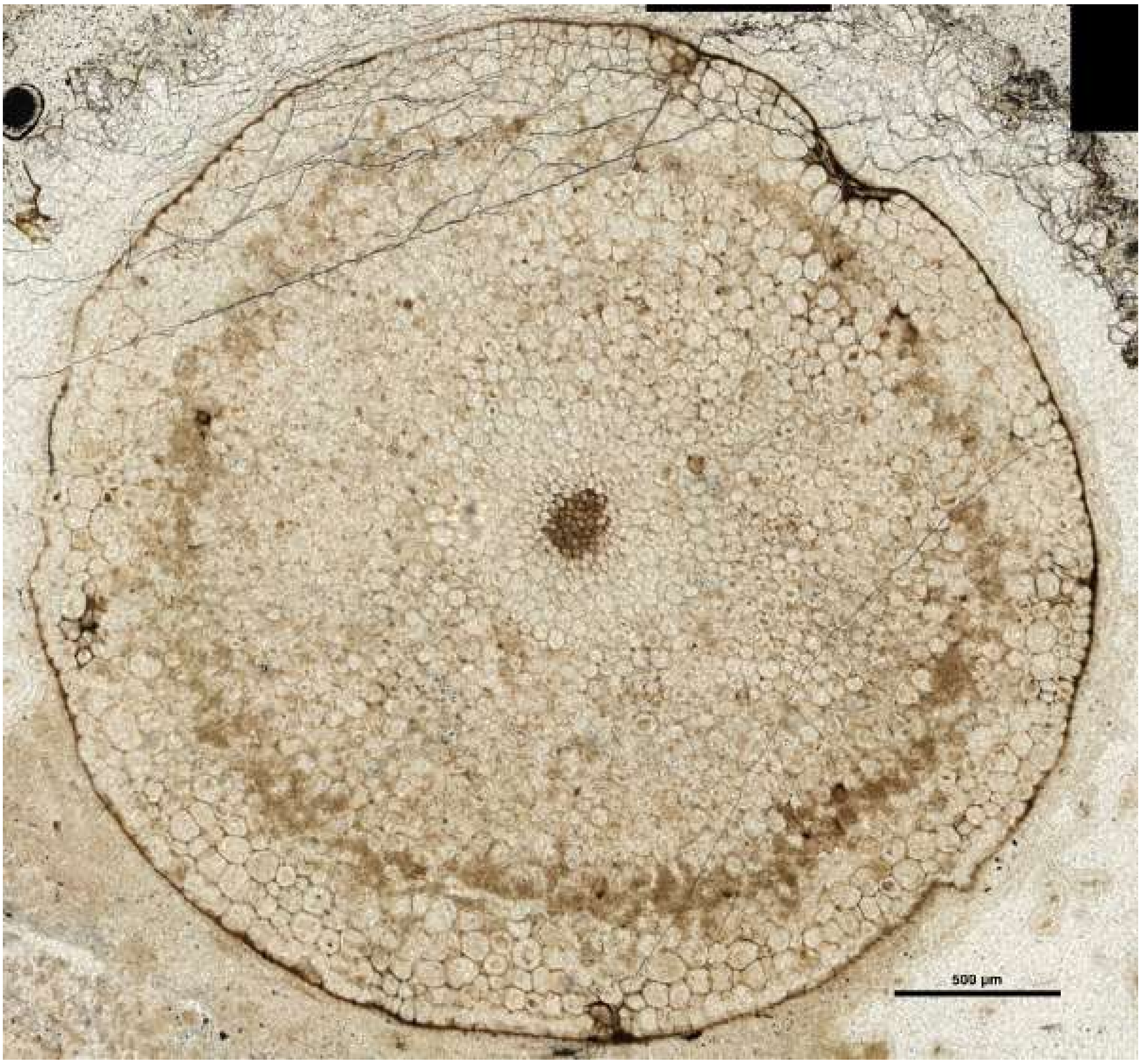
500 µm

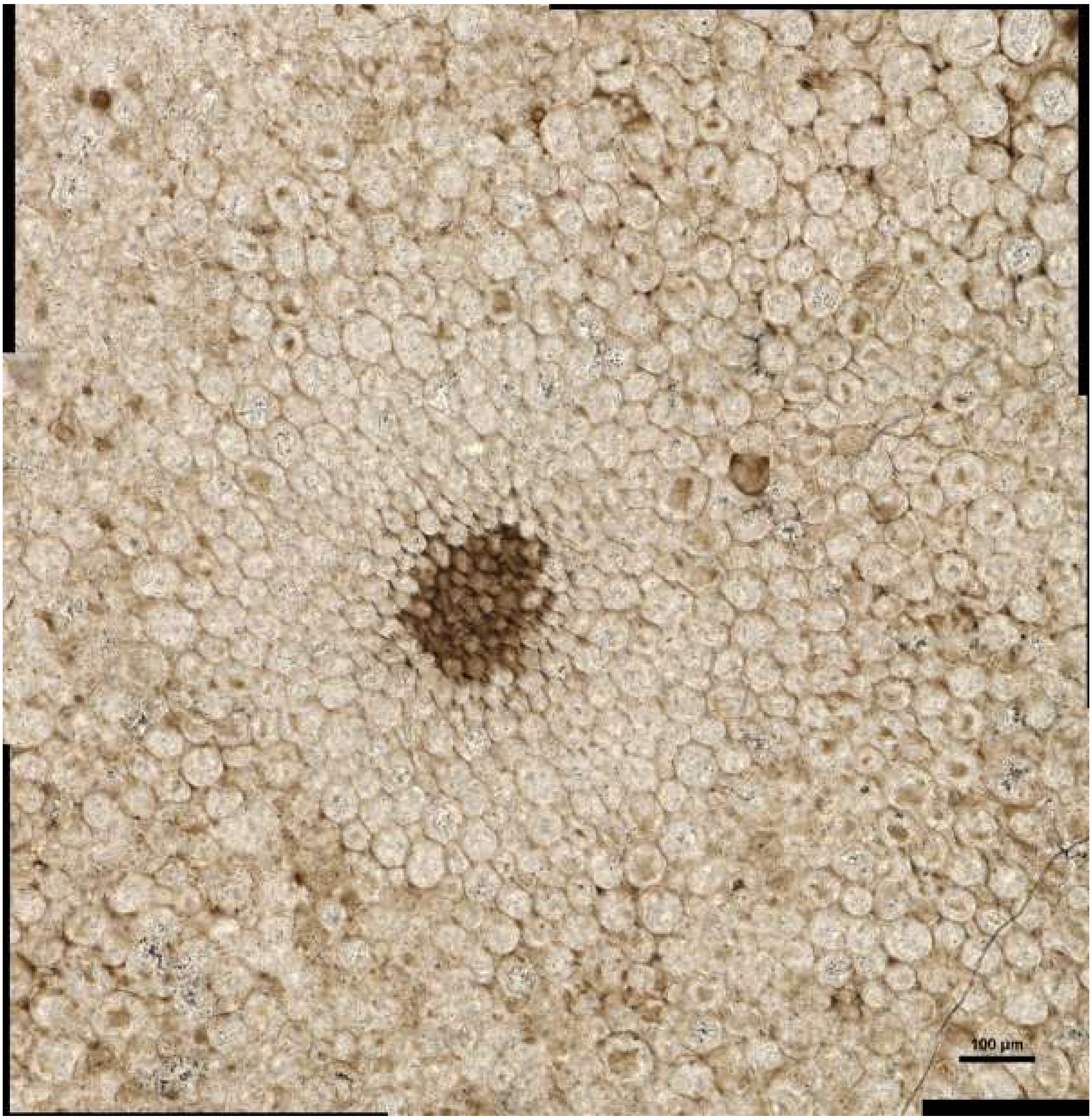


*Aglaophyton majus* (MPEG0015): Aglao_5

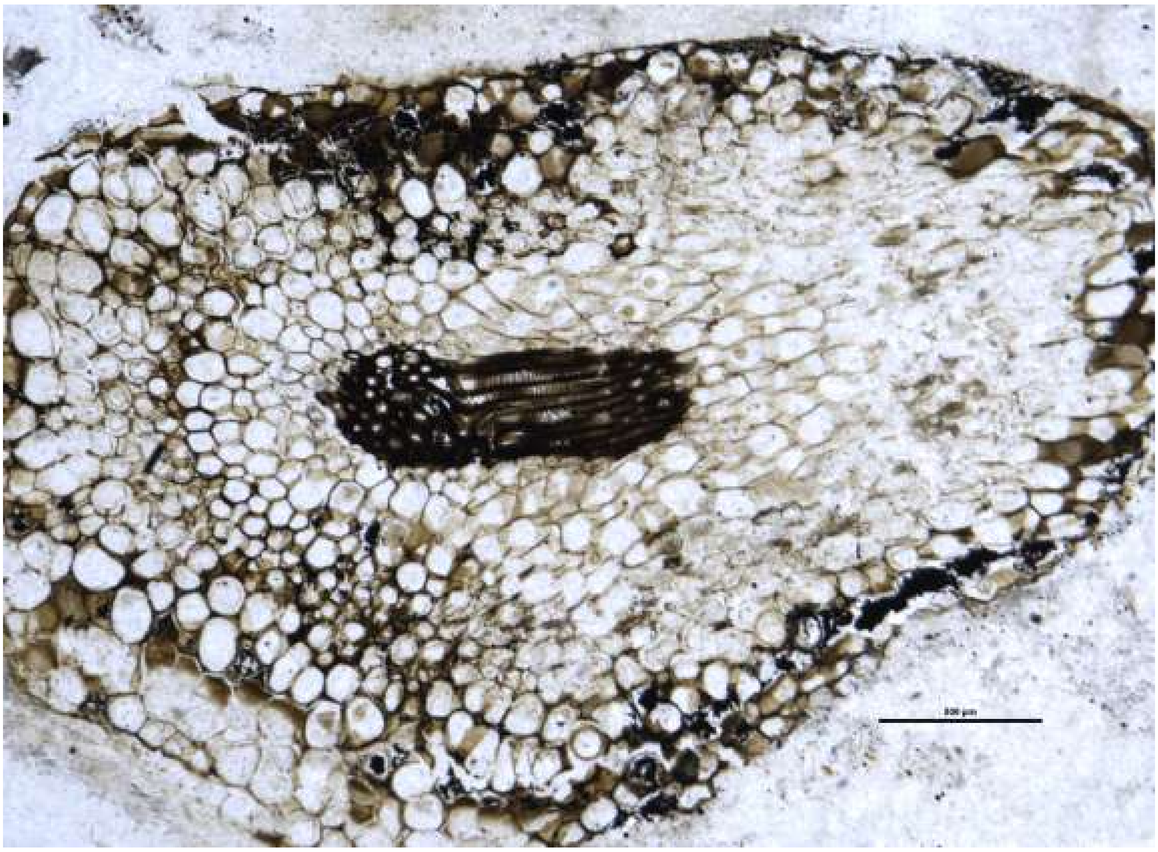

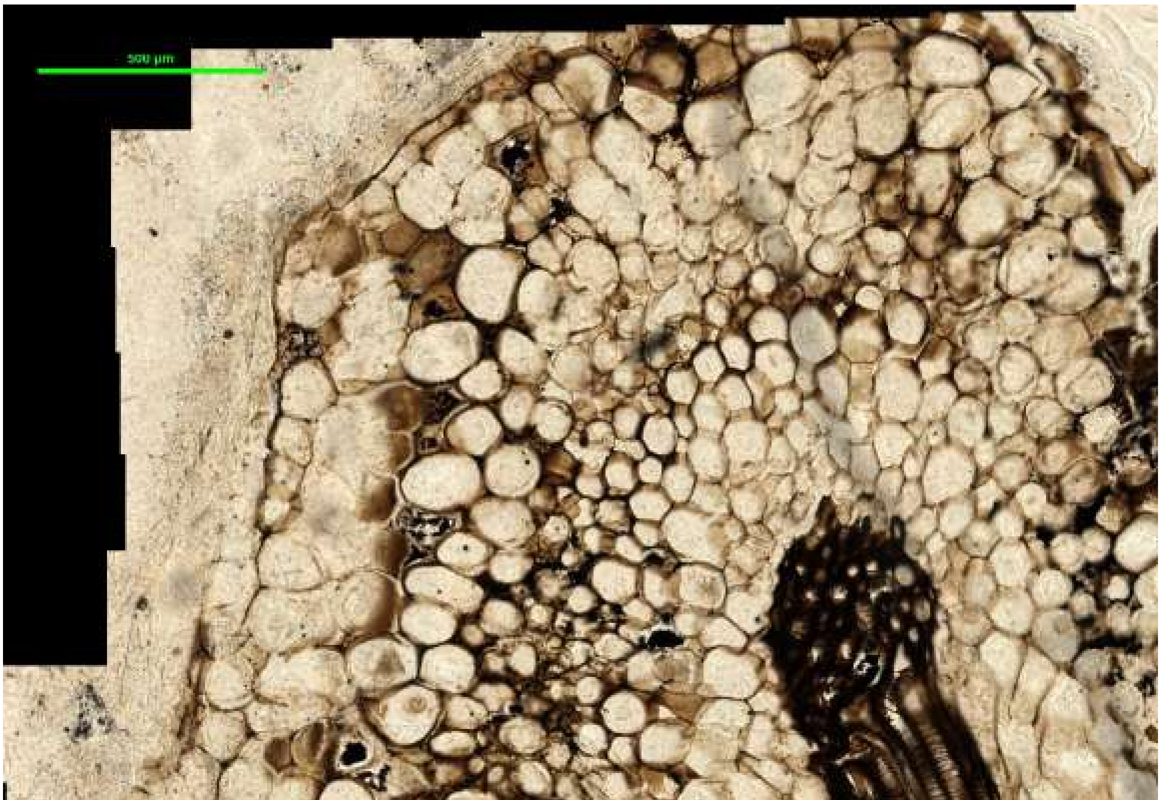


*Asteroxylon mackiei* rooting axis (STA 355.61): Aster_root_1

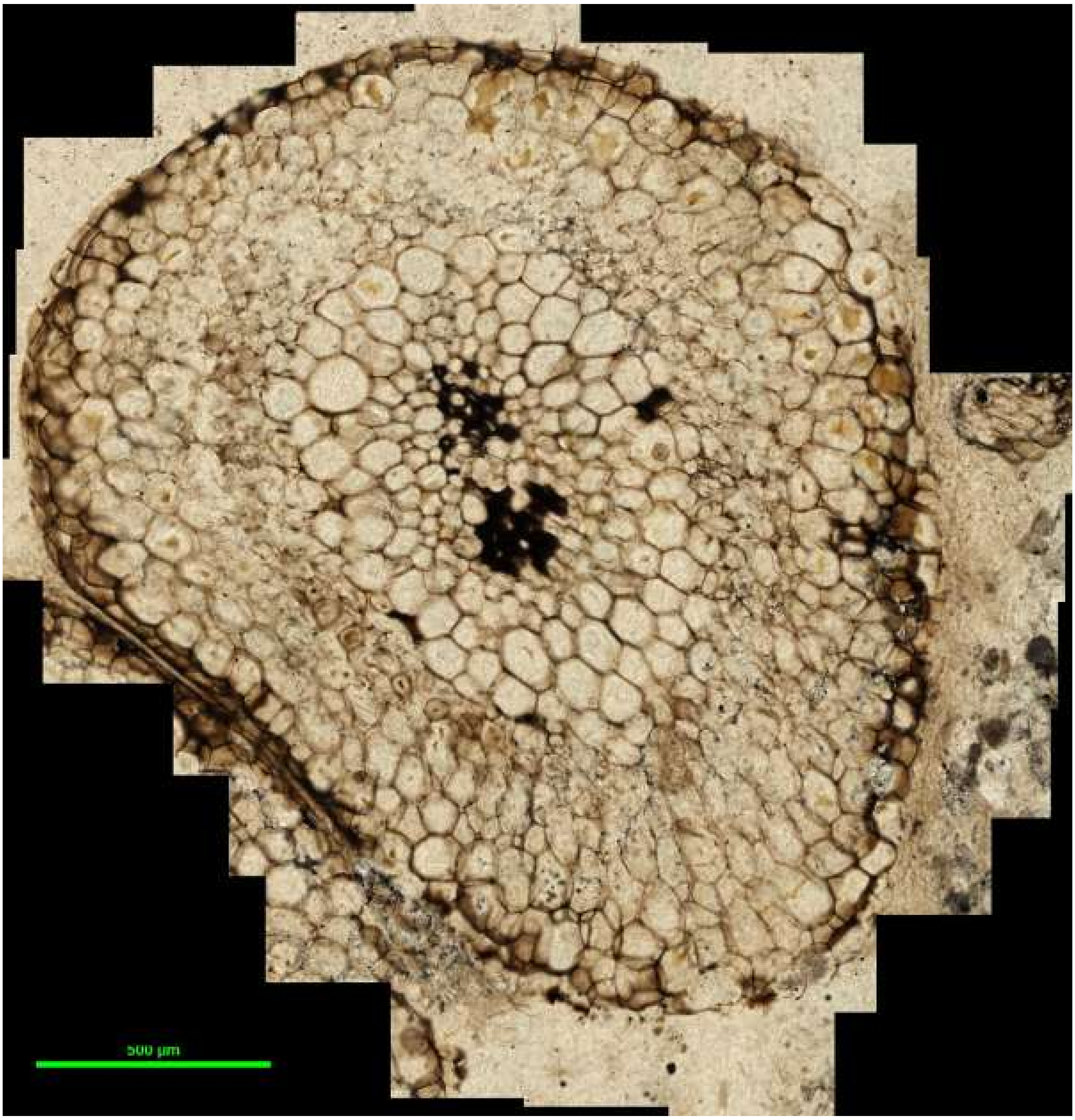


*Asteroxylon mackiei* rooting axis (R.1193): Aster_root_2

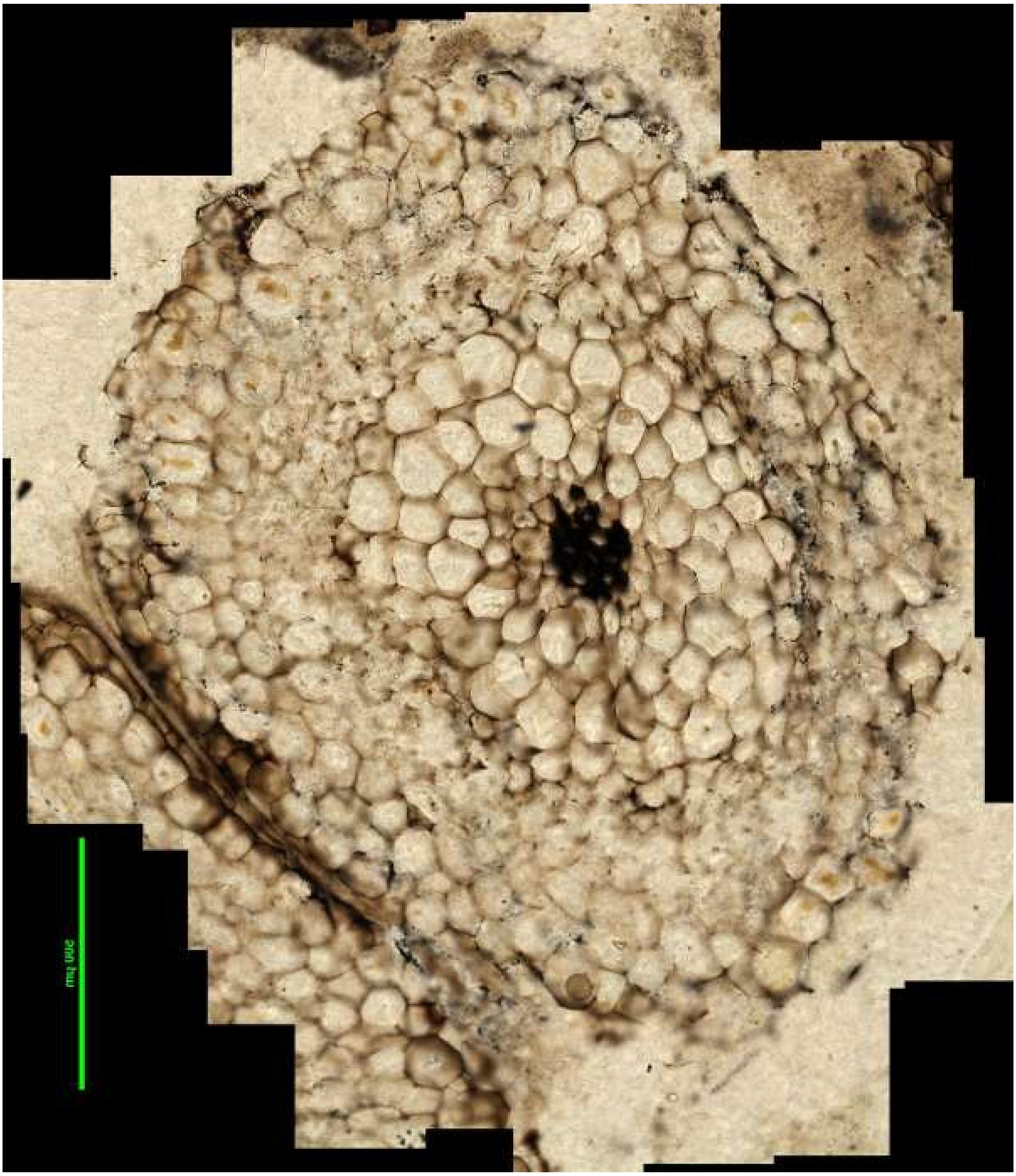

*Asteroxylon mackiei* rooting axis (R.1193): Aster_root_3

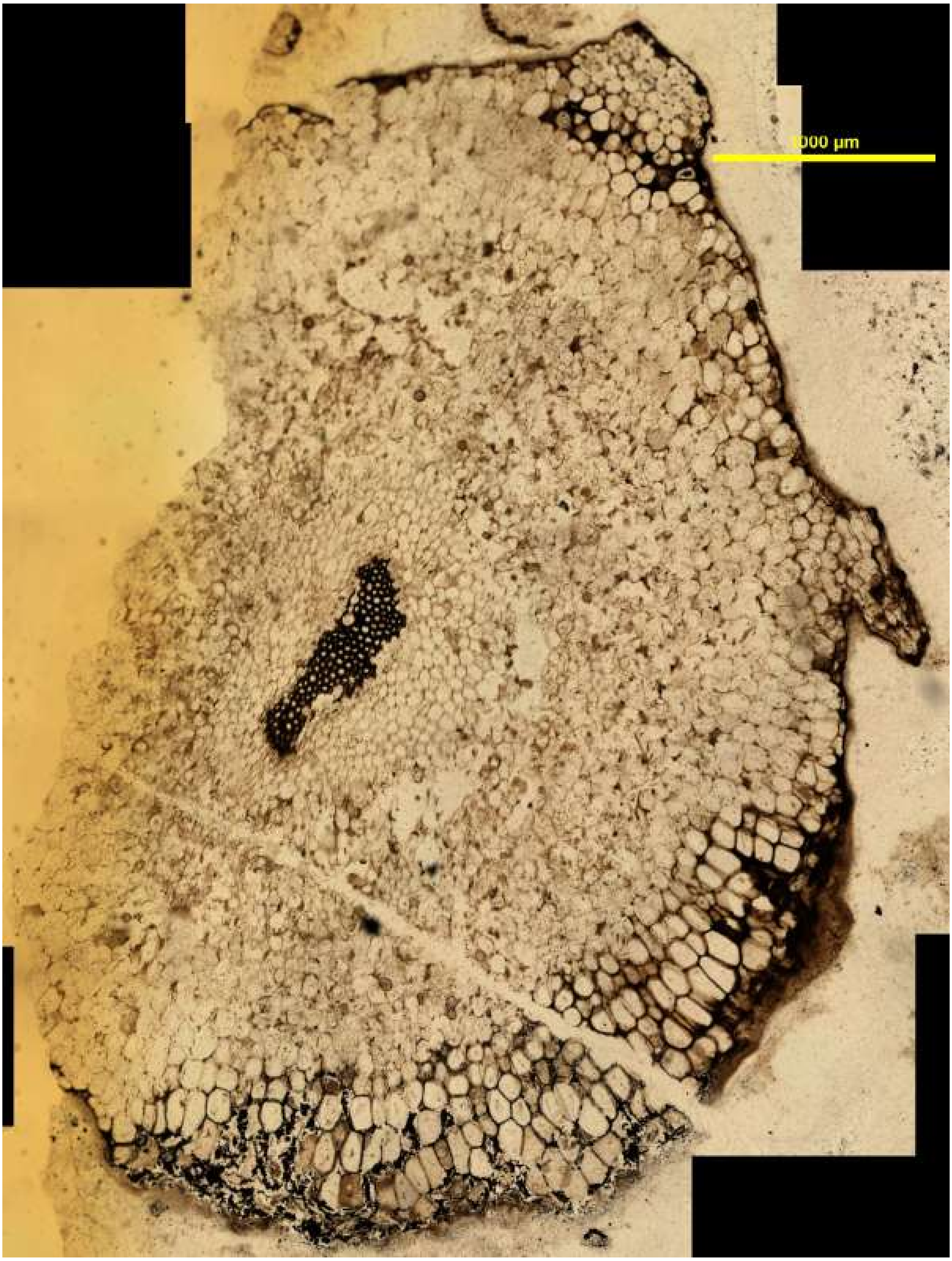


*Asteroxylon mackiei* root-bearing axis (OXF 423.70): Aster_root_bear_1

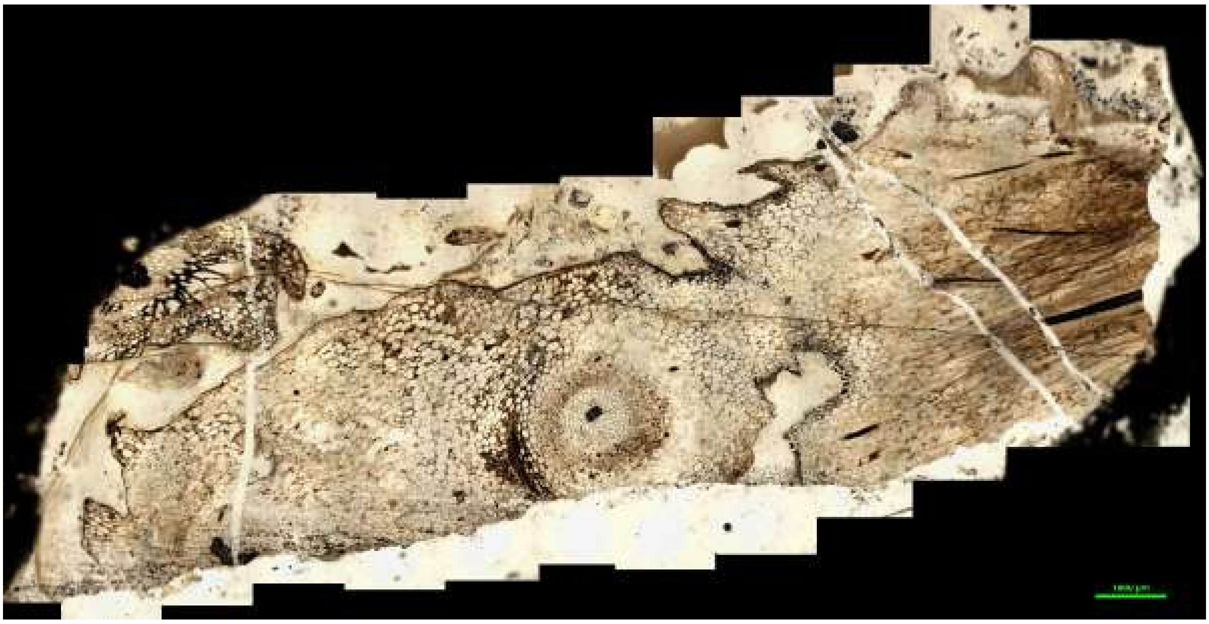

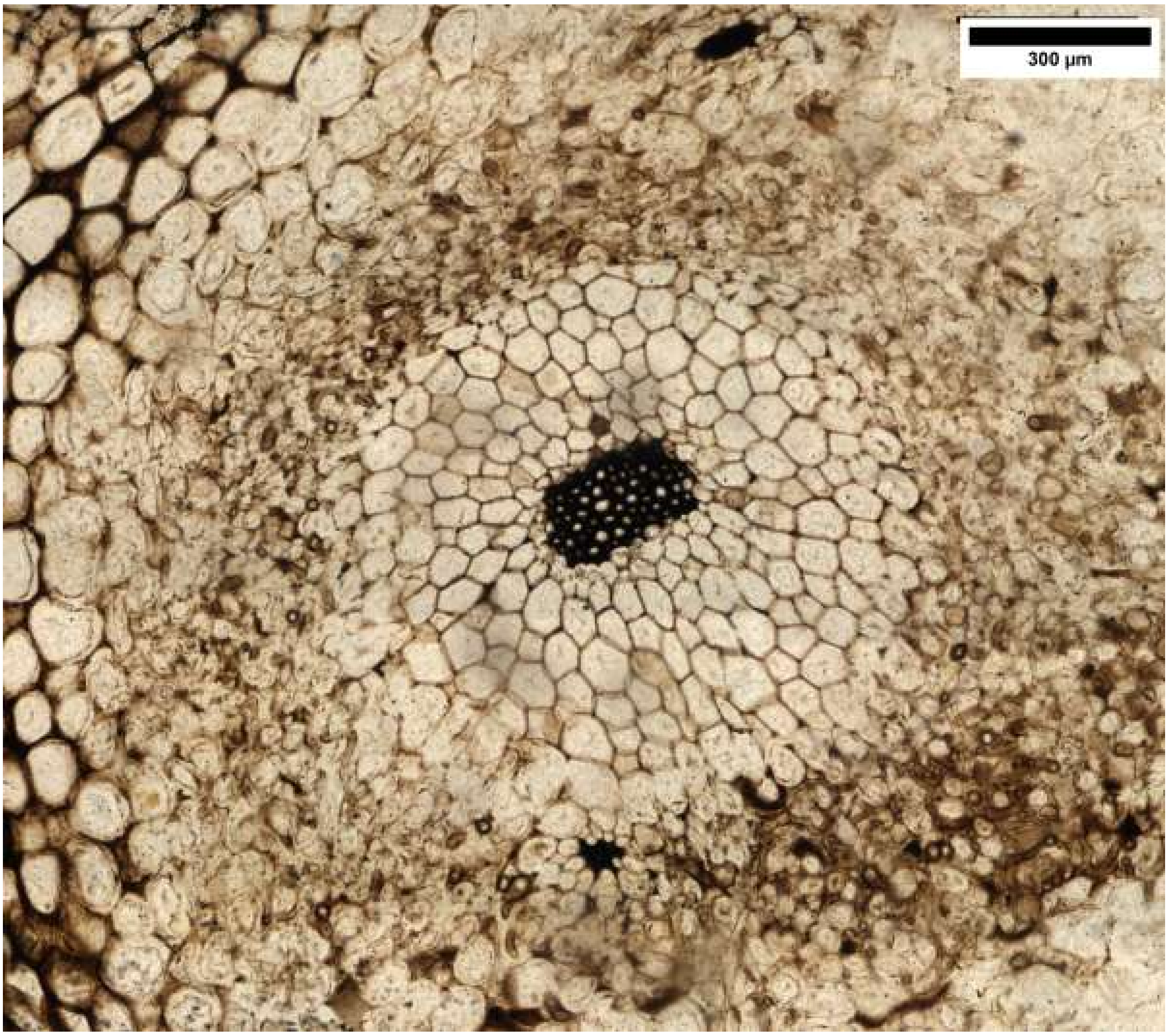


*Asteroxylon mackiei* root-bearing axis (R.1190): Aster_root_bear_2

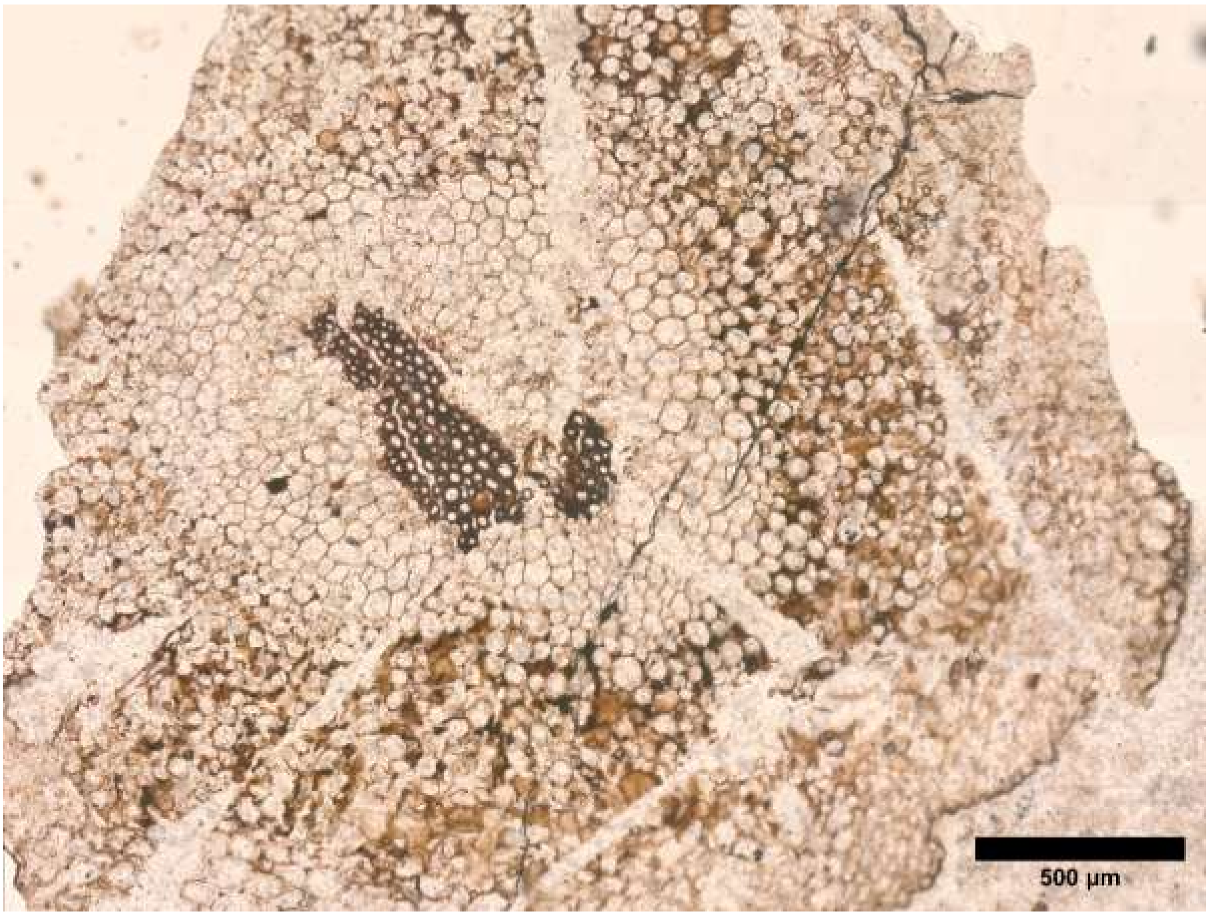


*Asteroxylon mackiei* root-bearing axis (GLAHM Kid 2472): Aster_root_bear_3

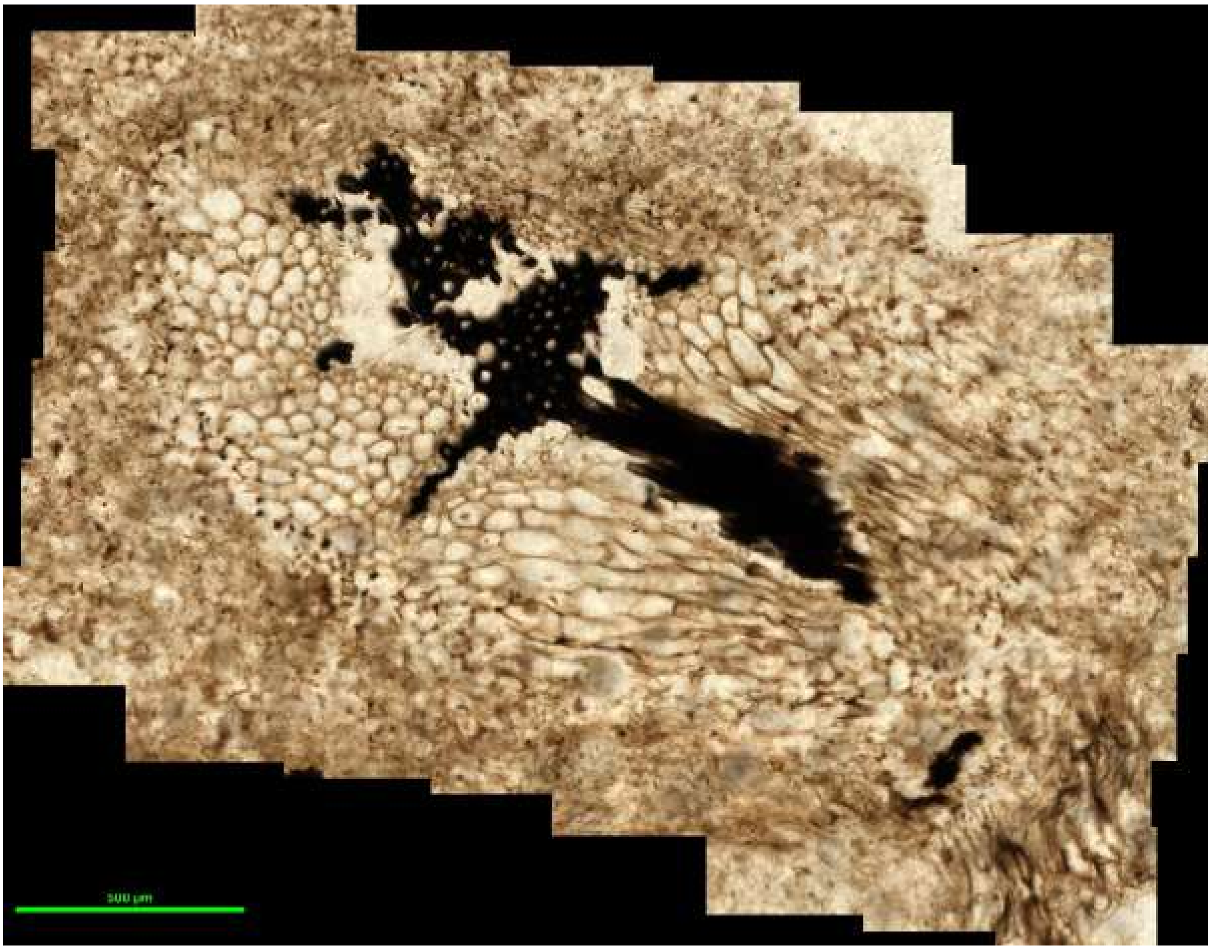

*Asteroxylon mackiei* leafy stem (NHMUK 16433): Aster_stem_1

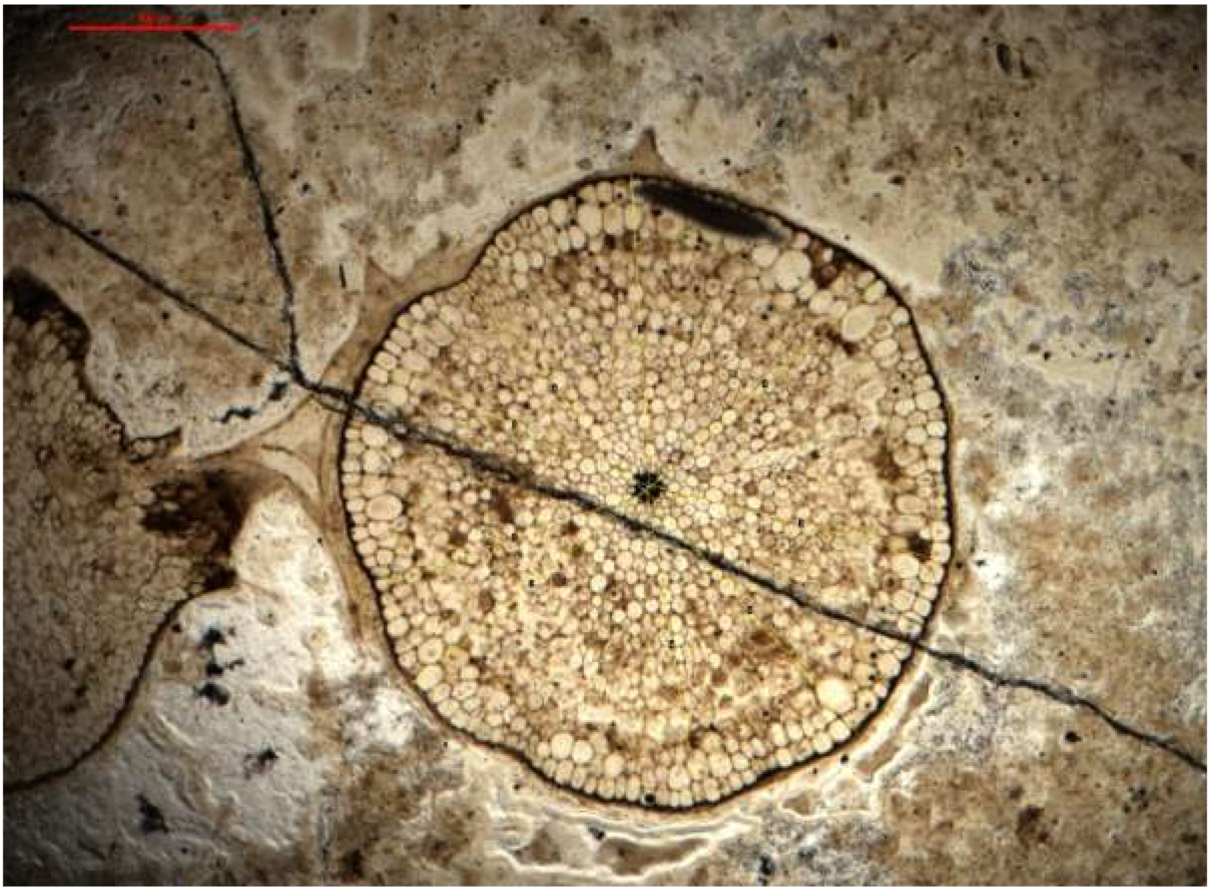

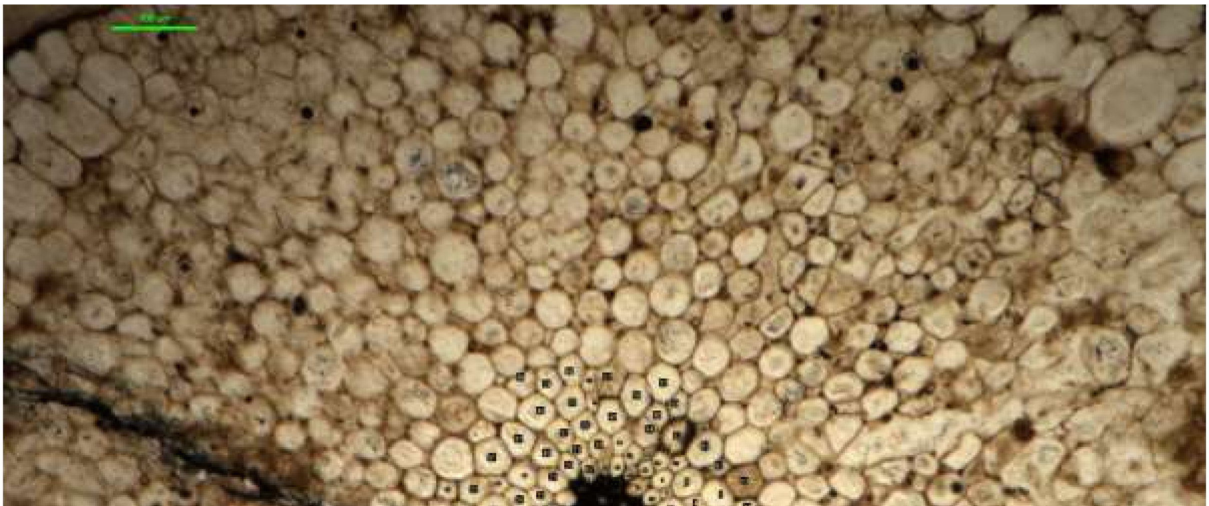

*Rhynia gwynne-vaughanii* (424.23): Rhynia_1

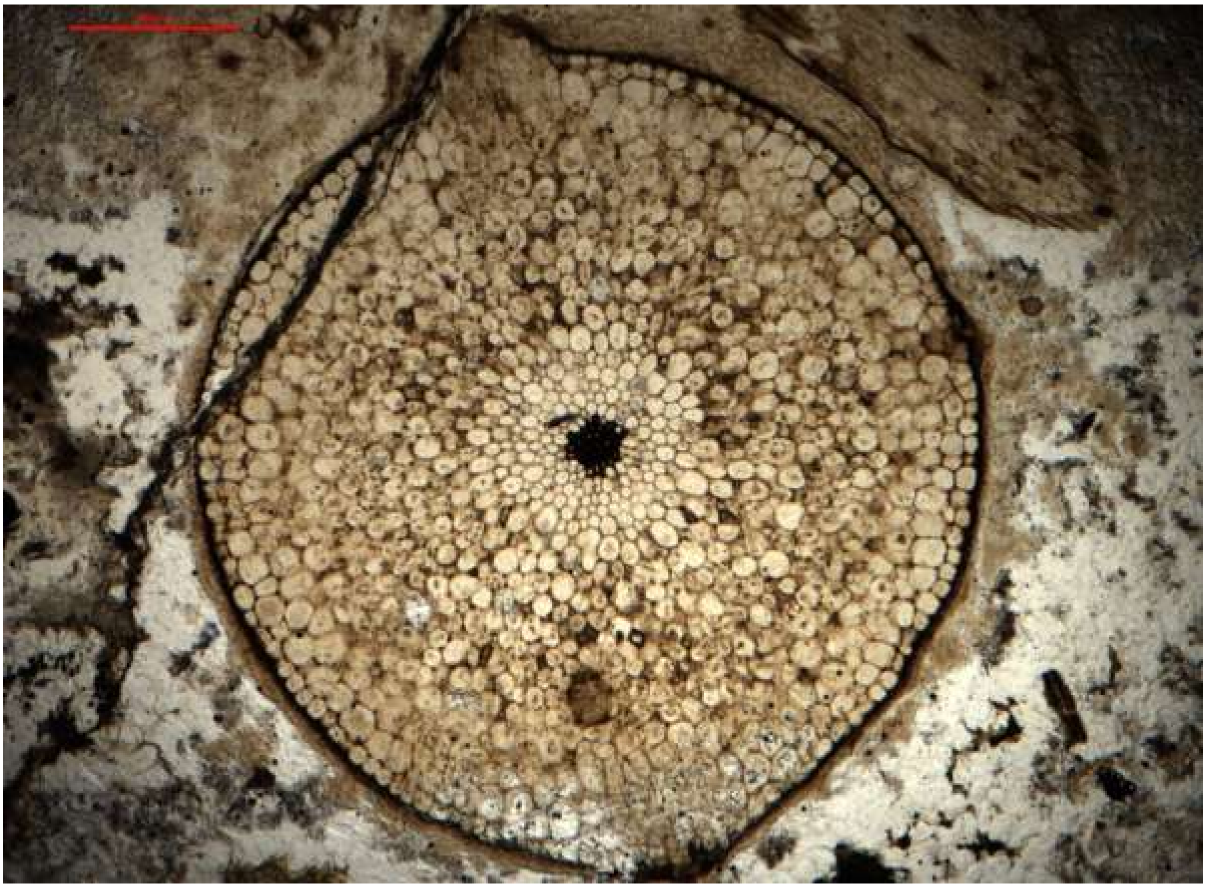

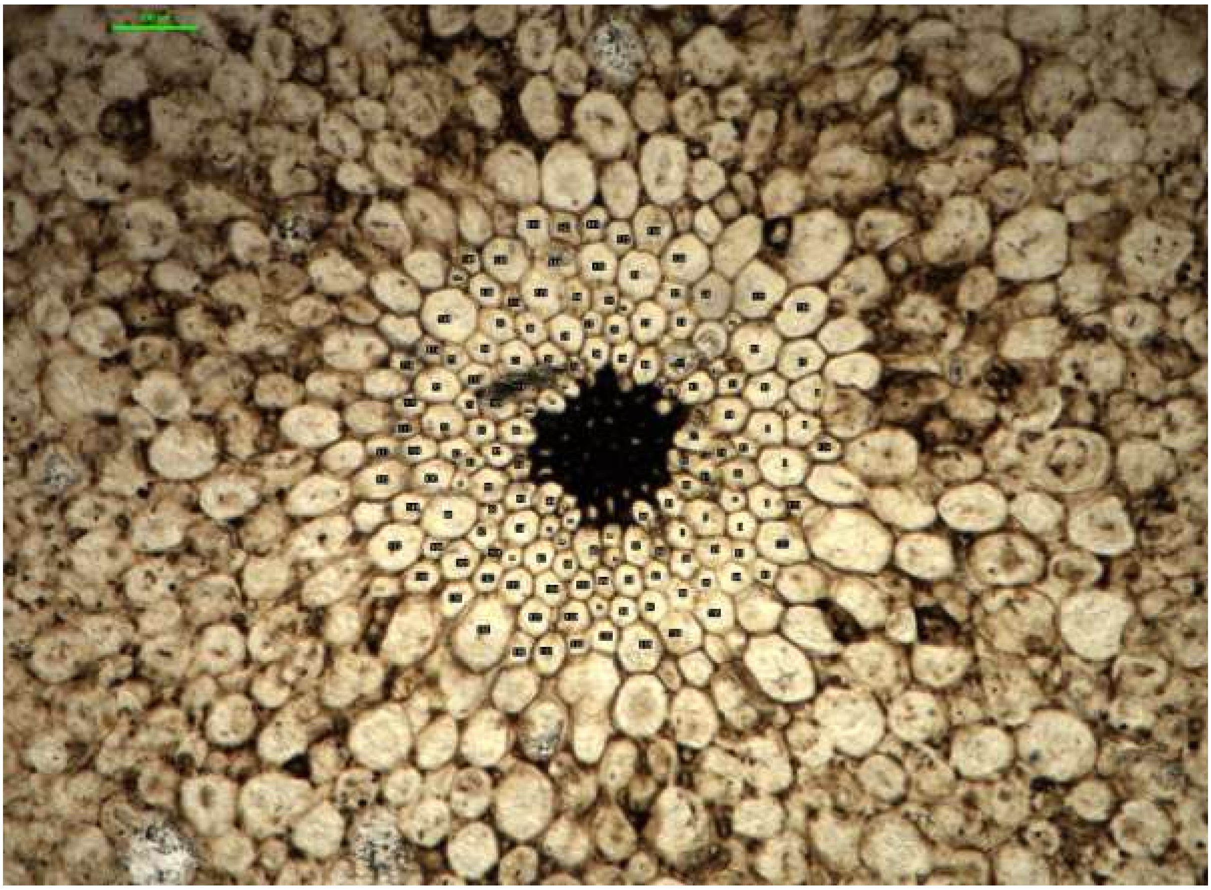

*Rhynia gwynne-vaughanii* (424.23): Rhynia_2

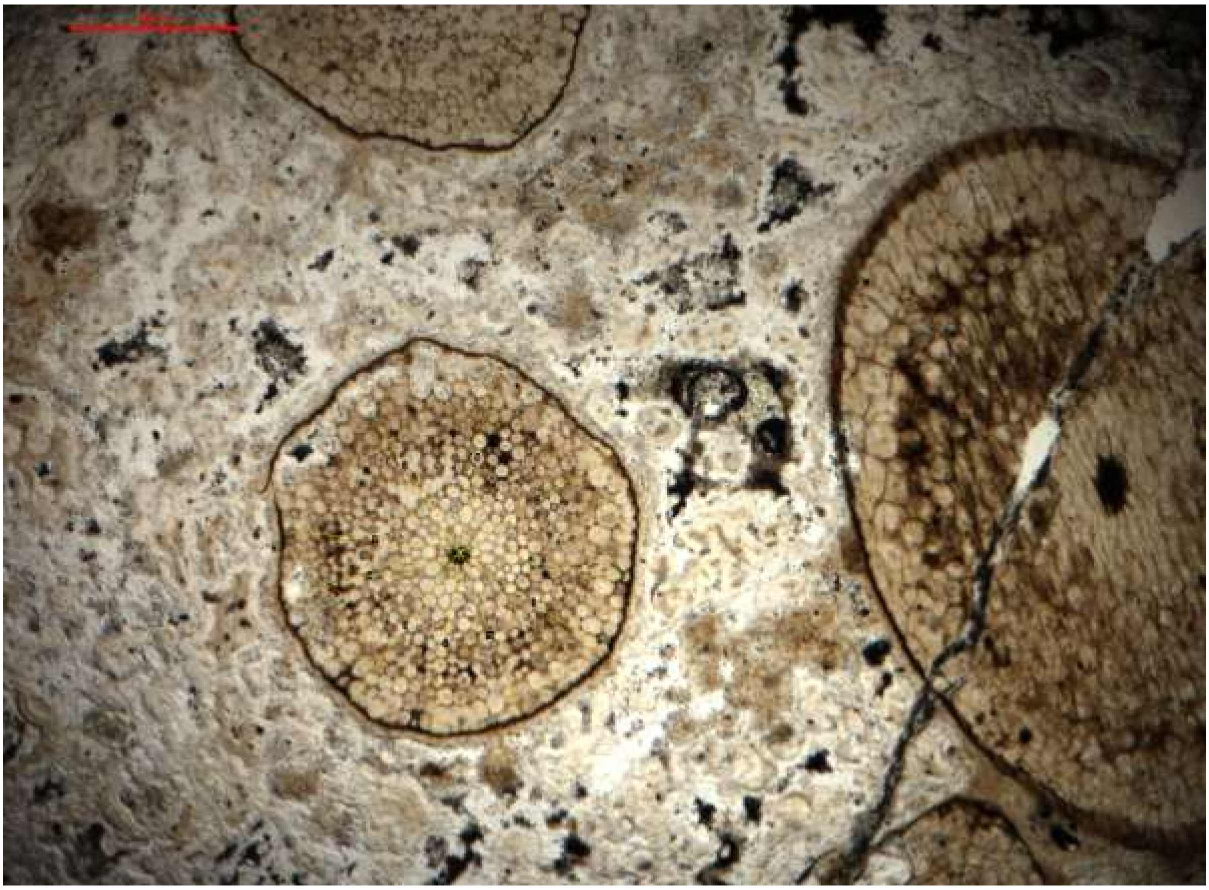

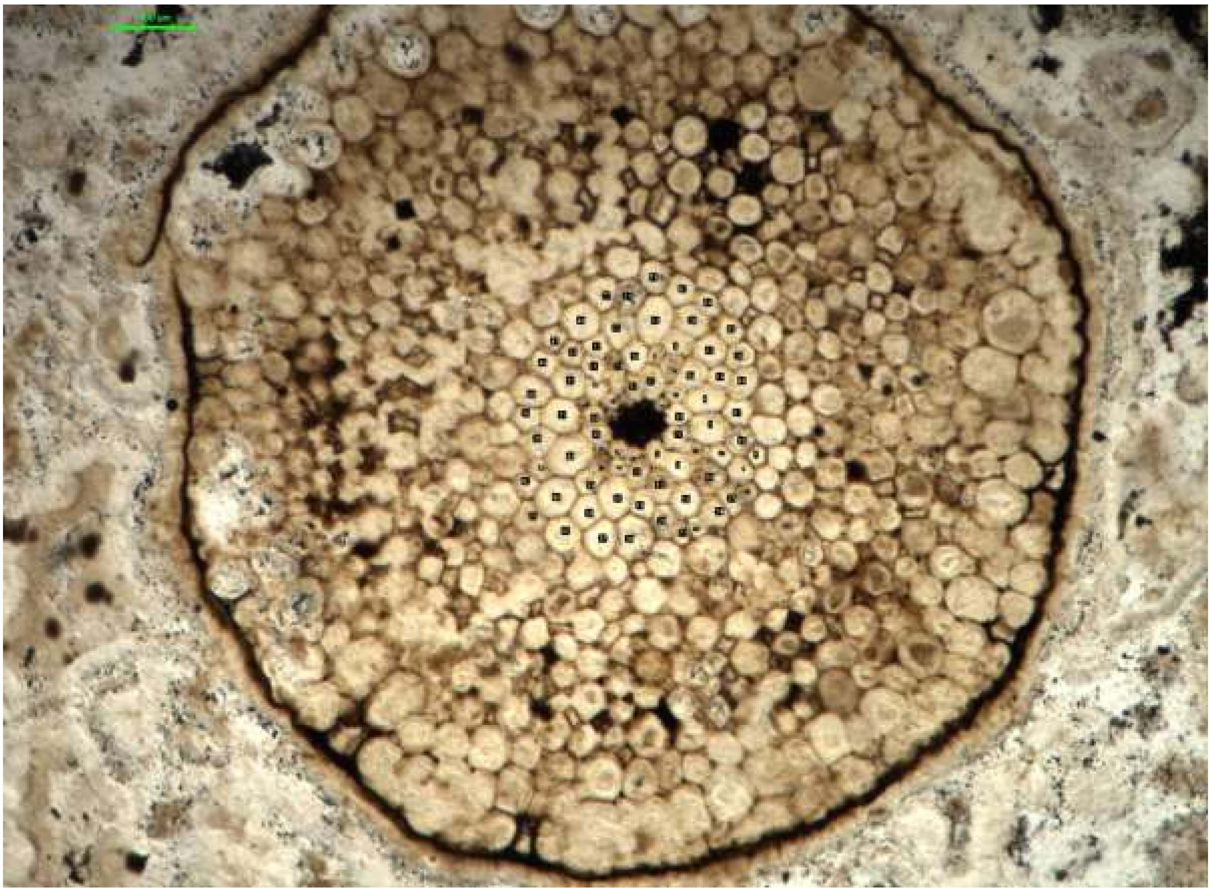

*Rhynia gwynne-vaughanii* (424.23): Rhynia_3

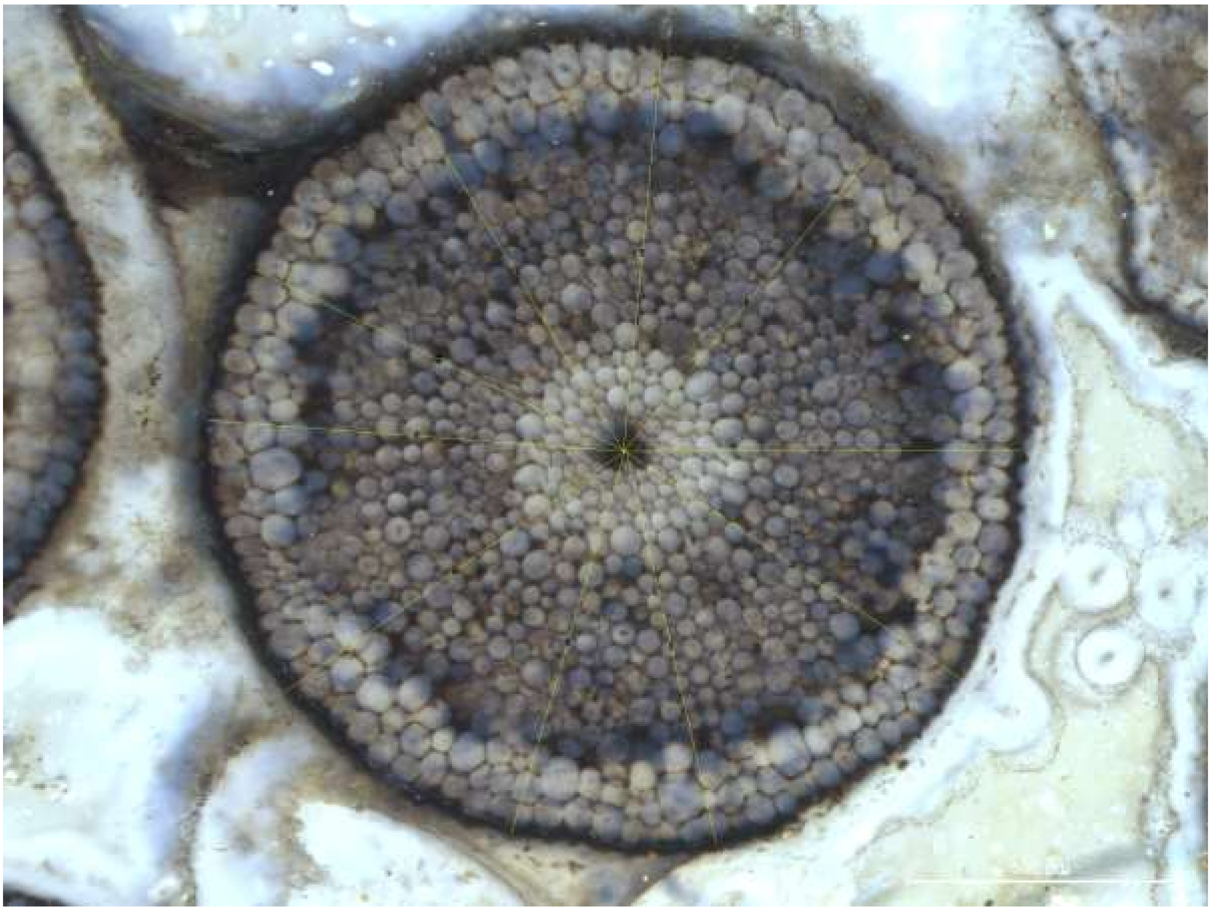

*Rhynia gwynne-vaughanii* (Pb 5007): Rhynia_4

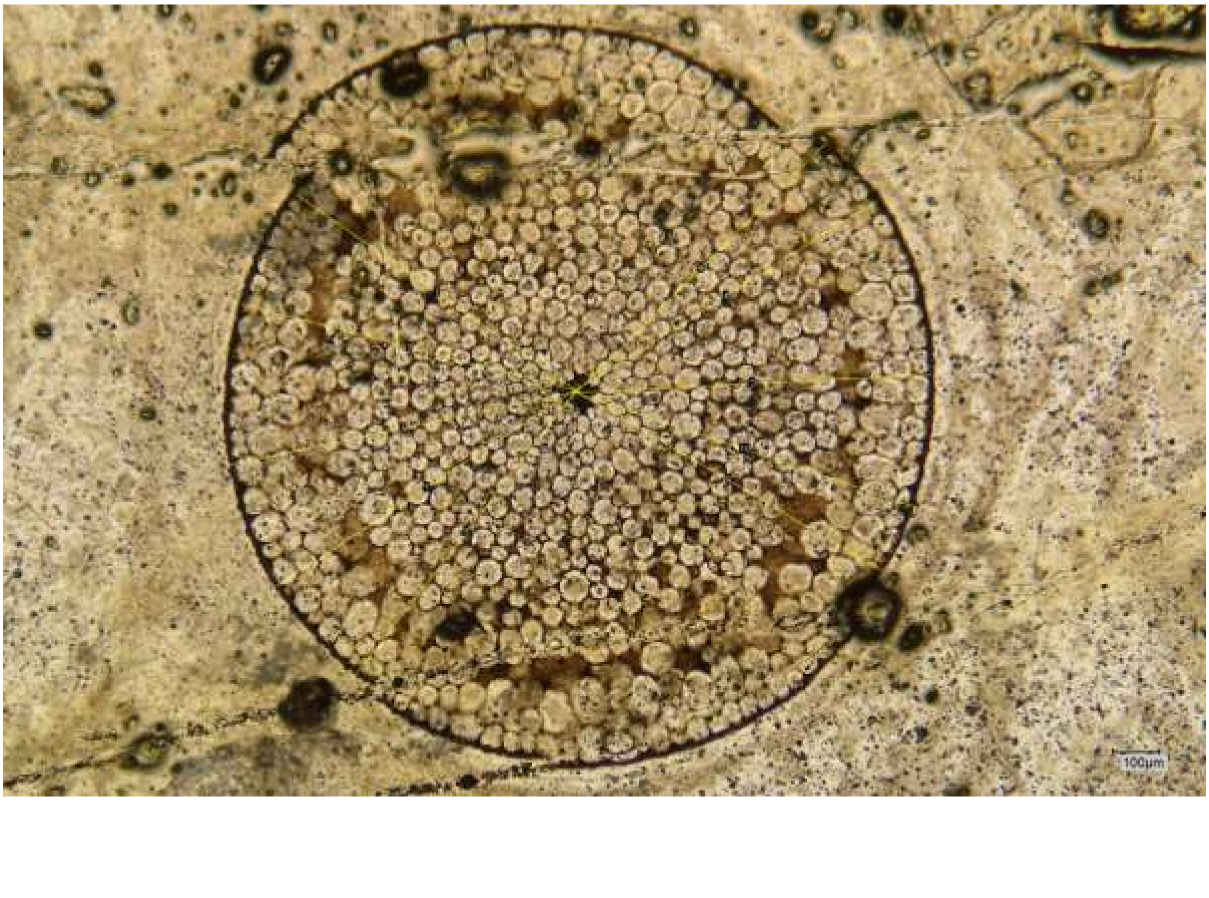

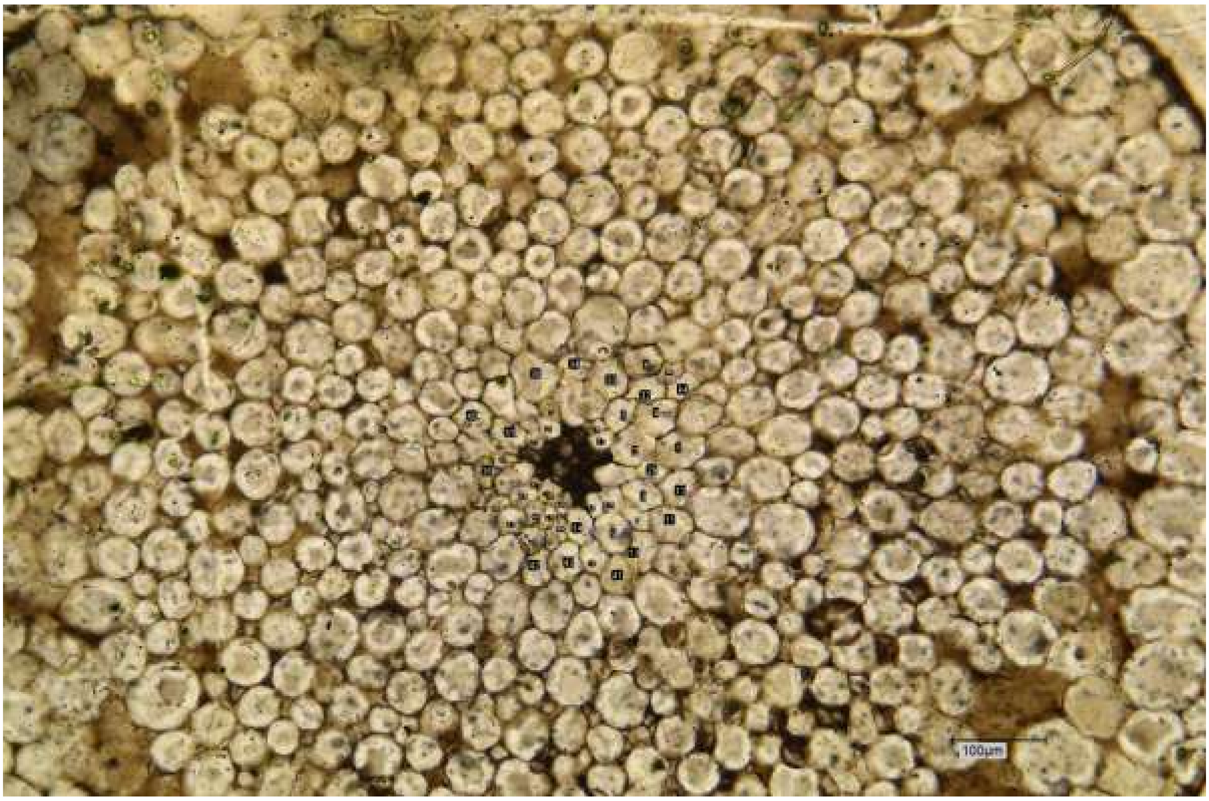


*Rhynia gwynne-vaughanii* (Pb 1588): Rhynia_5

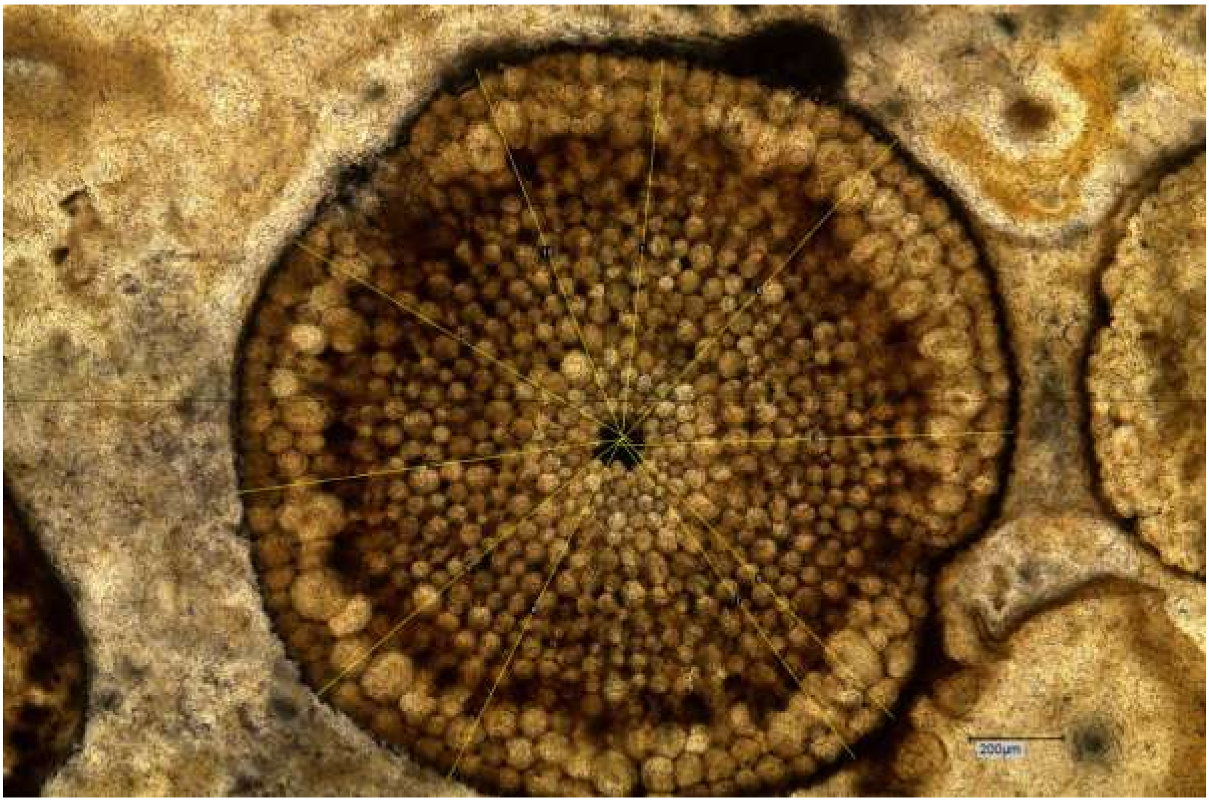

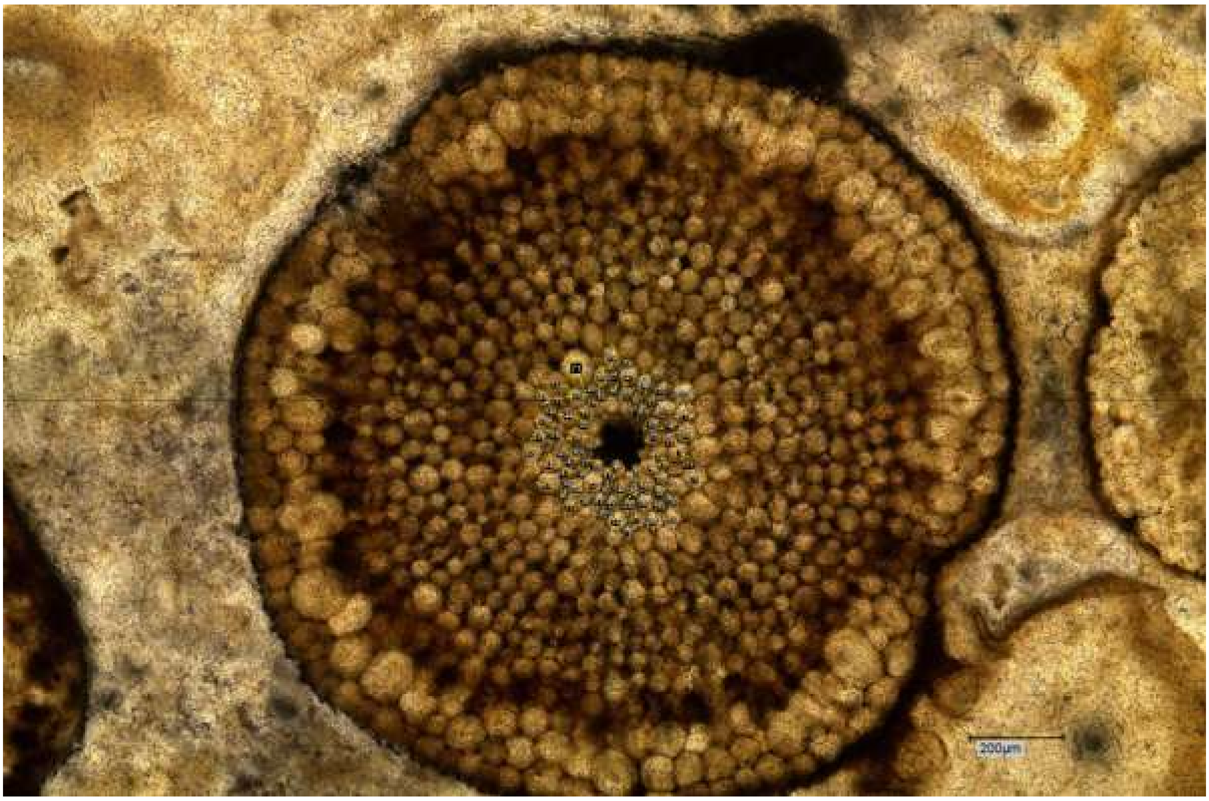


*Rhynia gwynne-vaughanii* (Pb 5006): Rhynia_6

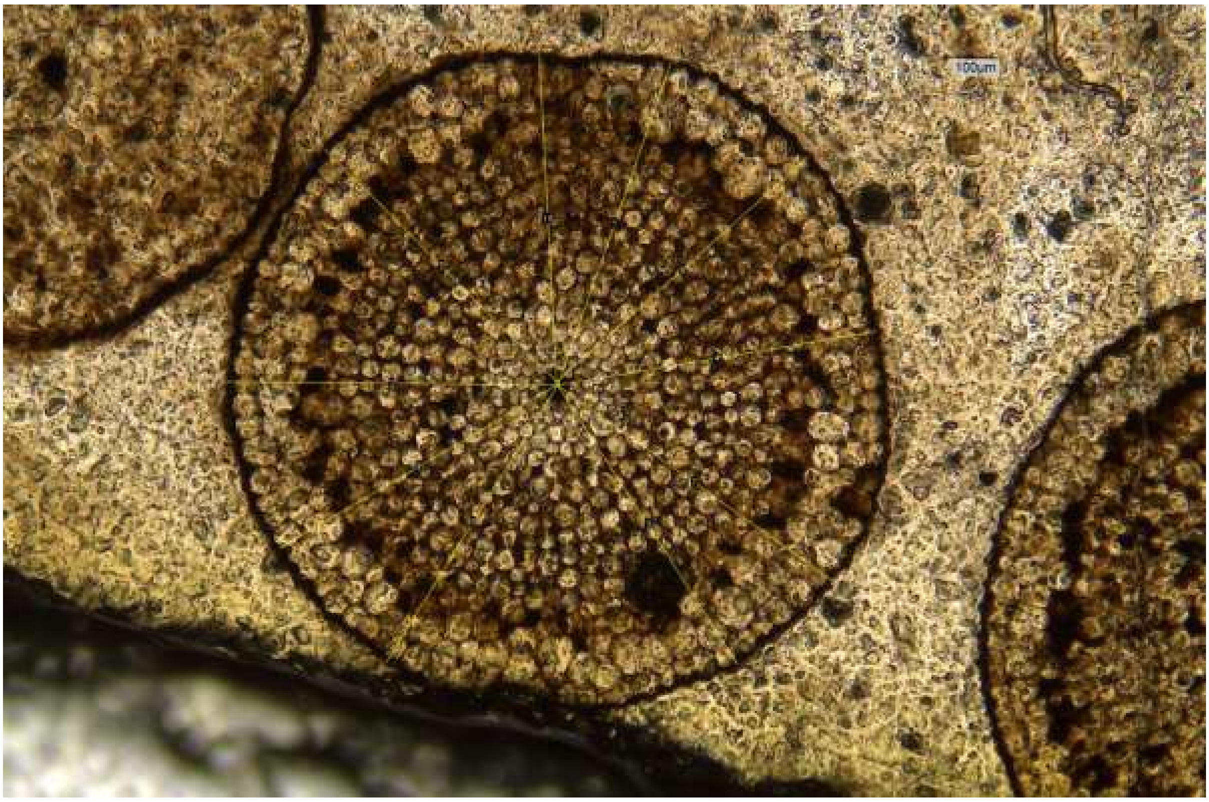

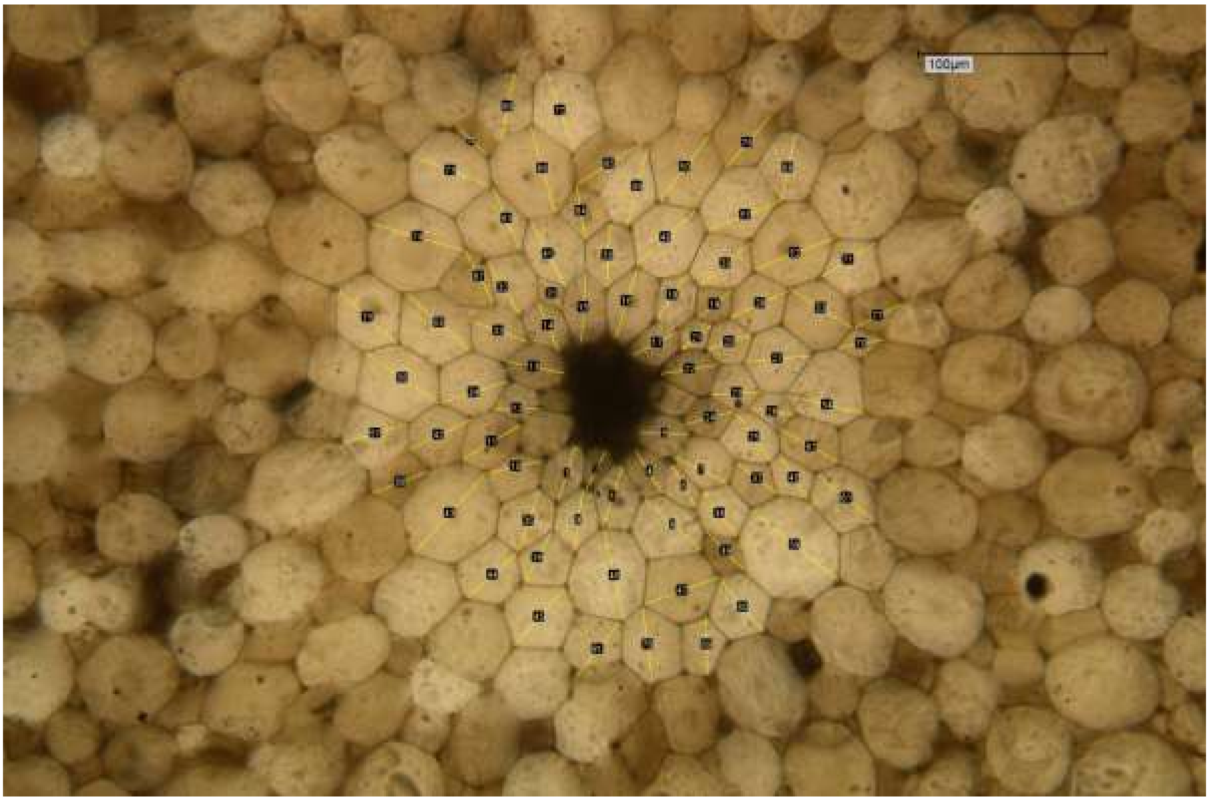


*Rhynia gwynne-vaughanii* (Pb 1809): Rhynia_7

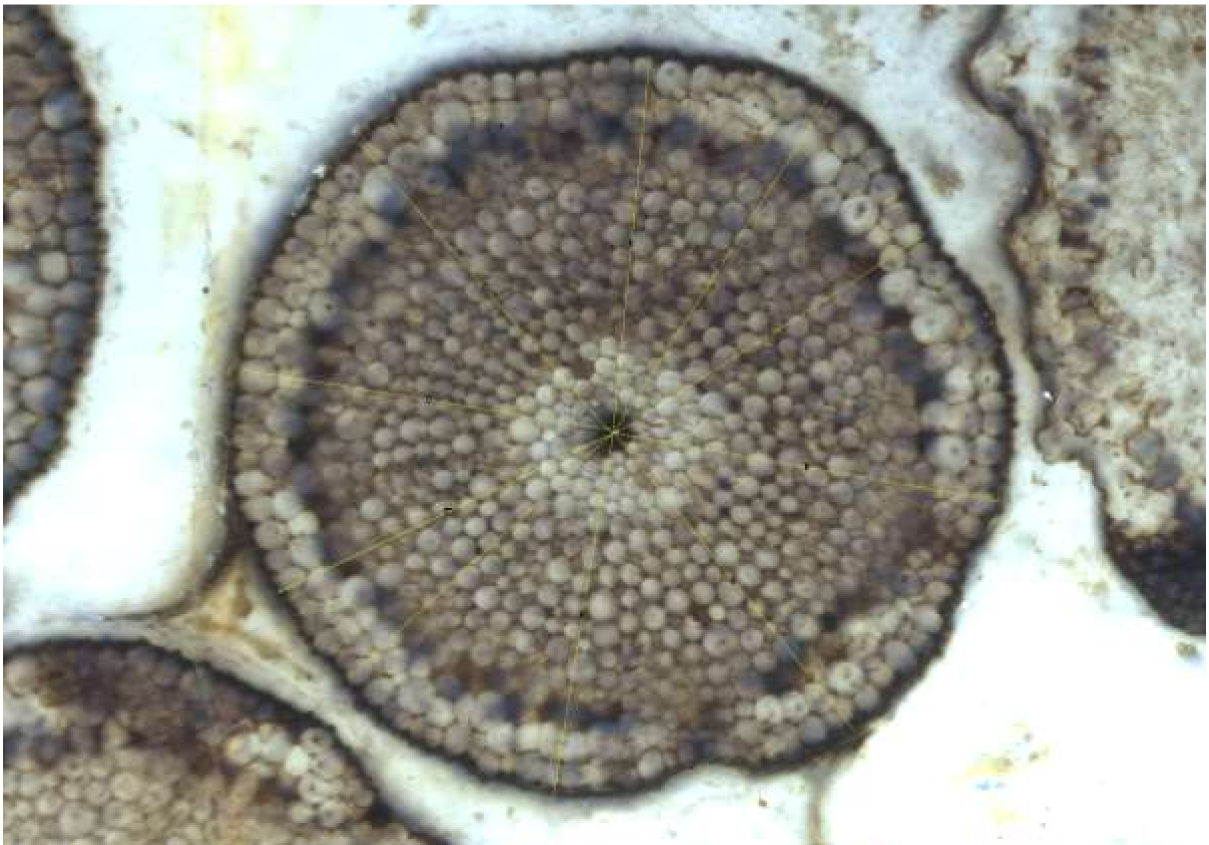

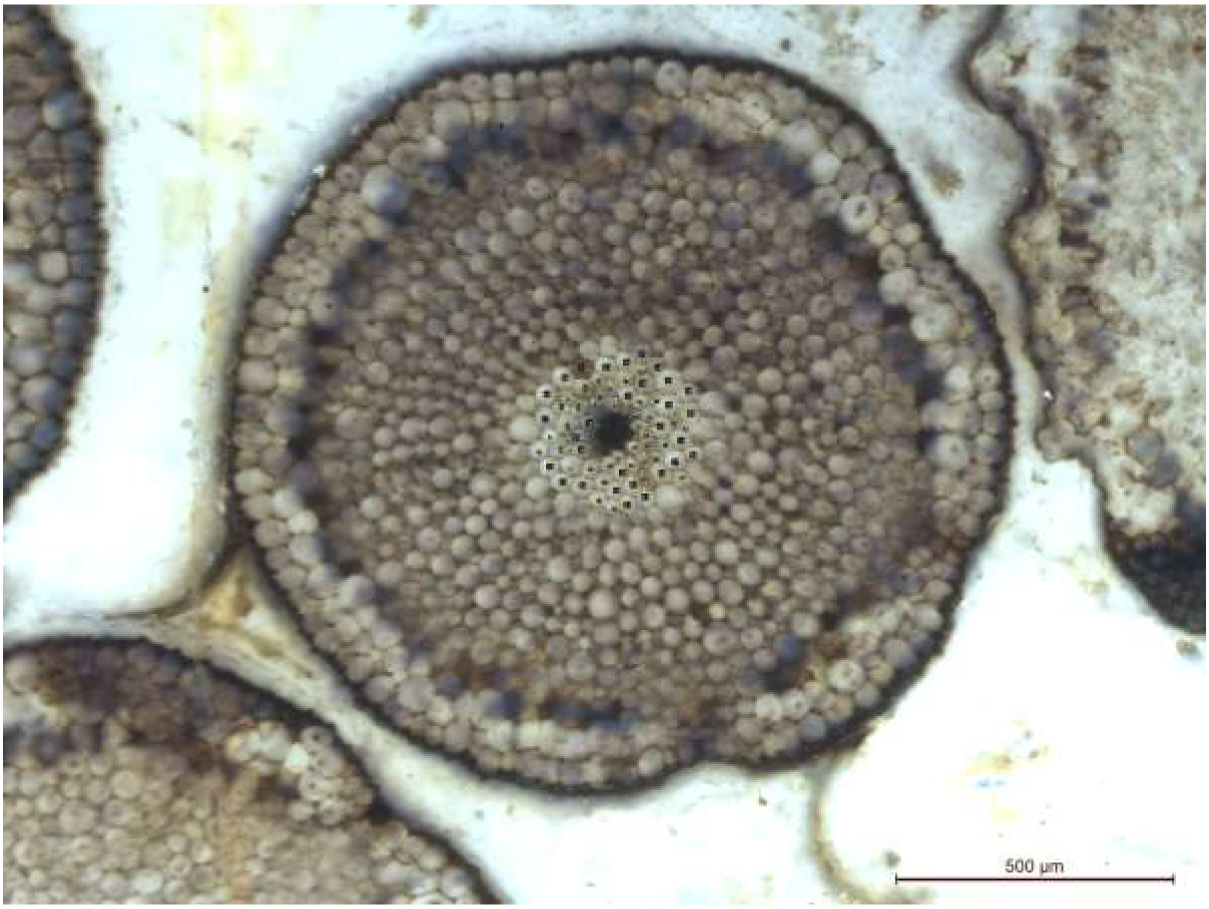


*Rhynia gwynne-vaughanii* (Pb 1876): Rhynia_8

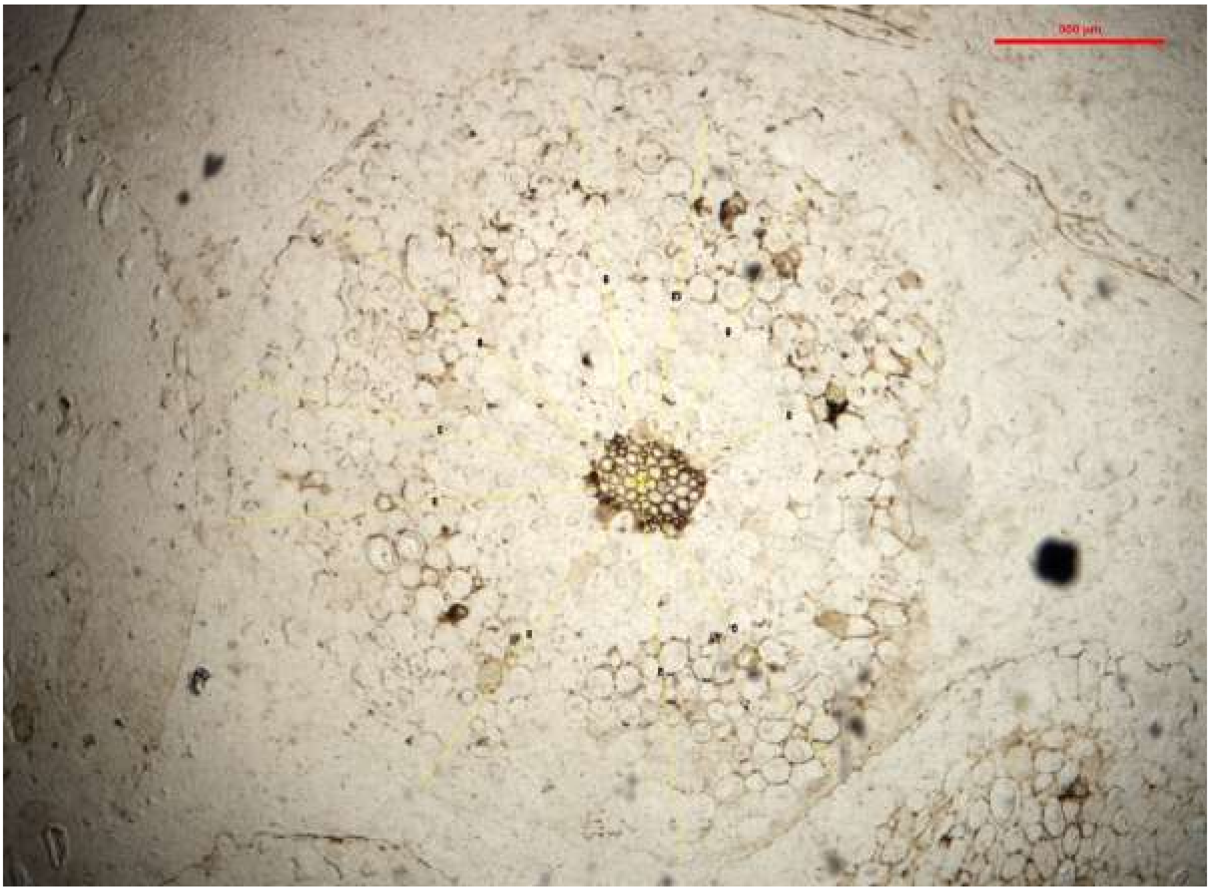

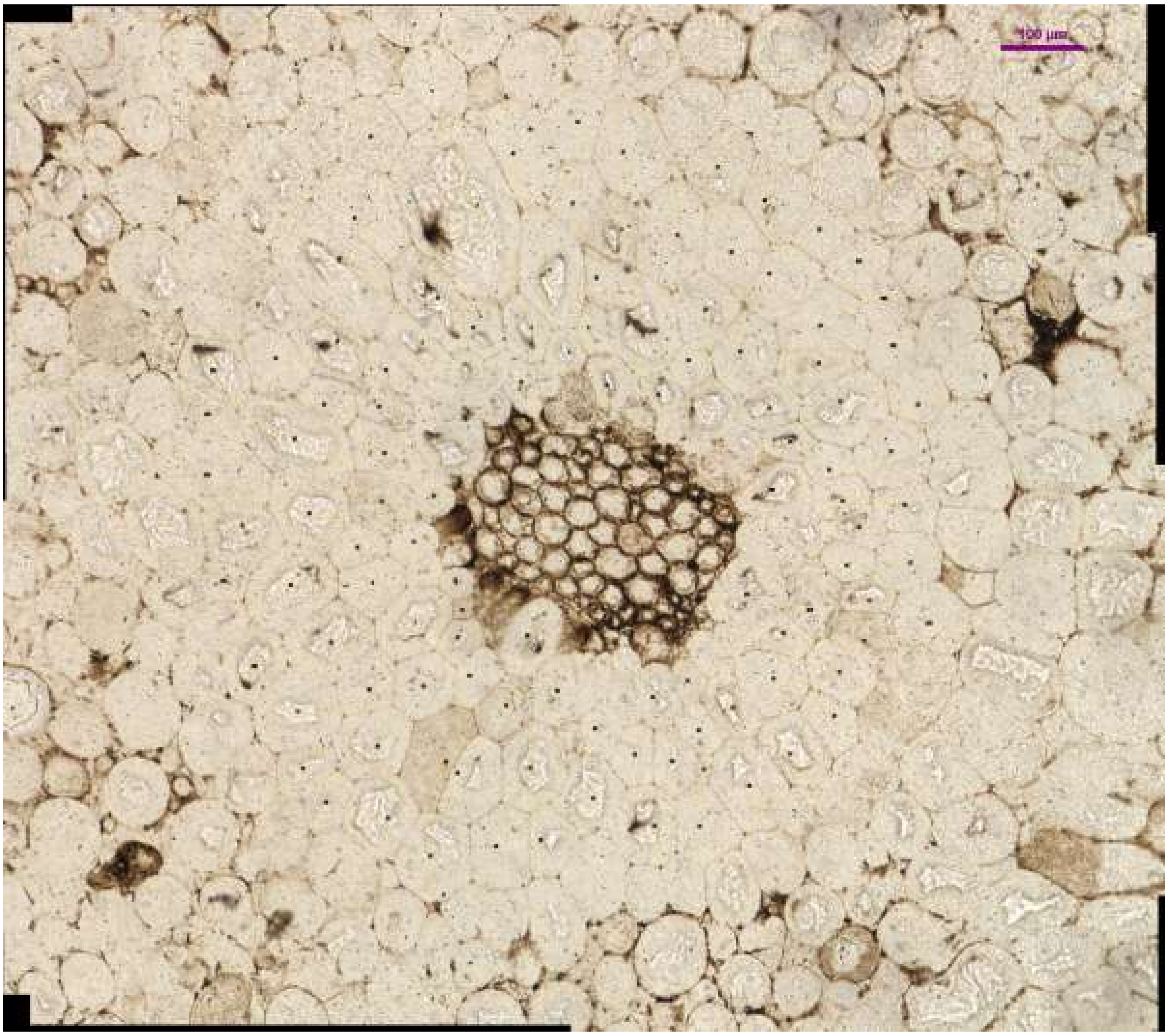


*Trichopherophyton teuchansii* (LYON 93.15): Trich_1

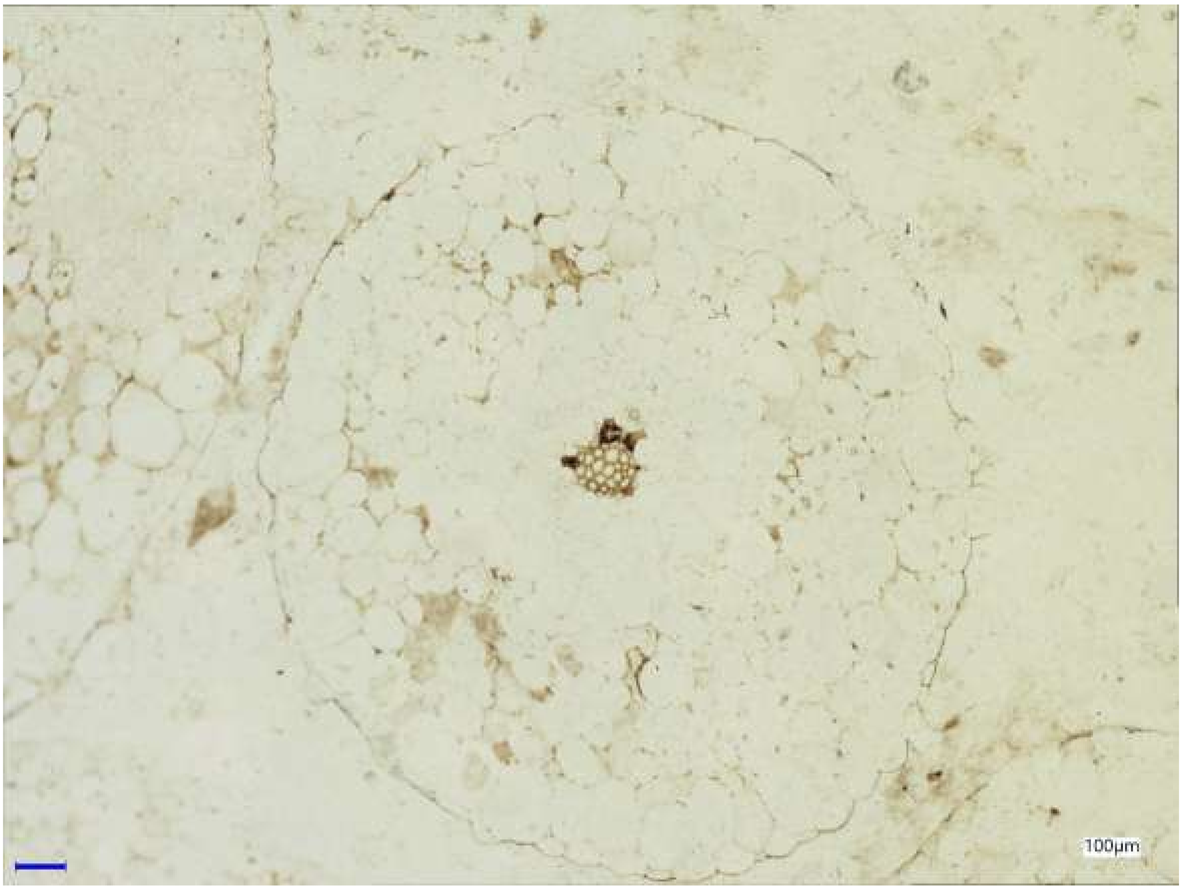


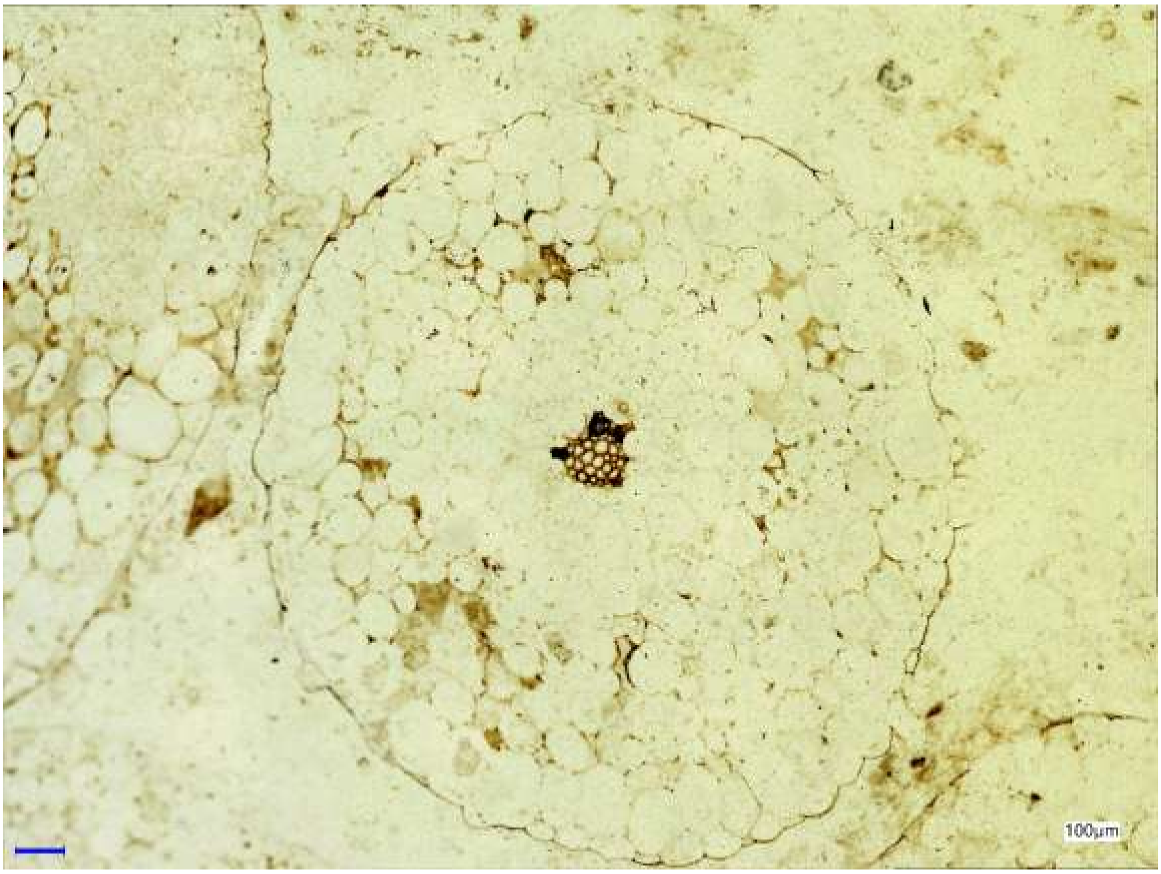


*Trichopherophyton teuchansii* (LYON 93.11): Trich_2
